## Supplementary Text for "Addressing age related hearing loss through engineering accessible and affordable hearing technology"

### Supplementary Information - The LoCHAid: An low-cost, open-source hearing aid for Age Related Hearing Loss

(Dated: September 30, 2019)

#### CONTENTS


I. Supplementary Movies 2
- A. S1: Construction of the LoCHAid 2
- B. S2: Preparing Earphones for Testing 2
- C. S3: Water Depth Testing of Device 2
- D. S4: Drop Testing Device 2

II. Schematic 2

III. ARHL Profiles 2

IV. Discussion on Equivalent Input Noise 2

V. Supplementary Master Graphs of Audiological Profiles and Targets (dB SPL) 3
- A. ISTS 55 dB SPL Master Graph 3
- B. ISTS 80 dB SPL Master Graph 3
- C. G.R.A.S KEMAR Left Ear ISTS 65 dB SPL Master Graph 3
- D. G.R.A.S KEMAR Right Ear ISTS 65 dB SPL Master Graph 3
- E. G.R.A.S KEMAR Right Ear Graph with ISTS 65 Speechmap Response (dB SPL) 3
- F. G.R.A.S KEMAR Left Ear Graph with ISTS 65 Speechmap Response (dB SPL) 3

VI. Quantification of Fits via Strict and Loose Criteria 3

VII. Supplementary Individual Graphs of Audiological Profiles and Targets (dB SPL) 3
- A. Male Age 60-69 Left Ear 3
  1. ISTS 55 dB SPL 3
  2. ISTS 65 dB SPL 3
  3. ISTS 80 dB SPL 3
  4. KEMAR Right Ear 65 dB SPL 3
  5. KEMAR Left Ear 65 dB SPL 3
- B. Male Age 60-69 Right Ear 4
  1. ISTS 55 dB SPL 4
  2. ISTS 65 dB SPL 4
  3. ISTS 80 dB SPL 4
  4. KEMAR Right Ear 65 dB SPL 4
  5. KEMAR Left Ear 65 dB SPL 4
- C. Male Age 70-79 Left Ear 4
  1. ISTS 55 dB SPL 4
  2. ISTS 65 dB SPL 4
  3. ISTS 80 dB SPL 4
  4. KEMAR Right Ear 65 dB SPL 4
  5. KEMAR Left Ear 65 dB SPL 4
- D. Male Age 70-79 Right Ear 4
  1. ISTS 55 dB SPL 4
  2. ISTS 65 dB SPL 4
  3. ISTS 80 dB SPL 4
  4. KEMAR Right Ear 65 dB SPL 4
  5. KEMAR Left Ear 65 dB SPL 4
- E. Female Age 60-69 Left Ear 4
  1. ISTS 55 dB SPL 4
  2. ISTS 65 dB SPL 4
  3. ISTS 80 dB SPL 4
  4. KEMAR Right Ear 65 dB SPL 4
  5. KEMAR Left Ear 65 dB SPL 4
- F. Female Age 60-69 Right Ear 5
  1. ISTS 55 dB SPL 5
  2. ISTS 65 dB SPL 5
  3. ISTS 80 dB SPL 5
  4. KEMAR Right Ear 65 dB SPL 5
  5. KEMAR Left Ear 65 dB SPL 5
- G. Female Age 70-79 Left Ear 5
  1. ISTS 55 dB SPL 5
  2. ISTS 65 dB SPL 5
  3. ISTS 80 dB SPL 5
  4. KEMAR Right Ear 65 dB SPL 5
  5. KEMAR Left Ear 65 dB SPL 5
- H. Female Age 70-79 Right Ear 5
  1. ISTS 55 dB SPL 5
  2. ISTS 65 dB SPL 5
  3. ISTS 80 dB SPL 5
  4. KEMAR Right Ear 65 dB SPL 5
  5. KEMAR Left Ear 65 dB SPL 5
- I. Hearing Profile X 5
  1. ISTS 55 dB SPL 5
  2. ISTS 65 dB SPL 5
  3. ISTS 80 dB SPL 5
  4. KEMAR Right Ear 65 dB SPL 5
  5. KEMAR Left Ear 65 dB SPL 5
- J. Hearing Profile Y 6
  1. ISTS 55 dB SPL 6

2. ISTS 65 dB SPL 6
3. ISTS 80 dB SPL 6
4. KEMAR Right Ear 65 dB SPL 6
5. KEMAR Left Ear 65 dB SPL 6
K. Hearing Profile Y 6
1. ISTS 55 dB SPL 6
2. ISTS 65 dB SPL 6
3. ISTS 80 dB SPL 6
4. KEMAR Right Ear 65 dB SPL 6
5. KEMAR Left Ear 65 dB SPL 6
L. Hearing Profile Z 6
1. ISTS 55 dB SPL 6
2. ISTS 65 dB SPL 6
3. ISTS 80 dB SPL 6
4. KEMAR Right Ear 65 dB SPL 6
5. KEMAR Left Ear 65 dB SPL 6

References 7

#### I. SUPPLEMENTARY MOVIES

##### A. S1: Construction of the LoCHAid

Video outlining construction of the LoCHAid with Lithium Ion Coin Cell Battery. Video speed has been increased to 15x; however, the average time of construction is 25 minutes. Figure S6 shows the schematic of the hearing aid.

##### B. S2: Preparing Earphones for Testing

Video outlining how to properly set up device earbuds for audiological testing in AudioScan Verifit with 0.2-cc coupler.

##### C. S3: Water Depth Testing of Device

Video showing device after being submerged in 6cm of water, and shows that it still is in working condition. A still photo is shown in Figure S5b.

##### D. S4: Drop Testing Device

Video showing device after repeated drop tested (n=10) from a height of 5 feet, and showing that the device is in working condition. A still photo is shown in Figure S5a.

#### II. SCHEMATIC

The schematic of the LoCHAid is given in Figure S6. The .brd files can be found in the github folder associated with this manuscript, which can be uploaded to fab-house website such as OSHPARK (www.oshpark.com) for PCB fabrication.

#### III. ARHL PROFILES

Figure S1 shows the audiograms used for analysis and comparison. They contain both male and female aged 60-69, 70-79 for both left and right ears. The male audiograms are steeper and have sharper fall offs, as compared to female audiograms. The data comes from two studies. Cruikshanks, et al from 1993-1995 [1] conducted hearing tests of both ears of people living in Beaver Dam, Wisconsion, and Ciletti and Flamme [2] who conducted hearing loss analysis cross country in USA. Cruikshanks, et al from 1993-1995 conducted hearing tests of both ears of people living in Beaver Dam, Wisconsion, and pulls data from 566 females from the age of 60-69, 534 females from ages 70-79, 489 males from ages 60-69, and 355 males from the ages of 70-79 and detail left and right ear hearing losses. Hearing Profiles X,Y, and Z are gender neutral audiograms displaying ARHL characteristics pulled from a study done by Ciletti and Flamme, which was conducted over 1999-2005, based on the National Health and Nutrition Examination Survey (NHANES) which was a cross-country sample study of common audiogram trends, based on 2819 women, and 2525 men ages 20-69. These results are in agreement with other work [3, 4].

#### IV. DISCUSSION ON EQUIVALENT INPUT NOISE

EIN at 40 dB SPL can be problematic as many people with mild hearing loss at a threshold of 0-20 dB HL are likely to hear this sound and this internal noise can interfere with their speech understanding. However, for the majority of the hearing loss profiles investigated the average threshold is 40 dB HL [1, 2]. A study conducted by Macae and Dillon in 1996 specified a relaxation criteria up to 46.3 dB SPL for a gain of 15 dB at a threshold of 20-50 dB HL. Based on this, we think that the EIN will interfere very little for patients exhibiting mild to moderate ARHL [5].

We were able to track the source of the EIN In the circuit to the MAX9814 module itself at 30 nV/$\sqrt{Hz}$ [6], and to the best of our knowledge, no filter was able to drop the EIN to below 40 dB SPL. We undertook experimentation with different Maxim models such as MAX4466, Silicon MEMS microphone SPW2430 (www.adafruit.com). We undertook different model configurations with using op-amp TLV2462 in replacing the MAX98306, and found out that it did not only did it not reduce the EIN, but it also impacted the frequency response. Overall, we tested 12 different configurations with different components, both passive and active filter circuits, to reduce EIN, but we were not successful in bringing down the EIN at the cost boundary of 1 USD. Hence, a decision was made not to address this feature at this stage as the electroacoustic analysis showed that the LoCHAid has high frequency gain that is necessary for mild-moderate ARHL.

#### V. SUPPLEMENTARY MASTER GRAPHS OF AUDIOLOGICAL PROFILES AND TARGETS (DB SPL)

The following figures detail all audiological profiles and their respective Targets (dB SPL) for each differing input ISTS 55 dB SPL,65 dB SPL, 80 dB SPL, and G.R.A.S KEMAR Left and Right Ears at ISTS 65 dB SPL

##### A. ISTS 55 dB SPL Master Graph

The graph is shown in figure number S7.

##### B. ISTS 80 dB SPL Master Graph

The graph is shown in figure number S8

##### C. G.R.A.S KEMAR Left Ear ISTS 65 dB SPL Master Graph

The graph is shown in figure number S10.

##### D. G.R.A.S KEMAR Right Ear ISTS 65 dB SPL Master Graph

The graph is shown in figure number S9.

##### E. G.R.A.S KEMAR Right Ear Graph with ISTS 65 Speechmap Response (dB SPL)

The graph is shown in figure number S11.

##### F. G.R.A.S KEMAR Left Ear Graph with ISTS 65 Speechmap Response (dB SPL)

The graph is shown in figure number S12.

#### VI. QUANTIFICATION OF FITS VIA STRICT AND LOOSE CRITERIA

We present a visual graph indicating how well LoCHAid fits the profiles in Figure S2. The blue squares indicate that it met, and red squares indicate that it did not meet. It is organised by profiles in the horizontal axis, and the target frequencies on the vertical axis. Each profile has two sections denoting strict and loose criteria. The profiles are ordered from most number of blue squares to least number of blue squares. This is when the device is at full on gain, so to meet individual profiles, the device volume has to be lowered by 5-10 dB SPL depending on the profile being considered. Overall, this shows how well the device fits the targets. We also showed how the profiles met in G.R.A.S KEMAR, where we show that without volume considerations, the profiles offer very good fit in Figure S3.

#### VII. SUPPLEMENTARY INDIVIDUAL GRAPHS OF AUDIOLOGICAL PROFILES AND TARGETS (DB SPL)

The following figures detail individual audiological profile, and their respective Targets (dB SPL) for each differing input (ISTS 55,65 dB SPL, and 80 dB SPL, and G.R.A.S KEMAR Left and Right Ears at 65 dB SPL). Meets Strict Criteria refers to the Response (dB SPL) being within 5 dB SPL of the target. Meets Loose Criteria Refers to the Response (dB SPL) being within 10 dB SPL of the target.

##### A. Male Age 60-69 Left Ear

###### 1. *ISTS 55 dB SPL*

Figure S13. Table I.

###### 2. *ISTS 65 dB SPL*

Figure S14. Table II.

###### 3. *ISTS 80 dB SPL*

Figure S15. Table III

###### 4. *KEMAR Right Ear 65 dB SPL*

. Figure S16. Table V.

###### 5. *KEMAR Left Ear 65 dB SPL*

Figure S17. Table IV.

##### B. Male Age 60-69 Right Ear

###### *1. ISTS 55 dB SPL*

Figure S18. Table VI

###### *2. ISTS 65 dB SPL*

Figure S19. Table VII.

###### *3. ISTS 80 dB SPL*

Figure S20. Table VIII.

###### *4. KEMAR Right Ear 65 dB SPL*

Figure S21. Table X.

###### *5. KEMAR Left Ear 65 dB SPL*

Figure S22. Table IX.

##### C. Male Age 70-79 Left Ear

###### *1. ISTS 55 dB SPL*

Figure S33. Table XXI.

###### *2. ISTS 65 dB SPL*

Figure S34. Table XXII.

###### *3. ISTS 80 dB SPL*

Figure S35. Table XXIII.

###### *4. KEMAR Right Ear 65 dB SPL*

Figure S36. Table XXIV.

###### *5. KEMAR Left Ear 65 dB SPL*

Figure S37. Table XXV.

##### D. Male Age 70-79 Right Ear

###### *1. ISTS 55 dB SPL*

Figure S28. Table XVI.

###### *2. ISTS 65 dB SPL*

Figure S29. Table XVII.

###### *3. ISTS 80 dB SPL*

Figure S30. Table XVIII.

###### *4. KEMAR Right Ear 65 dB SPL*

Figure S31. Table XIX.

###### *5. KEMAR Left Ear 65 dB SPL*

Figure S32. Table XX.

##### E. Female Age 60-69 Left Ear

###### *1. ISTS 55 dB SPL*

Figure S38. Table XXVI.

###### *2. ISTS 65 dB SPL*

Figure S39. Table XXVII.

###### *3. ISTS 80 dB SPL*

Figure S40. Table XXVIII.

###### *4. KEMAR Right Ear 65 dB SPL*

Figure S41. Table XXIX.

###### *5. KEMAR Left Ear 65 dB SPL*

Figure S42. Table XXX.

#### F. Female Age 60-69 Right Ear

##### 1. *ISTS 55 dB SPL*

Figure S43. Table XXXI.

##### 2. *ISTS 65 dB SPL*

Figure S44. Table XXXII.

##### 3. *ISTS 80 dB SPL*

Figure S45. Table XXXIII.

##### 4. *KEMAR Right Ear 65 dB SPL*

Figure S46. Table XXXIV.

##### 5. *KEMAR Left Ear 65 dB SPL*

Figure S47. Table XXXV.

#### G. Female Age 70-79 Left Ear

##### 1. *ISTS 55 dB SPL*

Figure S58. Table XXXVI.

##### 2. *ISTS 65 dB SPL*

Figure S59. Table XXXVII.

##### 3. *ISTS 80 dB SPL*

Figure S60. Table XXXVIII.

##### 4. *KEMAR Right Ear 65 dB SPL*

Figure S61. Table XXXIX.

##### 5. *KEMAR Left Ear 65 dB SPL*

Figure S62. Table XL.

#### H. Female Age 70-79 Right Ear

##### 1. *ISTS 55 dB SPL*

Figure S53. Table XLI.

##### 2. *ISTS 65 dB SPL*

Figure S54. Table XLII.

##### 3. *ISTS 80 dB SPL*

Figure S55. Table XLIII.

##### 4. *KEMAR Right Ear 65 dB SPL*

Figure S56. Table XLIV.

##### 5. *KEMAR Left Ear 65 dB SPL*

Figure S57. Table XLV.

#### I. Hearing Profile X

##### 1. *ISTS 55 dB SPL*

Figure S63. Table XLVI.

##### 2. *ISTS 65 dB SPL*

Figure S64. Table XLVII.

##### 3. *ISTS 80 dB SPL*

Figure S65. Table XLVIII.

##### 4. *KEMAR Right Ear 65 dB SPL*

Figure S66. Table XLIX.

##### 5. *KEMAR Left Ear 65 dB SPL*

Figure S67. Table L.

##### J. Hearing Profile Y

###### *1. ISTS 55 dB SPL*

Figure S68. Table LI.

###### *2. ISTS 65 dB SPL*

Figure S69. Table LII.

###### *3. ISTS 80 dB SPL*

Figure S70. Table LIII.

###### *4. KEMAR Right Ear 65 dB SPL*

Figure S71. Table LIV.

###### *5. KEMAR Left Ear 65 dB SPL*

Figure S72. Table LV.

##### K. Hearing Profile Y

###### *1. ISTS 55 dB SPL*

Figure S68. Table LI.

###### *2. ISTS 65 dB SPL*

Figure S69. Table LII.

###### *3. ISTS 80 dB SPL*

Figure S70. Table LIII.

###### *4. KEMAR Right Ear 65 dB SPL*

Figure S71. Table LIV.

###### *5. KEMAR Left Ear 65 dB SPL*

Figure S72. Table LV.

##### L. Hearing Profile Z

###### *1. ISTS 55 dB SPL*

Figure S73. Table LVI.

###### *2. ISTS 65 dB SPL*

Figure S74. Table LVII.

###### *3. ISTS 80 dB SPL*

Figure S75. Table LVIII.

###### *4. KEMAR Right Ear 65 dB SPL*

Figure S76. Table LIX.

###### *5. KEMAR Left Ear 65 dB SPL*

Figure S77. Table LX

[6] Maxim Integrated Products, "Microphone Amplifier with AGC and Low-Noise Microphone Bias MAX9814 Microphone Amplifier with AGC and Low-Noise Microphone Bias: Data Sheet," Maxim Integrated Products (2016).

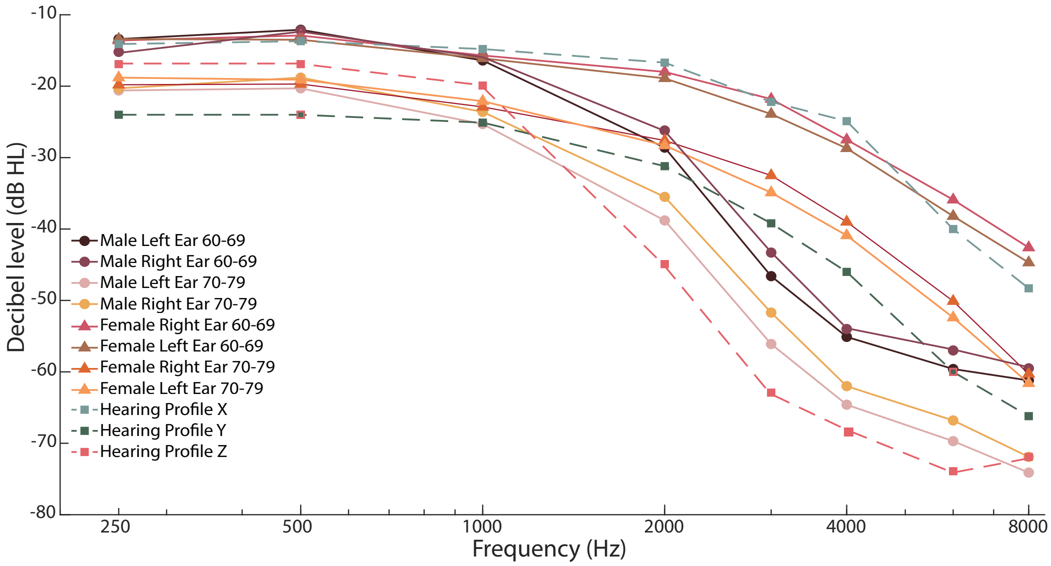


FIG. S1. Shows the 12 profiles used in the study detailing Males and Females both left and right ears from ages 60-79. The figure uses data pulled from two studies [2], [1]. Cruikshanks, et al from 1993-1995 [1] conducted hearing tests of both ears of people living in Beaver Dam, Wisconsion, and Ciletti and Flamme [2] who conducted hearing loss analysis cross country in USA. Cruikshanks, et al from 1993-1995 conducted hearing tests of both ears of people living in Beaver Dam, Wisconsion, and pulls data from 566 females from the age of 60-69, 534 females from ages 70-79, 489 males from ages 60-69, and 355 males from the ages of 70-79 and detail left and right ear hearing losses. Hearing Profiles X,Y, and Z are gender neutral audiograms of increasing severity displaying ARHL characteristics pulled from a study done by Ciletti and Flamme, which was conducted over 1999-2005, based on the National Health and Nutrition Examination Survey (NHANES) which was a cross-country sample study of common audiogram trends, based on 2819 women, and 2525 men ages 20-69. Hearing Profile X (mild) is exhibited by 204 men, 282 women; Hearing Profile Y (moderate) is exhibited by 123 women, and 145 men; Hearing Profile Z (Severe) is exhibited by 174 men, 19 women.

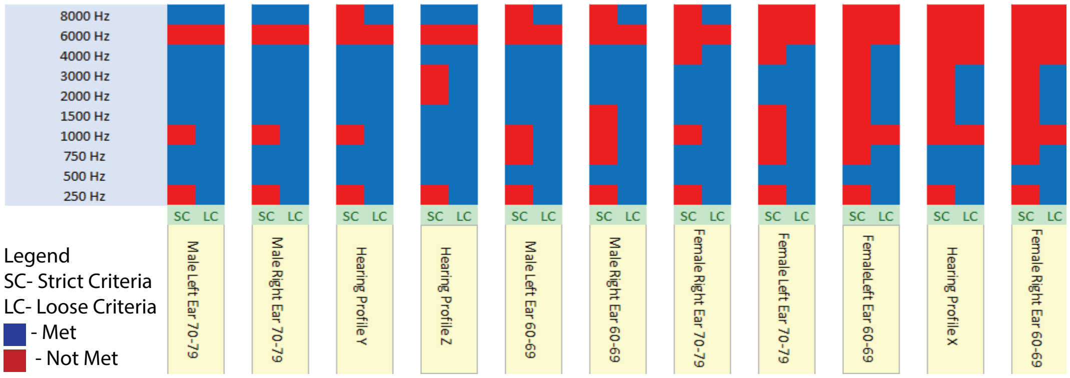


FIG. S2. Shows the quantification of fit in a visual matter for Speechmap. It is organised by profiles in the horizontal axis, and the target frequencies on the vertical axis. Each profile has two sections denoting strict and loose criteria. The profiles are ordered from most number of blue squares to least number of blue squares. This is when the device is at full on gain, so to meet individual profiles, the device volume has to be lowered by 5-10 dB SPL depending on the profile being considered.

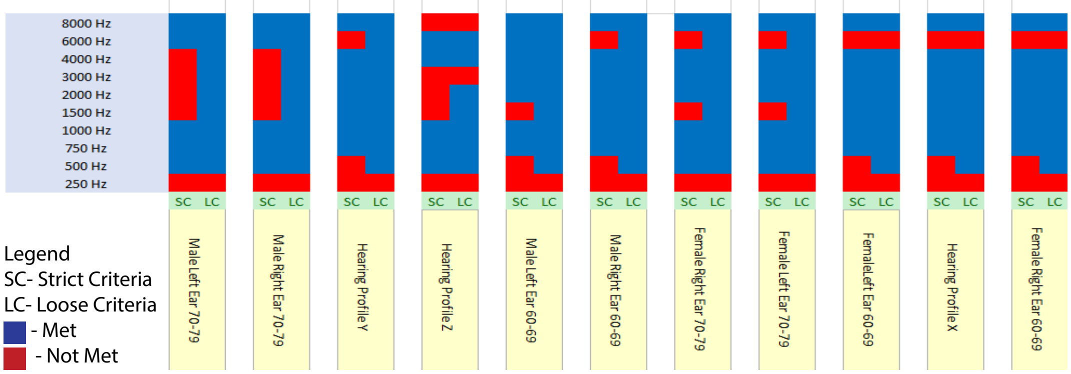


FIG. S3. Shows the quantification of fit in a visual matter for G.R.A.S KEMAR Response. It is organised by profiles in the horizontal axis, and the target frequencies on the vertical axis. Each profile has two sections denoting strict and loose criteria. The profiles are ordered from most number of blue squares to least number of blue squares. This is when the device is at full on gain, and shows that we do not necessarily need to lower the volume as the G.R.A.S KEMAR shows a better fit.

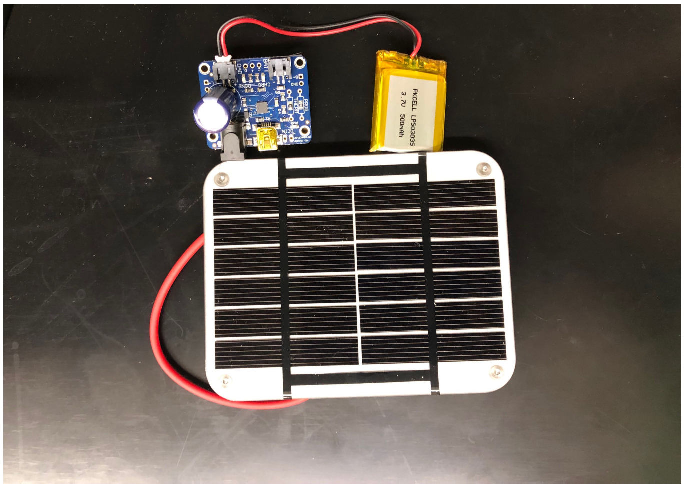

FIG. S4. Shows a solar panel, adapter, and lithium-ion packet battery for a rechargeable station. The costs are obtained: solar panel $1.85 (A Grade Small Solar Panel 1w 3w 5w 6v 9v 12v Solar Panel Low Price Mini Solar Panel from www.alibaba.com, from Shangdong, China, manufactured by Hinergy Energy, M/N HNP1W-5W6V); adapter $17.50 (USB / DC / Solar Lithium Ion/Polymer charger - v2 from www.adafruit.com, P/N 390), Lithium-ion packet battery $5.95 (Lithium Ion Polymer Battery with Short Cable - 3.7V 350mAh, from www.adafruit.com, P/N 4237).

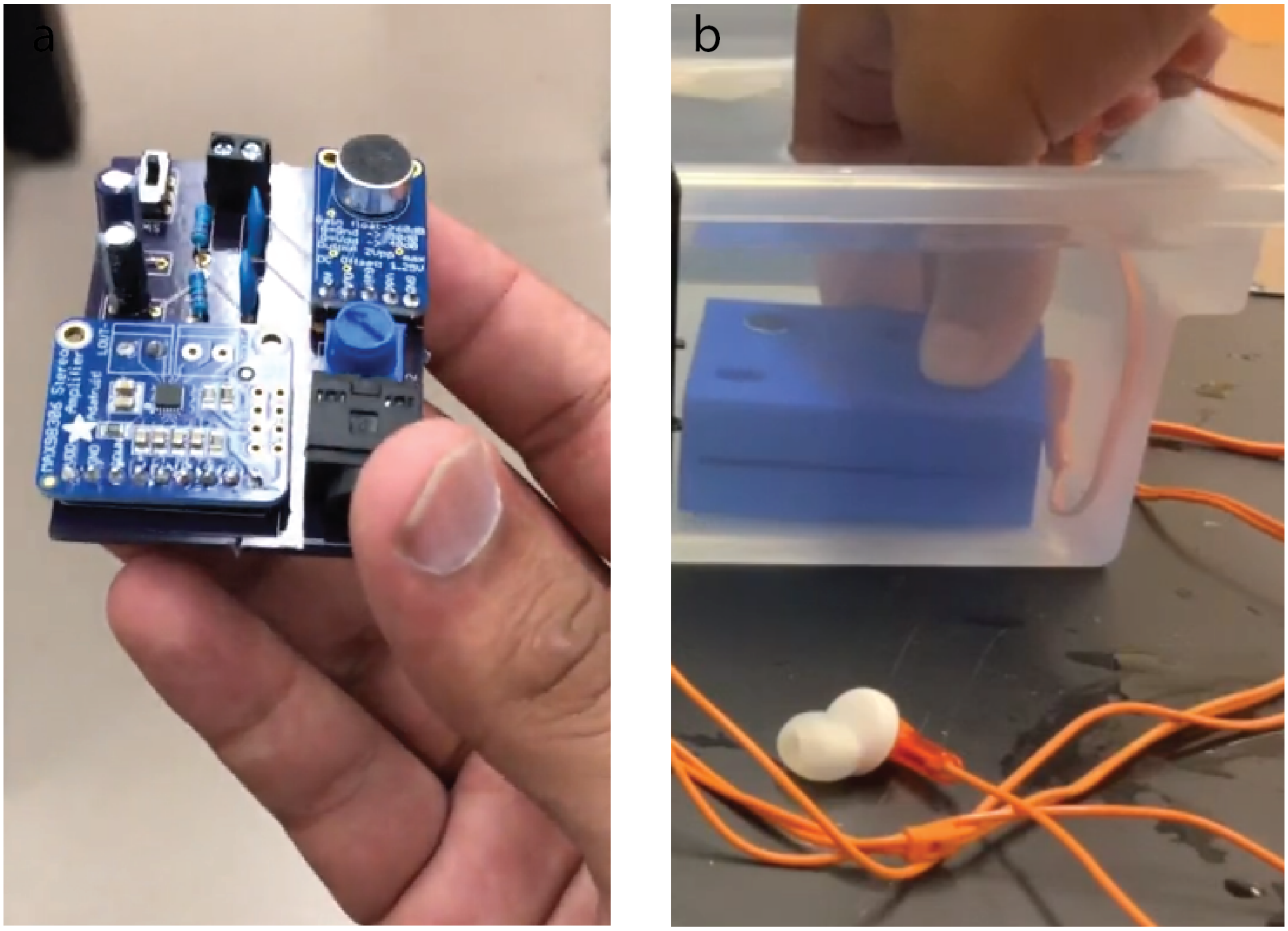


FIG. S5. **a** Still from SI Movie 4 detailing drop test of the device. **b** Still from SI Movie 3 detailing water test of the device.

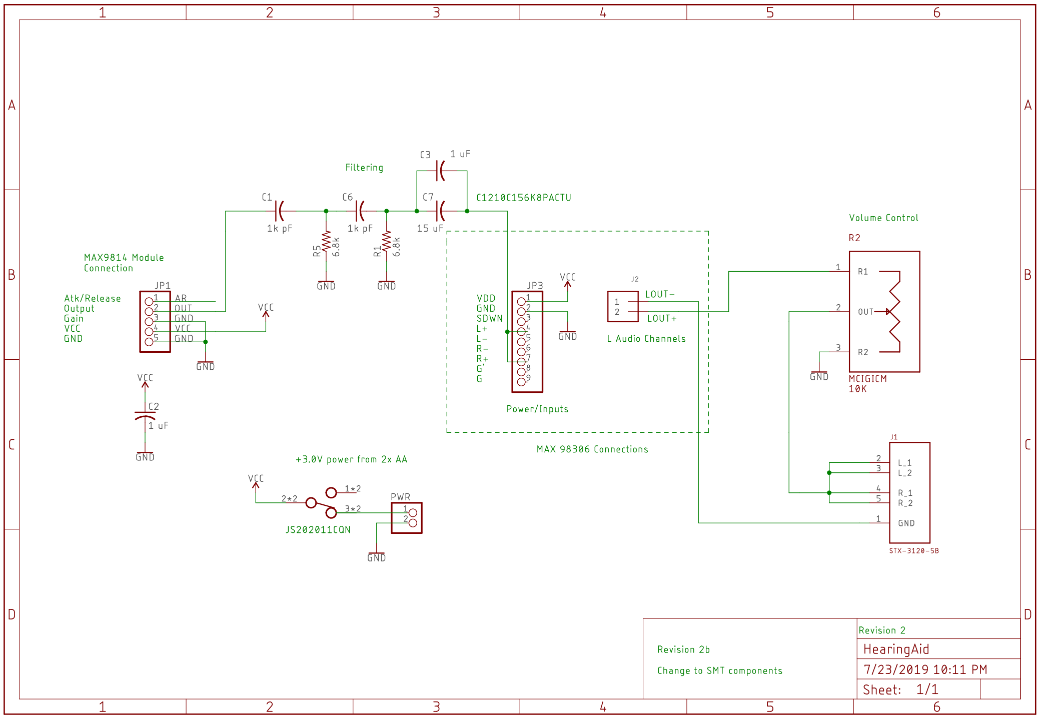


FIG. S6. Shows the schematic of the LoCHAid.

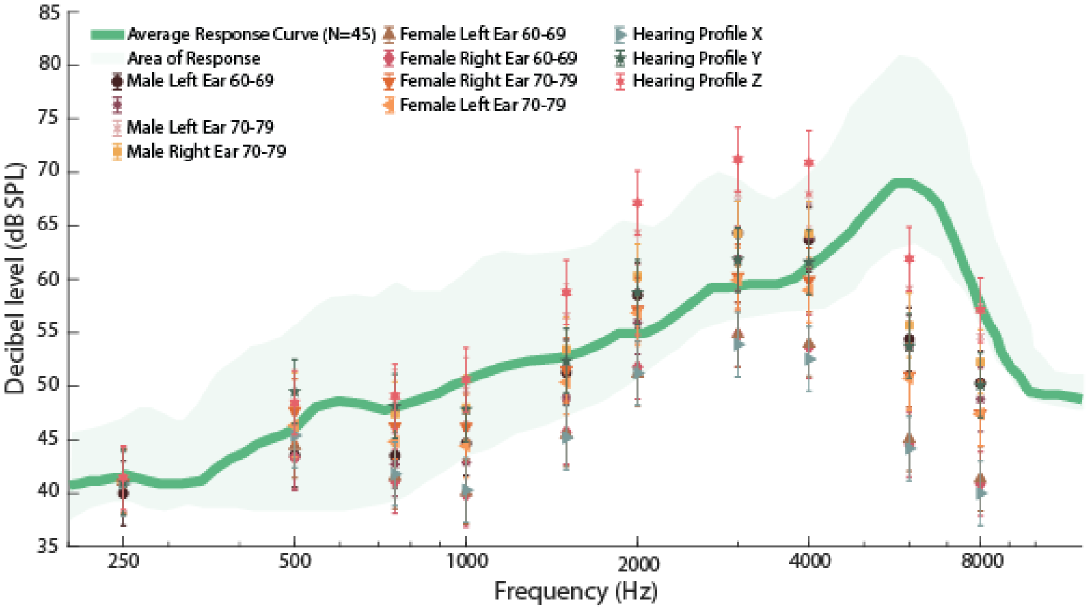


FIG. S7. Shows the ISTS Response (dB SPL) Curve for 55 dB SPL ISTS Input with Targets (dB SPL) for all profiles tested.

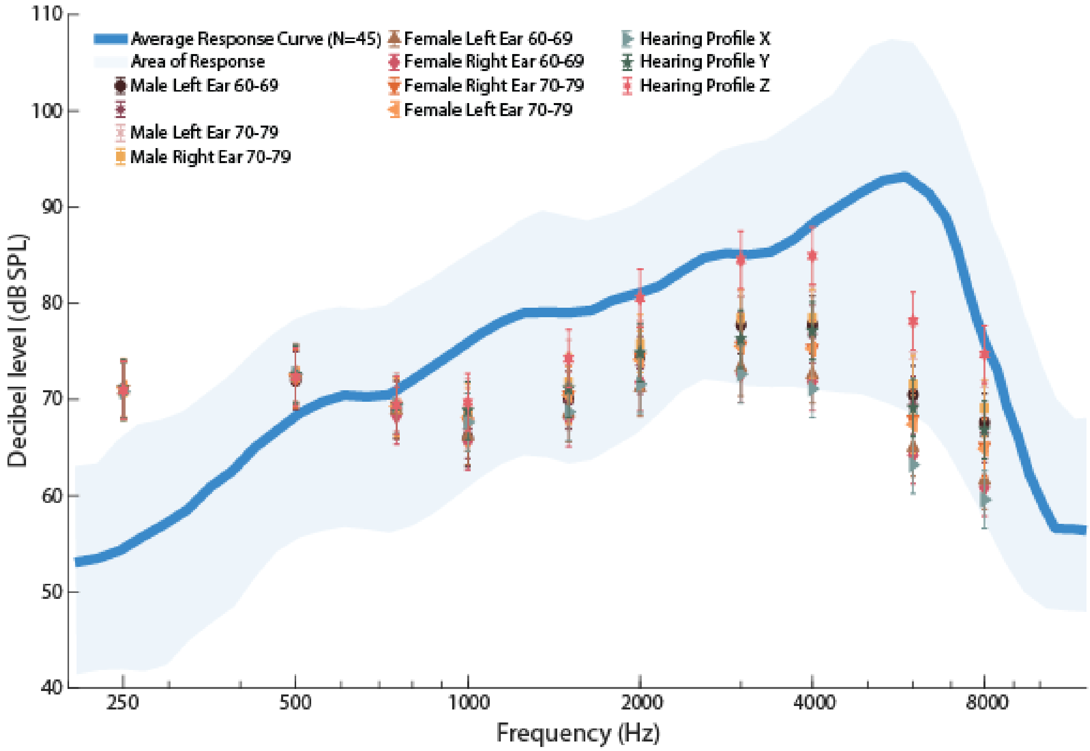


FIG. S8. Shows the ISTS Response (dB SPL) Curve for 80 dB SPL ISTS Input with Targets (dB SPL) for all profiles tested.

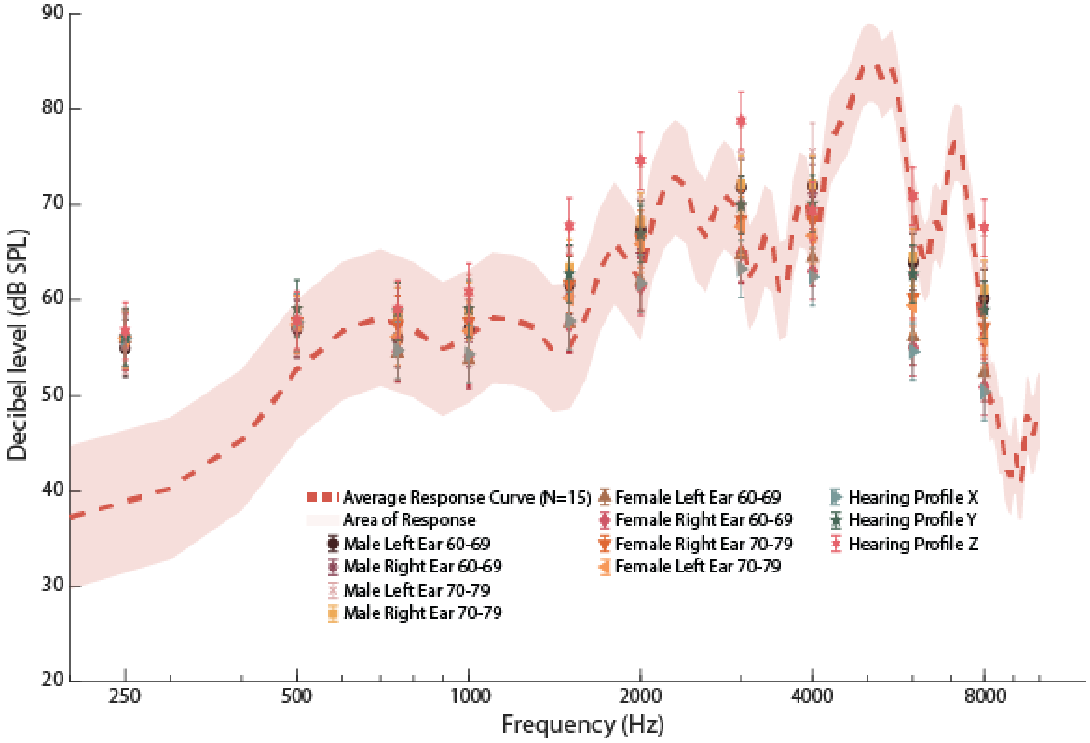


FIG. S9. Shows the KEMAR Response (dB SPL) Curve (Right Ear) for 65 dB SPL ISTS Input with Targets (dB SPL) for all profiles tested.

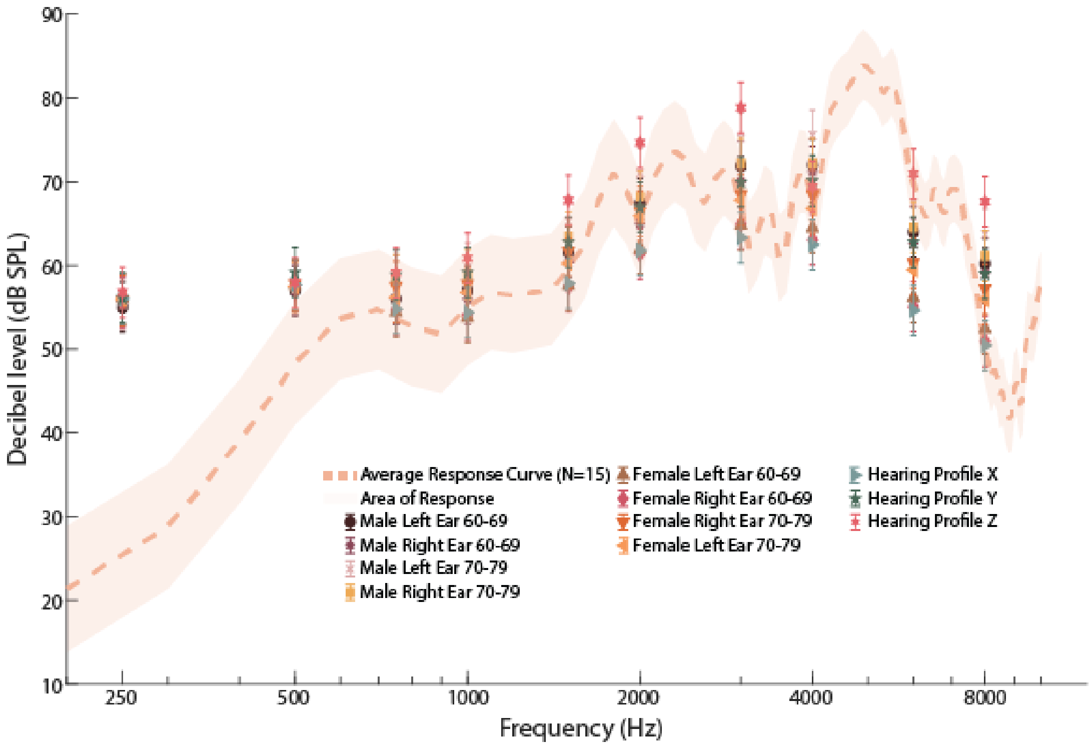


FIG. S10. Shows the KEMAR Response (dB SPL) Curve (Left Ear) for 65 dB SPL ISTS Input with Targets (dB SPL) for all profiles tested.

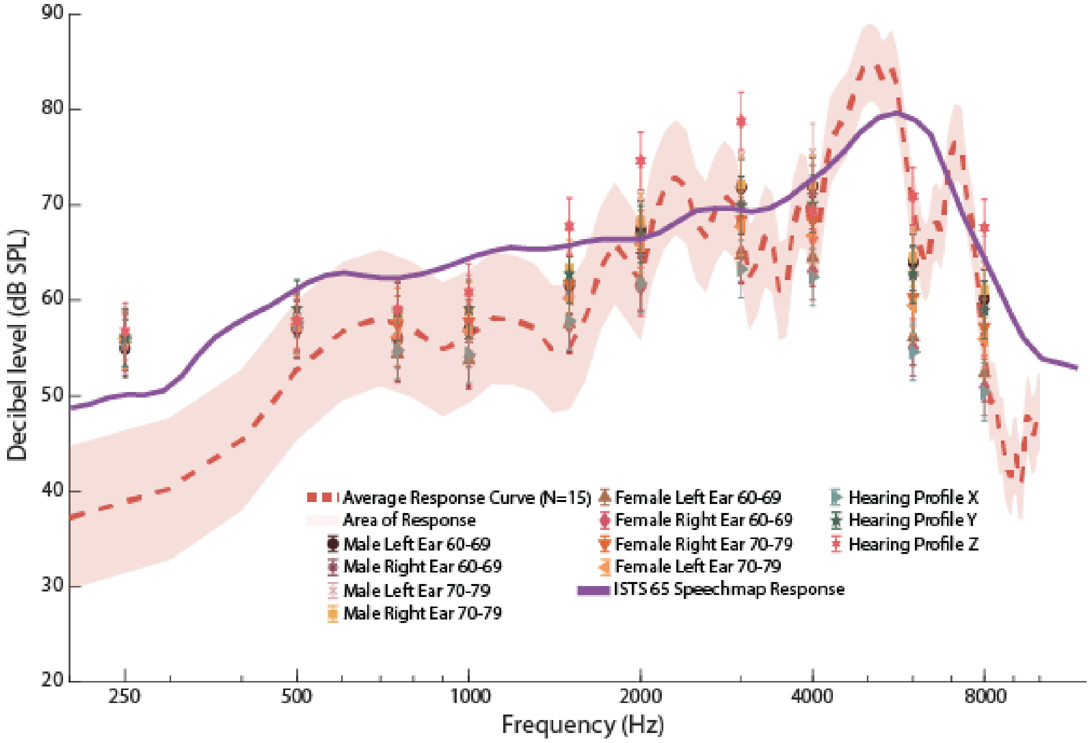


FIG. S11. Shows the KEMAR Response (dB SPL) Curve (Right Ear) for 65 dB SPL ISTS Input with Targets (dB SPL) for all profiles tested with ISTS 65 dB SPL Speechmap Response (dB SPL).

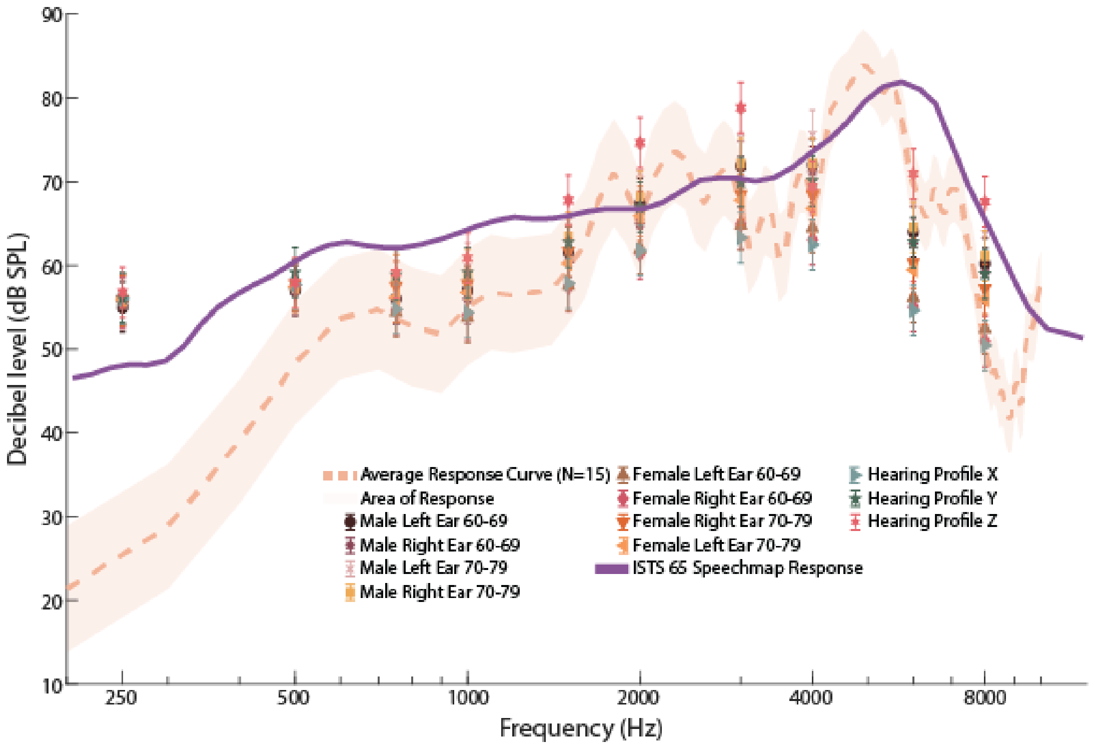


FIG. S12. Shows the KEMAR Response (dB SPL) Curve (Left Ear) for 65 dB SPL ISTS Input with Targets (dB SPL) for all profiles tested with ISTS 65 dB SPL Speechmap Response (dB SPL).

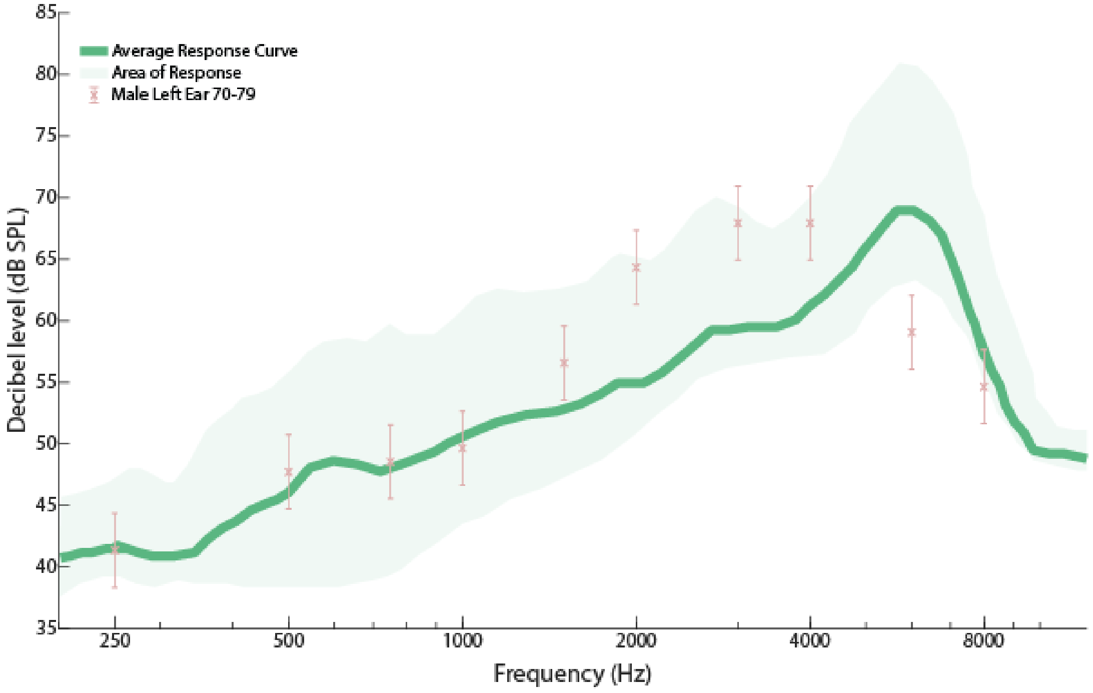


FIG. S13. Shows the ISTS 55 dB SPL Response (dB SPL) Curve with Male 60-69 Left Ear Targets (dB SPL).

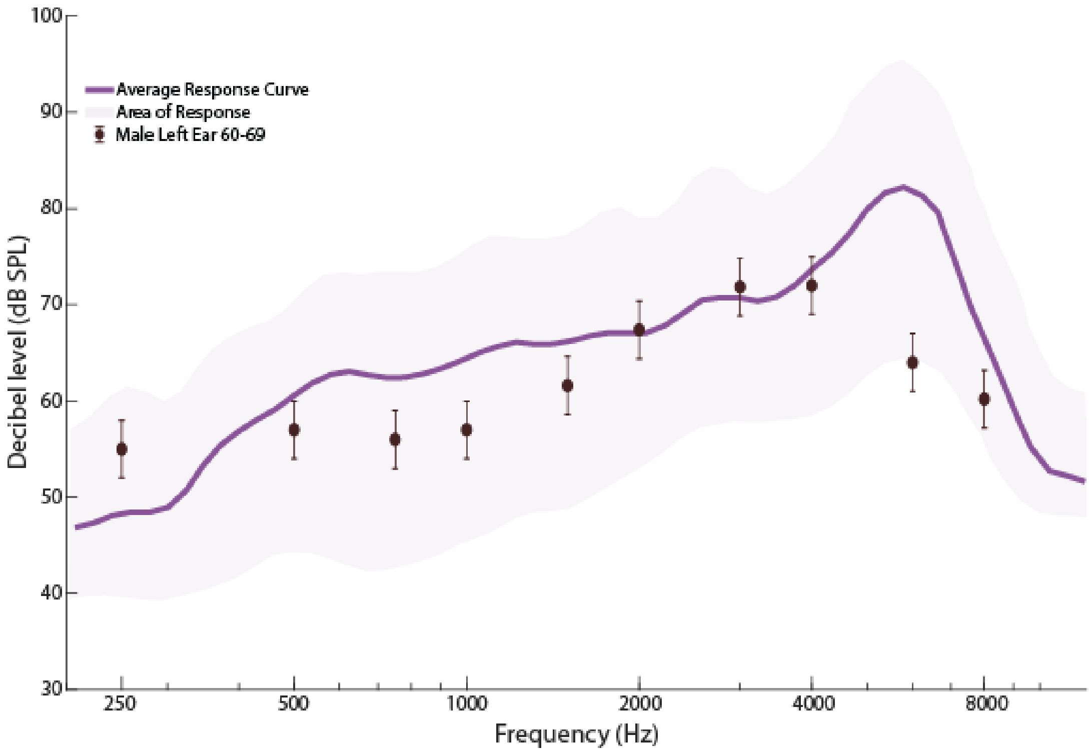


FIG. S14. Shows the ISTS 65 dB SPL Response (dB SPL) Curve with Male 60-69 Left Ear Targets (dB SPL).

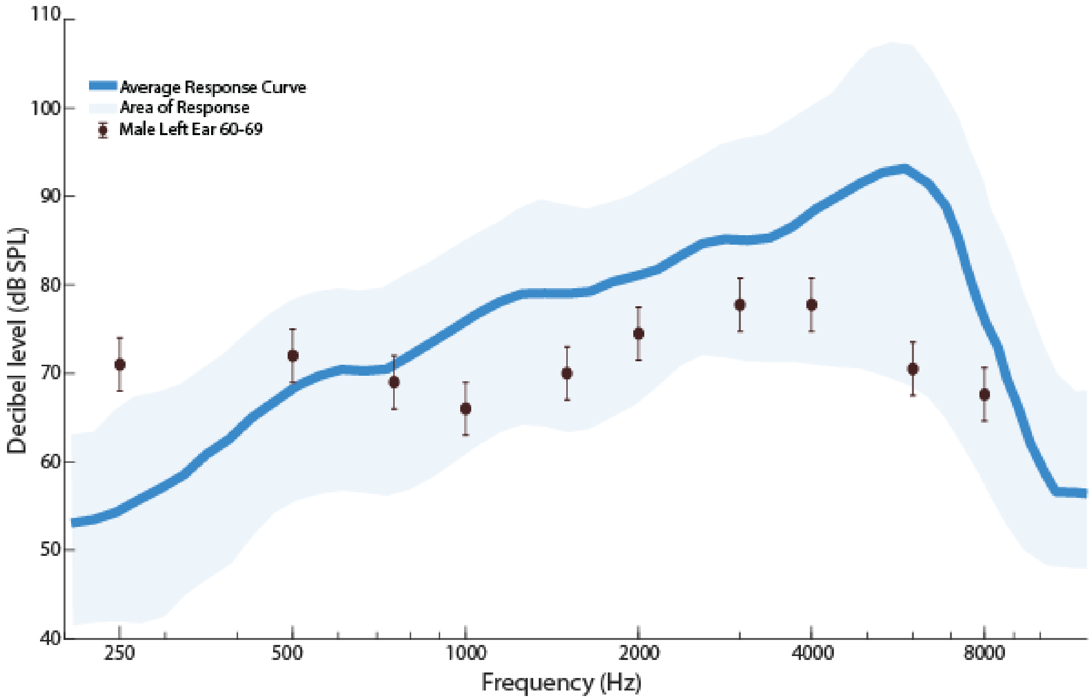


FIG. S15. Shows the ISTS 80 dB SPL Response (dB SPL) Curve with Male 60-69 Left Ear Targets (dB SPL).

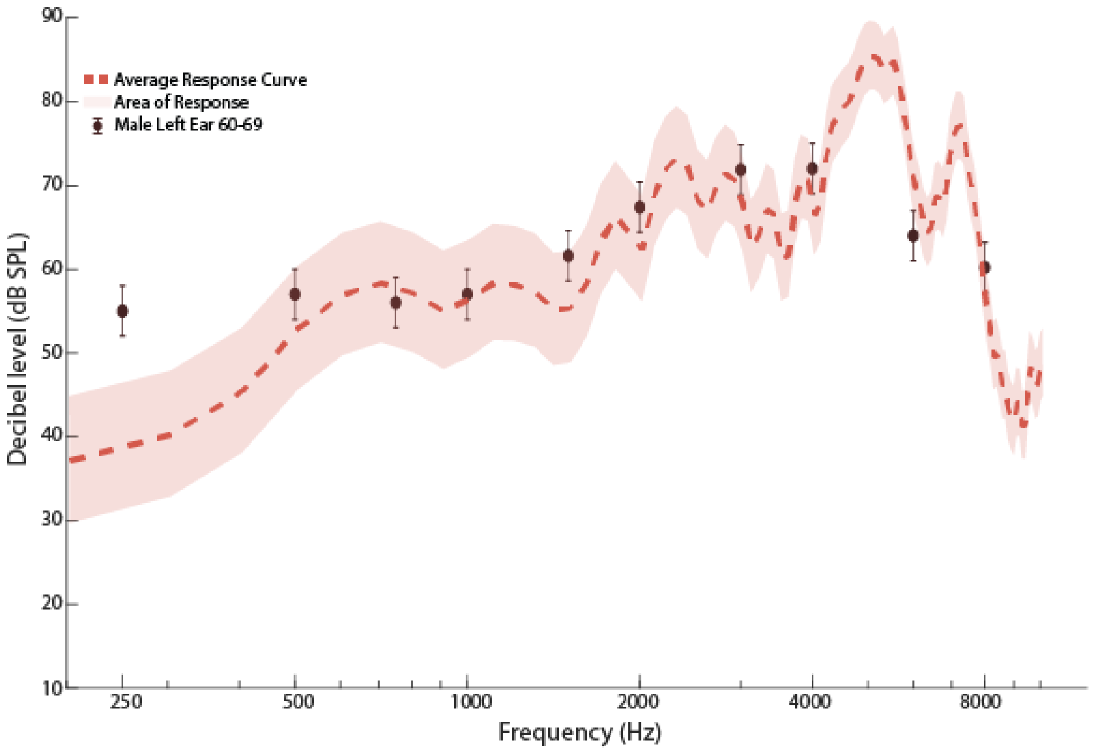


FIG. S16. Shows the G.R.A.S KEMAR 65 db SPL ISTS input Response (dB SPL) Curve (Right Ear) with Male 60-69 Left Ear Targets (dB SPL).

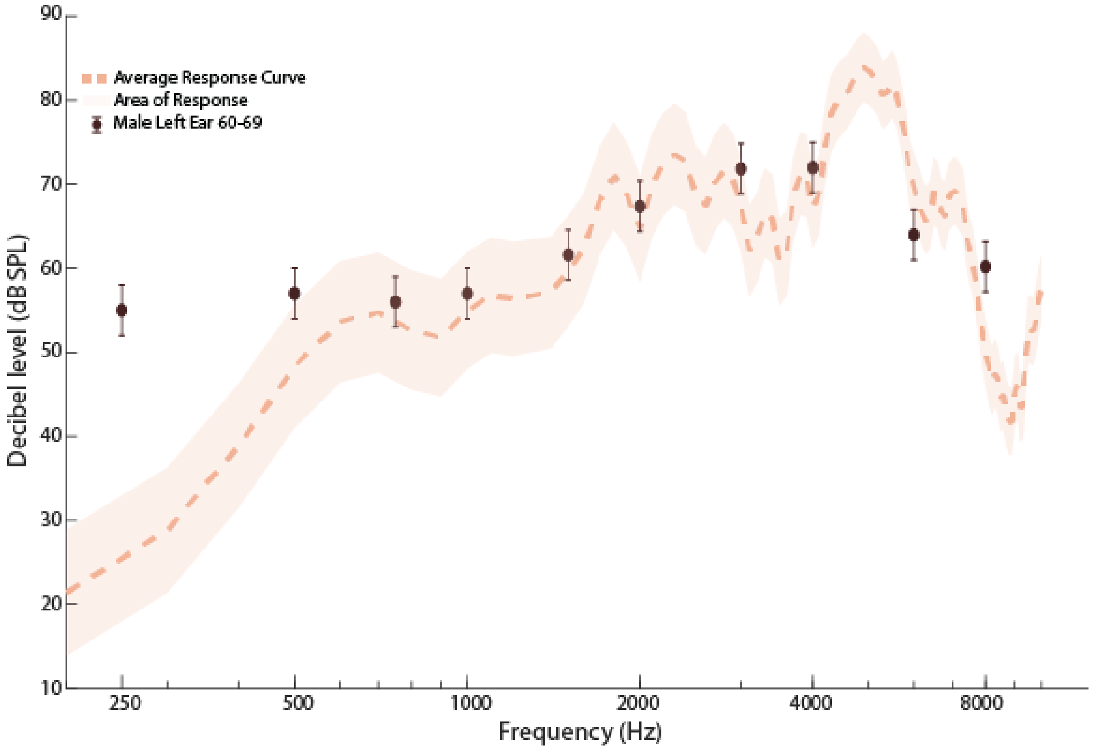


FIG. S17. Shows the G.R.A.S KEMAR 65 db SPL ISTS input Response (dB SPL) Curve (Left Ear) with Male 60-69 Left Ear Targets (dB SPL).

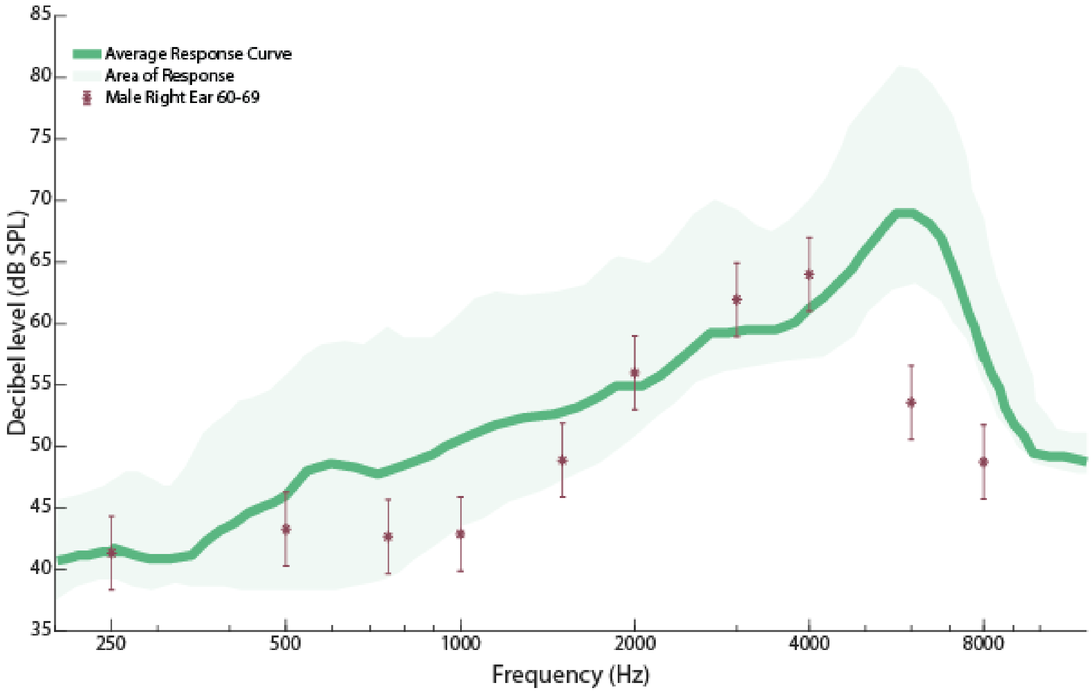


FIG. S18. Shows the ISTS 55 dB SPL Response (dB SPL) Curve with Male 60-69 Right Ear Targets (dB SPL).

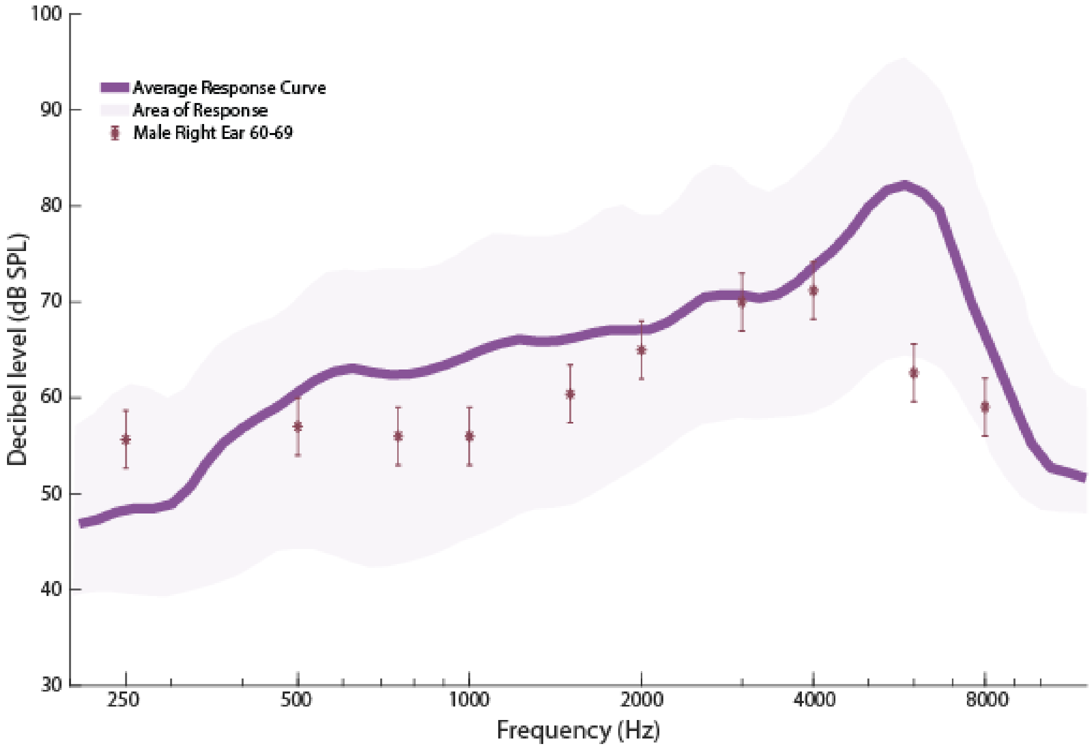


FIG. S19. Shows the ISTS 65 dB SPL Response (dB SPL) Curve with Male 60-69 Right Ear Targets (dB SPL).

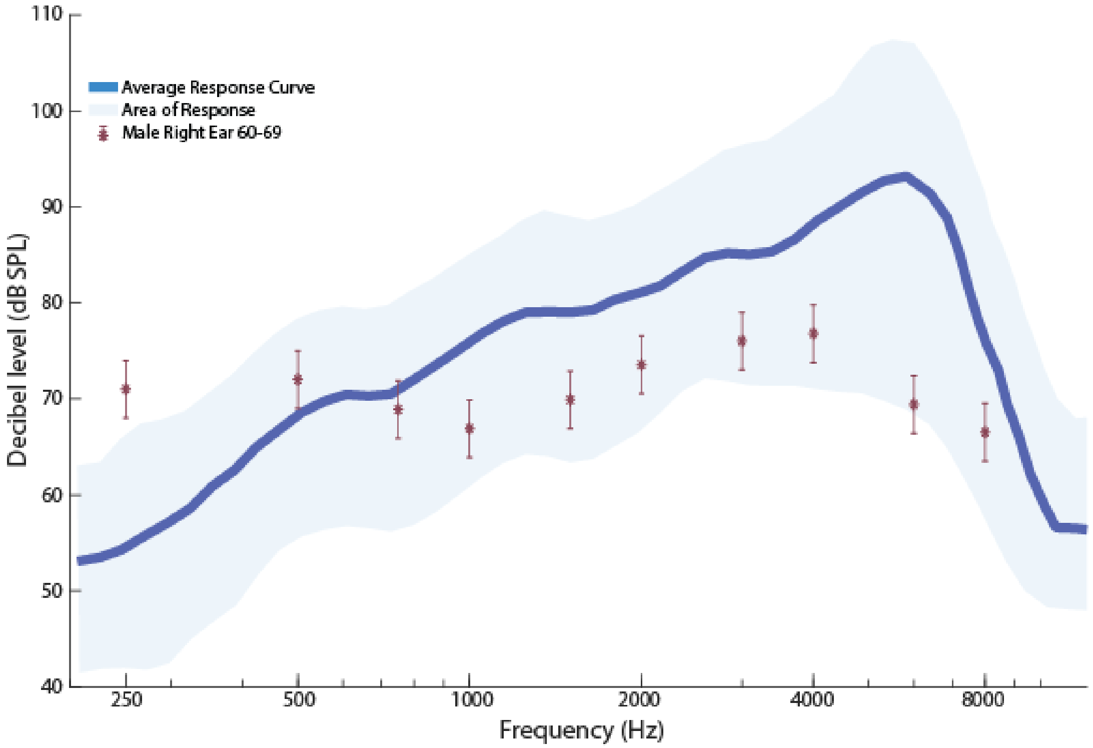


FIG. S20. Shows the ISTS 80 dB SPL Response (dB SPL) Curve with Male 60-69 Right Ear Targets (dB SPL).

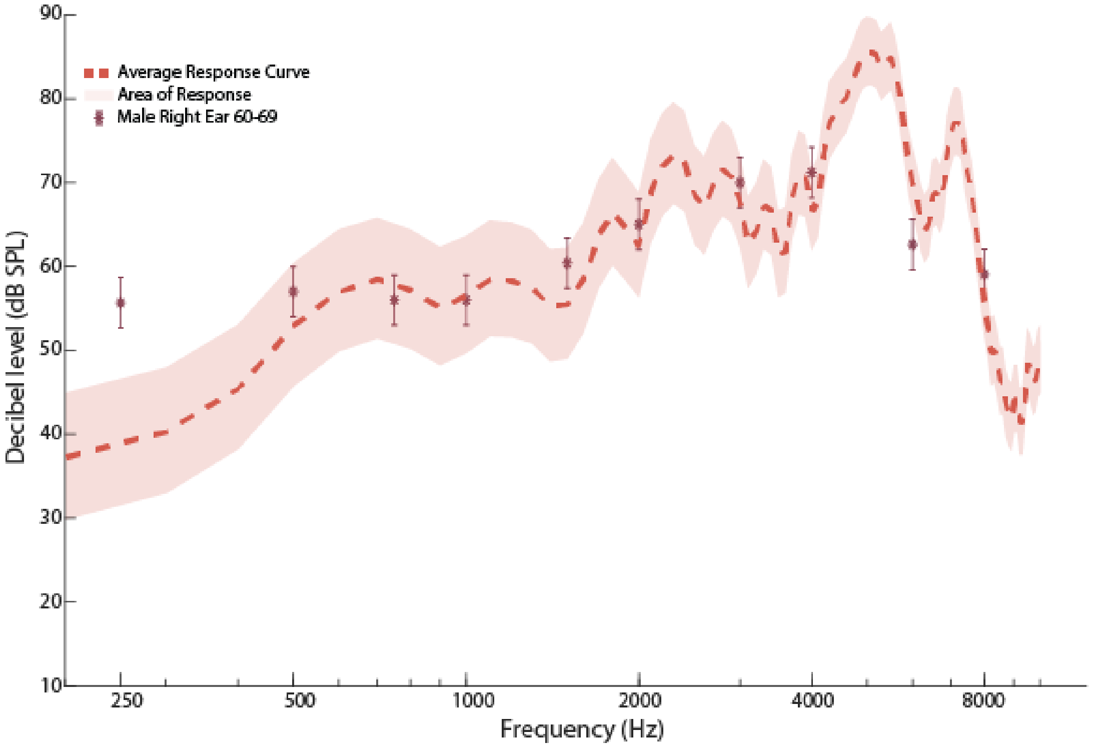


FIG. S21. Shows the G.R.A.S KEMAR 65 db SPL ISTS input Response (dB SPL) Curve (Right Ear) with Male 60-69 Right Ear Targets (dB SPL).

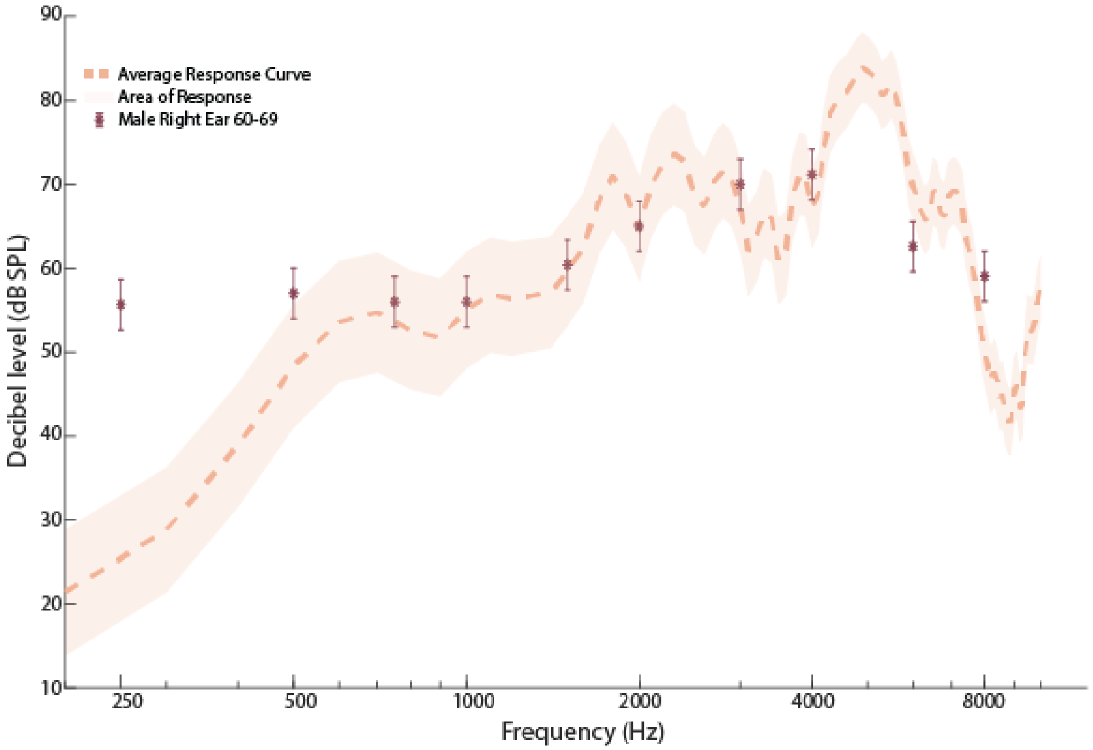


FIG. S22. Shows the G.R.A.S KEMAR 65 db SPL ISTS input Response (dB SPL) Curve (Left Ear) with Male 60-69 Right Ear Targets (dB SPL).

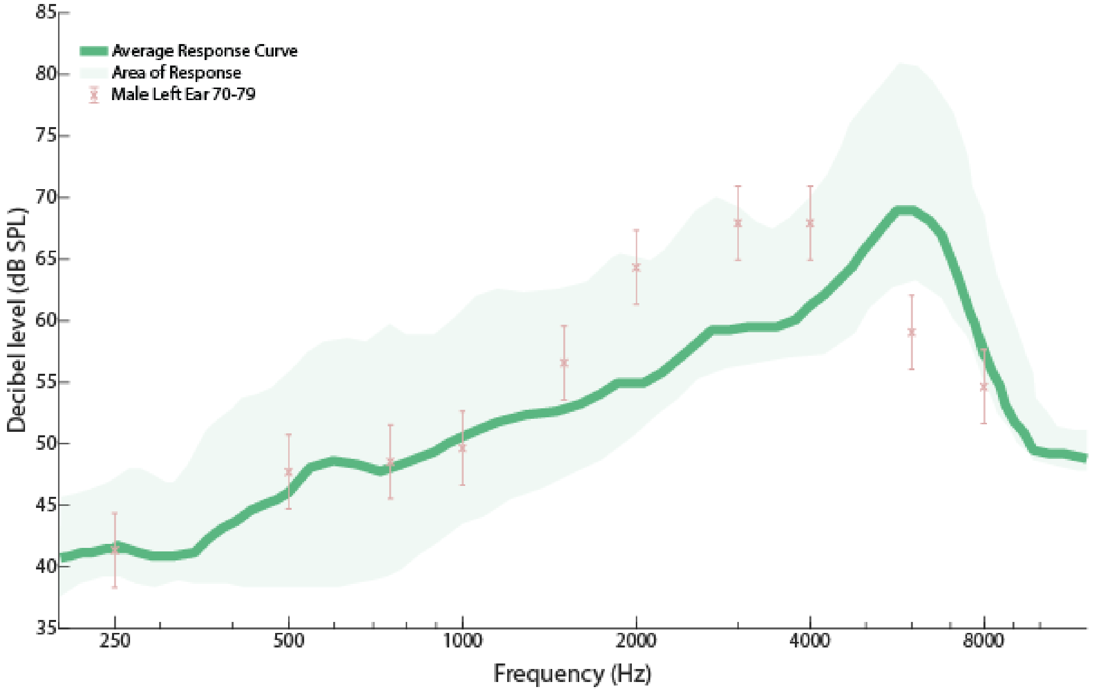


FIG. S23. Shows the ISTS 55 dB SPL Response (dB SPL) Curve with Male 70-79 Left Ear Targets (dB SPL).

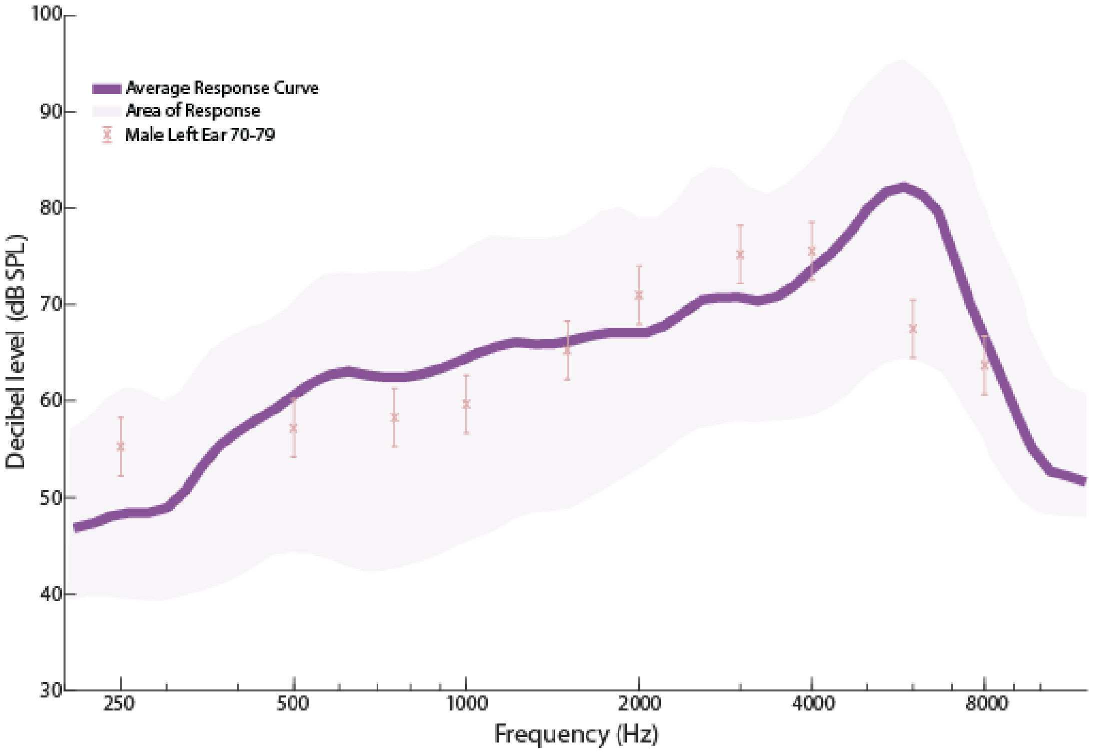


FIG. S24. Shows the ISTS 65 dB SPL Response (dB SPL) Curve with Male 70-79 Left Ear Targets (dB SPL).

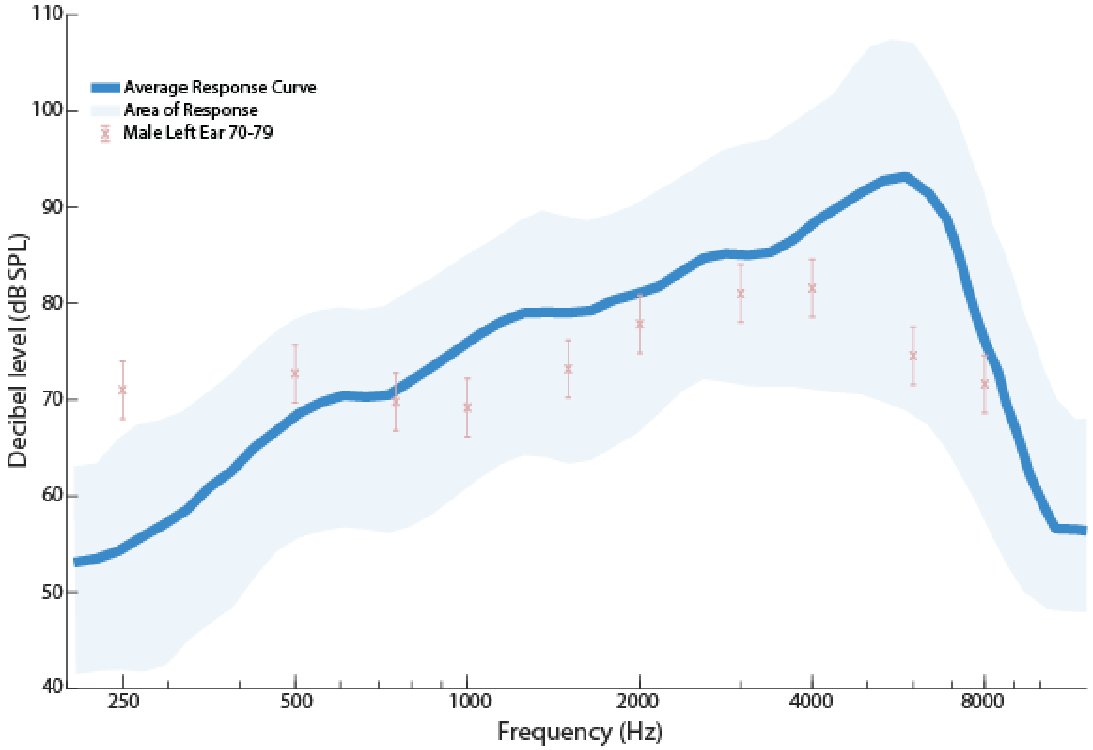


FIG. S25. Shows the ISTS 80 dB SPL Response (dB SPL) Curve with Male 70-79 Left Ear Targets (dB SPL).

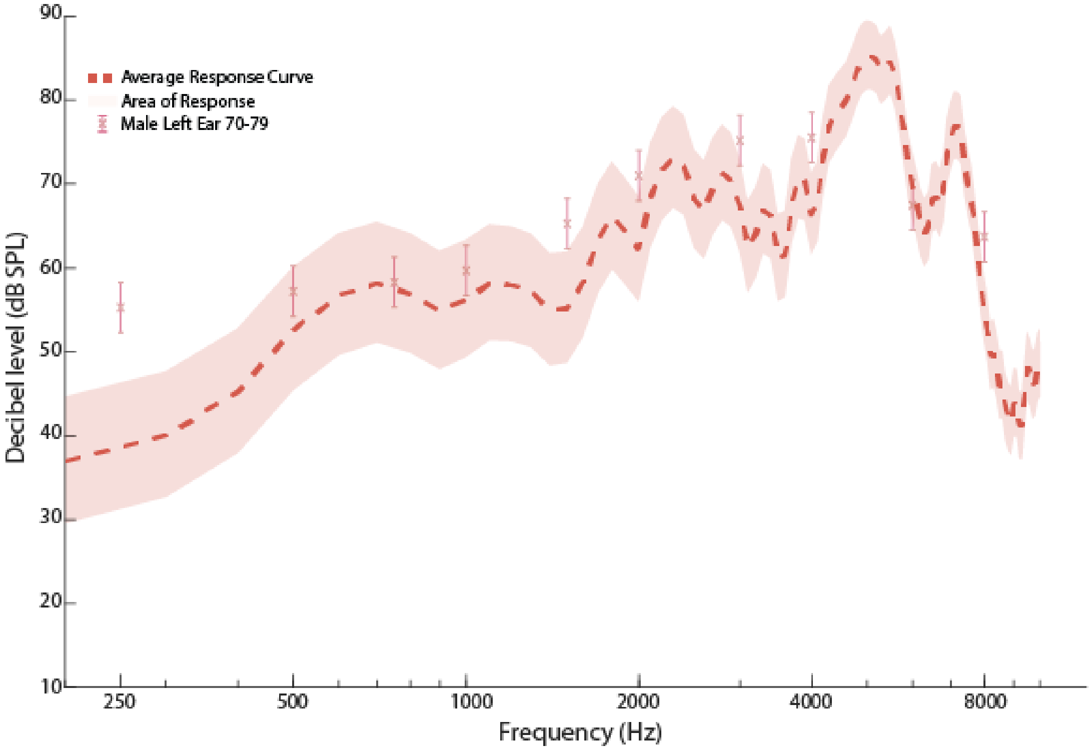


FIG. S26. Shows the G.R.A.S KEMAR 65 db SPL ISTS input Response (dB SPL) Curve (Right Ear) with Male 70-79 Left Ear Targets (dB SPL).

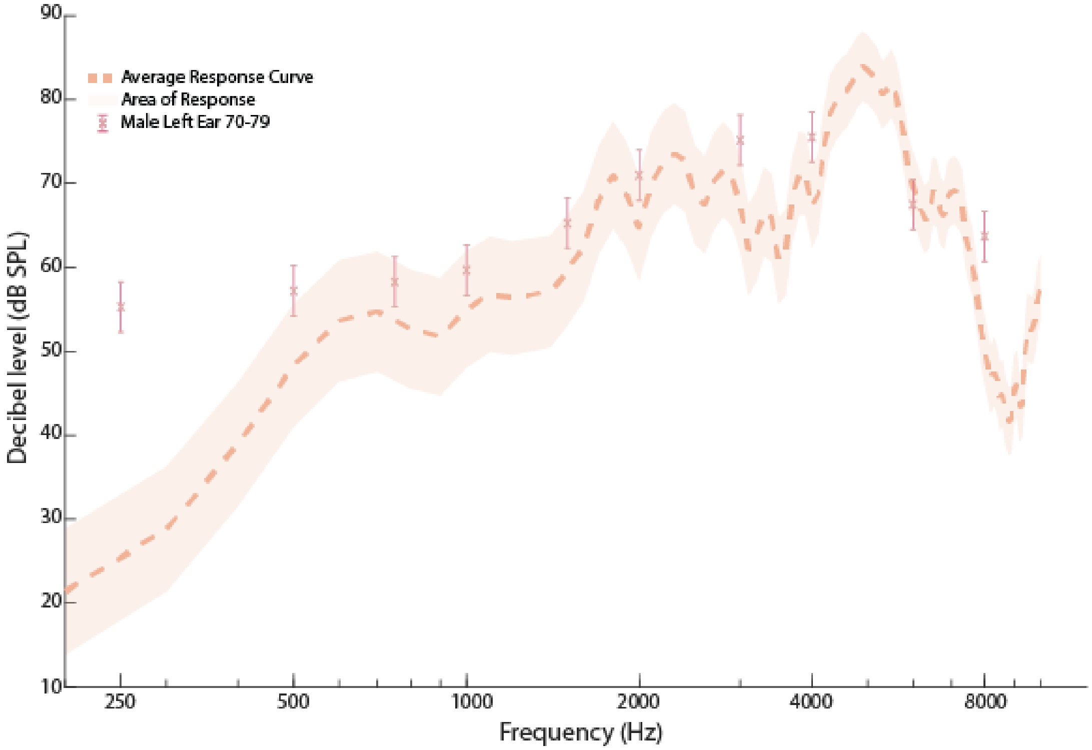


FIG. S27. Shows the G.R.A.S KEMAR 65 db SPL ISTS input Response (dB SPL) Curve (Left Ear) with Male 70-79 Left Ear Targets (dB SPL).

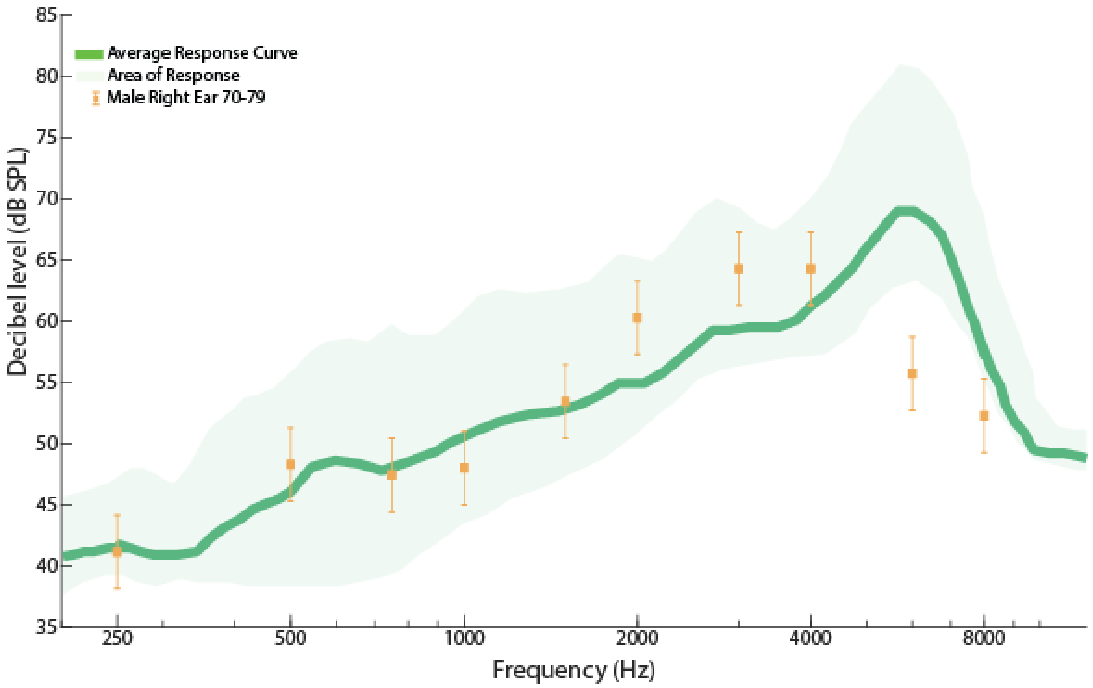


FIG. S28. Shows the ISTS 55 dB SPL Response (dB SPL) Curve with Male 70-79 Right Ear Targets (dB SPL).

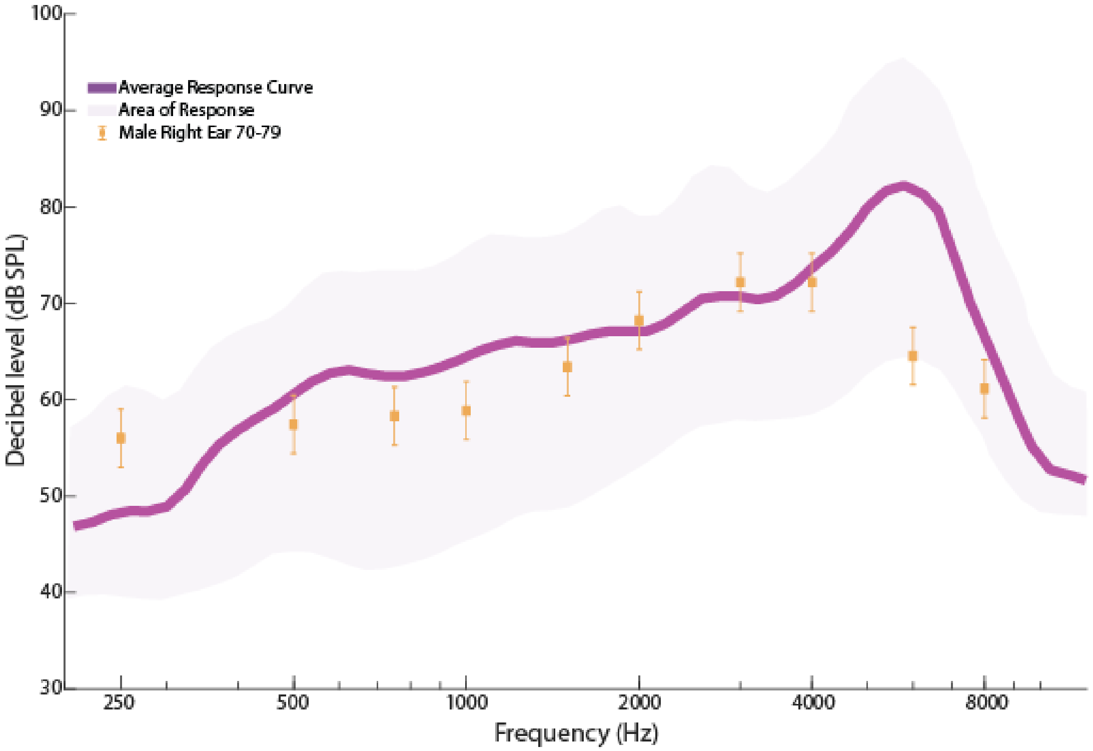


FIG. S29. Shows the ISTS 65 dB SPL Response (dB SPL) Curve with Male 70-79 Right Ear Targets (dB SPL).

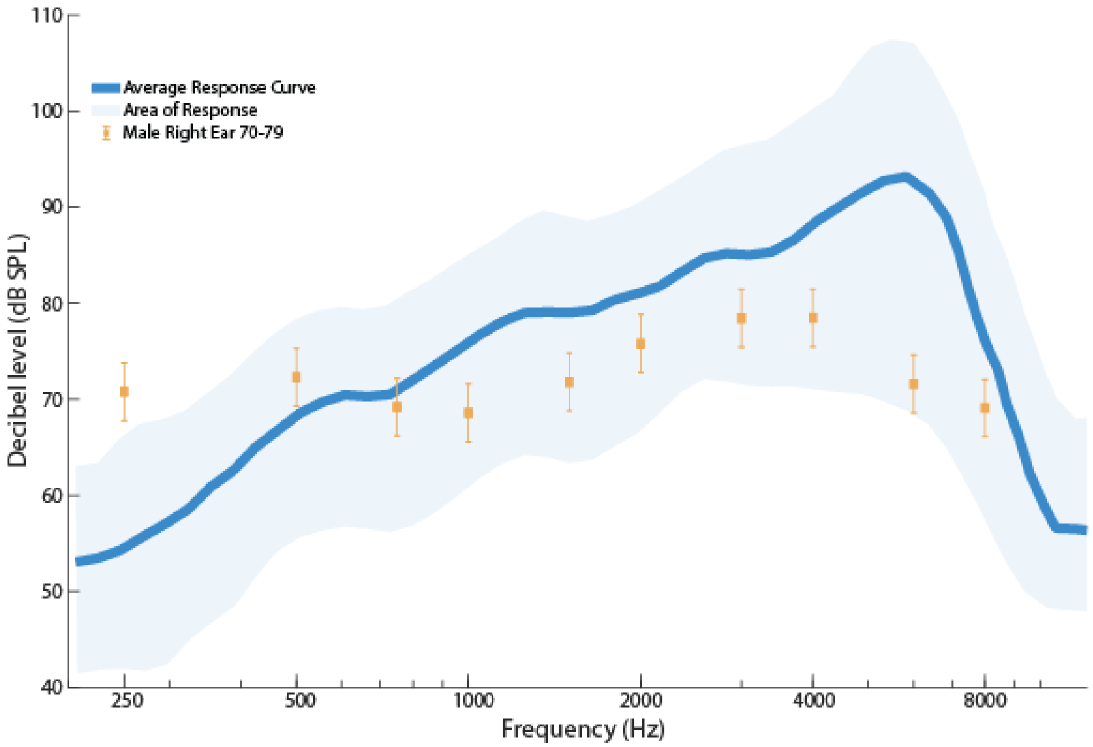


FIG. S30. Shows the ISTS 80 dB SPL Response (dB SPL) Curve with Male 70-79 Right Ear Targets (dB SPL).

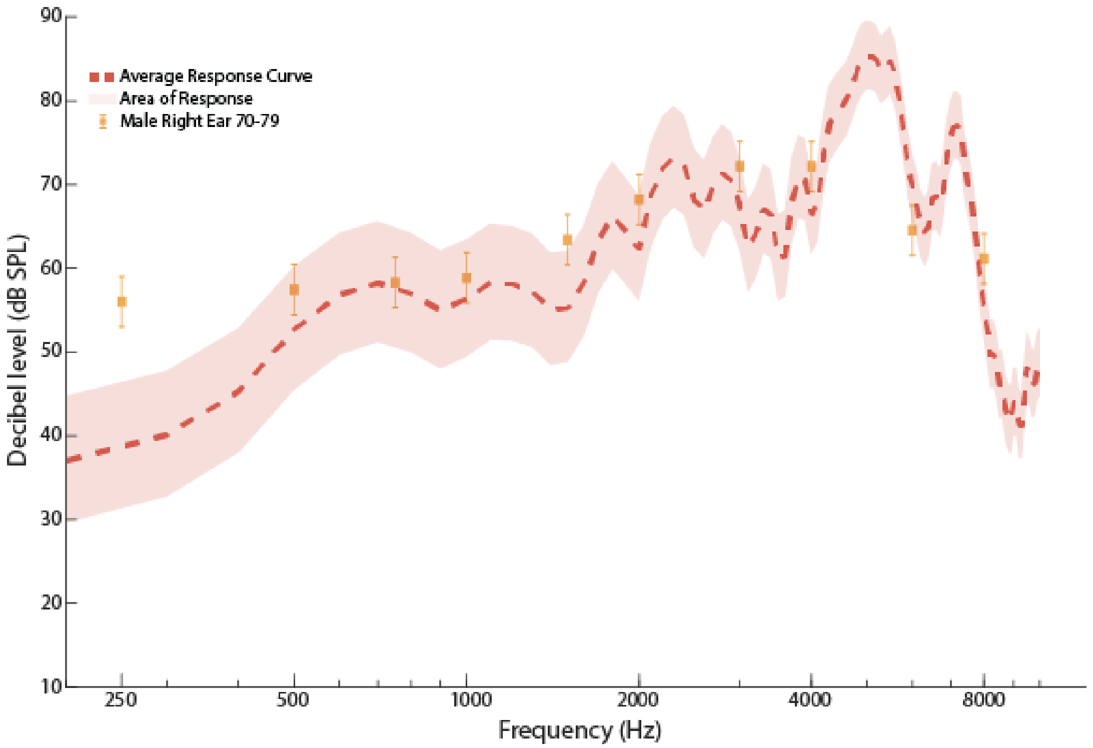


FIG. S31. Shows the G.R.A.S KEMAR 65 db SPL ISTS input Response (dB SPL) Curve (Right Ear) with Male 70-79 Right Ear Targets (dB SPL).

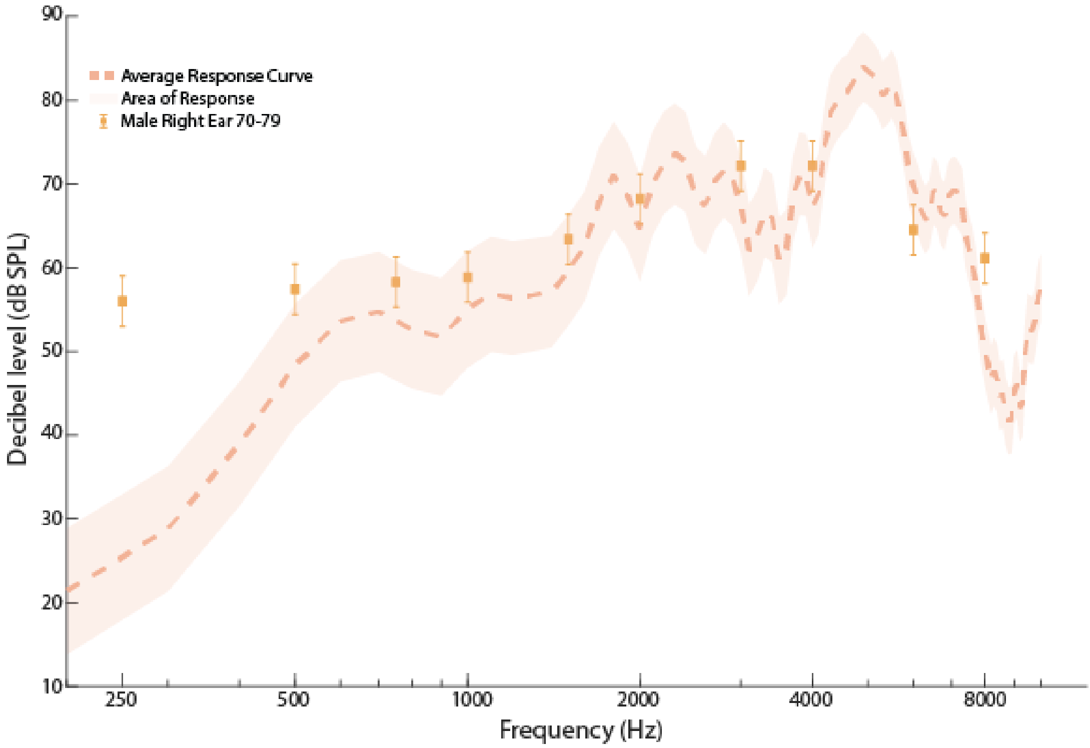


FIG. S32. Shows the G.R.A.S KEMAR 65 db SPL ISTS input Response (dB SPL) Curve (Left Ear) with Male 70-79 Right Ear Targets (dB SPL).

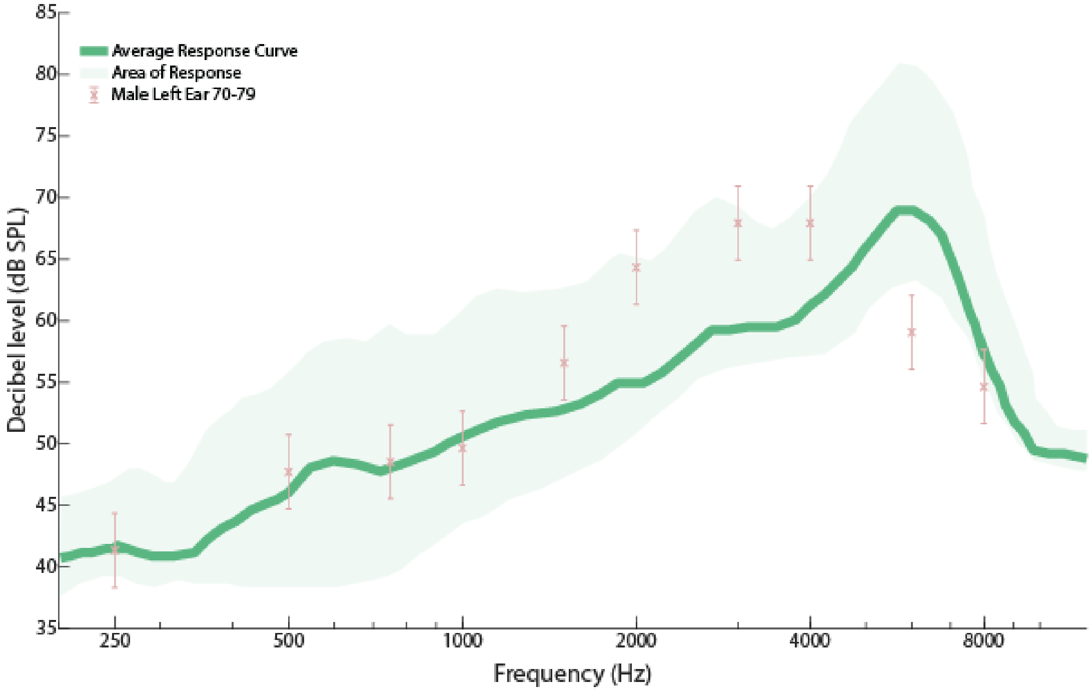


FIG. S33. Shows the ISTS 55 dB SPL Response (dB SPL) Curve with Male 70-79 Left Ear Targets (dB SPL).

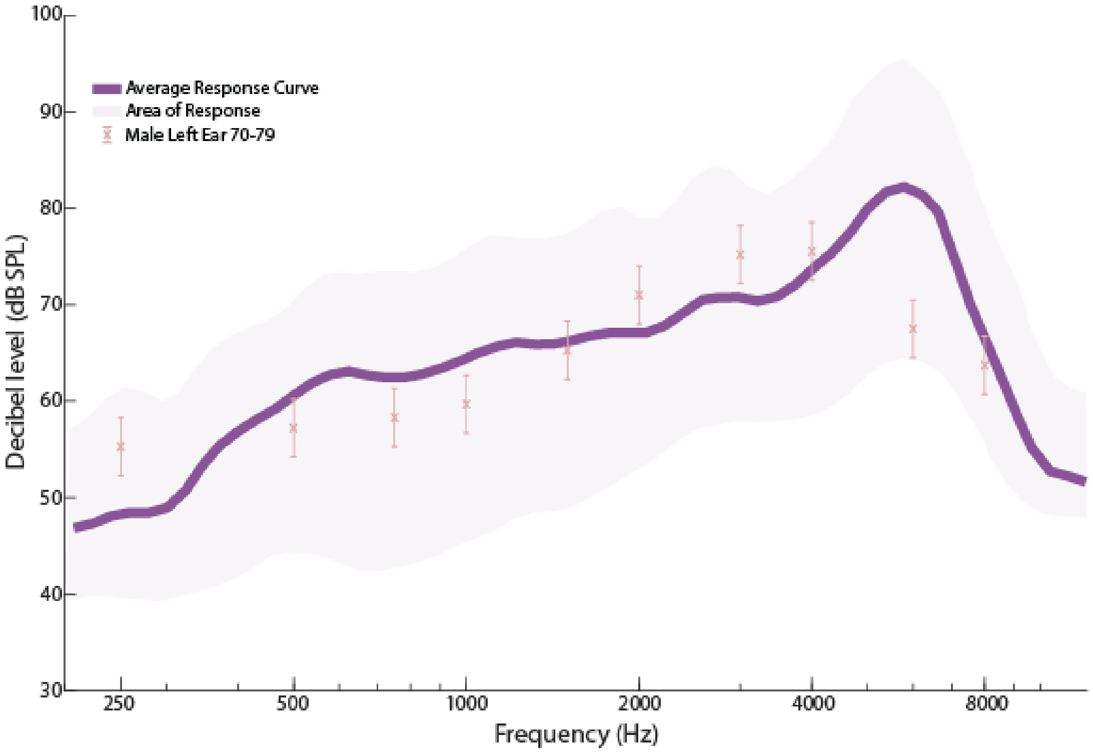


FIG. S34. Shows the ISTS 65 dB SPL Response (dB SPL) Curve with Male 70-79 Left Ear Targets (dB SPL).

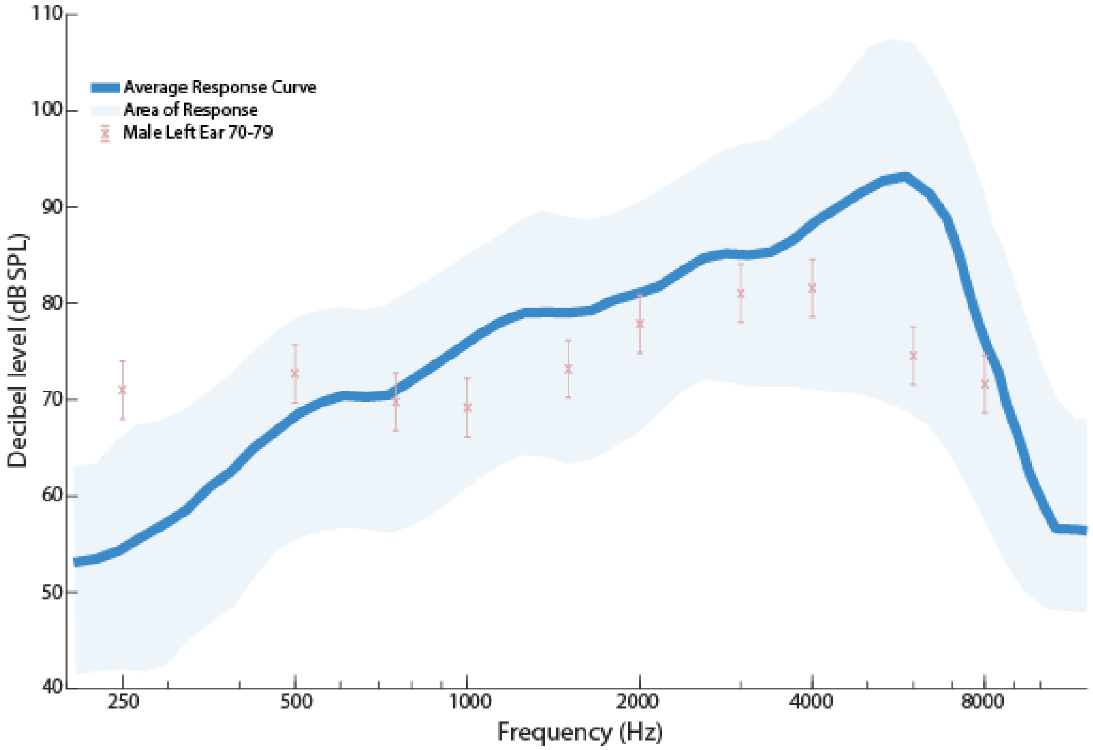


FIG. S35. Shows the ISTS 80 dB SPL Response (dB SPL) Curve with Male 70-79 Left Ear Targets (dB SPL).

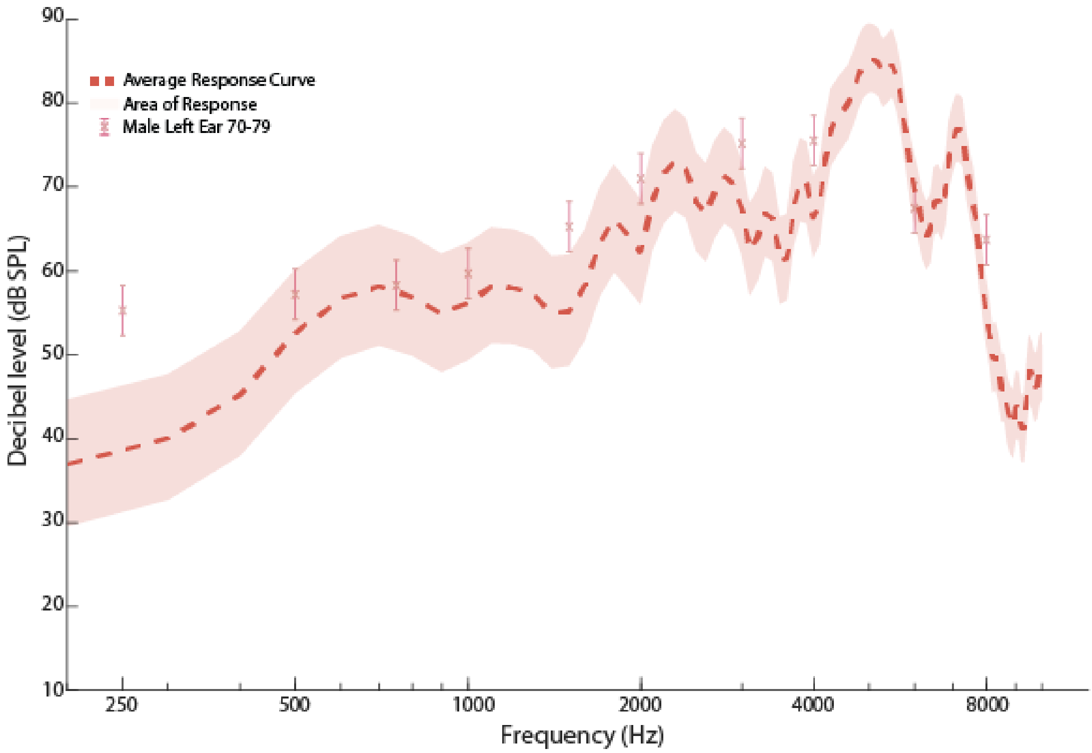


FIG. S36. Shows the G.R.A.S KEMAR 65 db SPL ISTS input Response (dB SPL) Curve (Right Ear) with Male 70-79 Left Ear Targets (dB SPL).

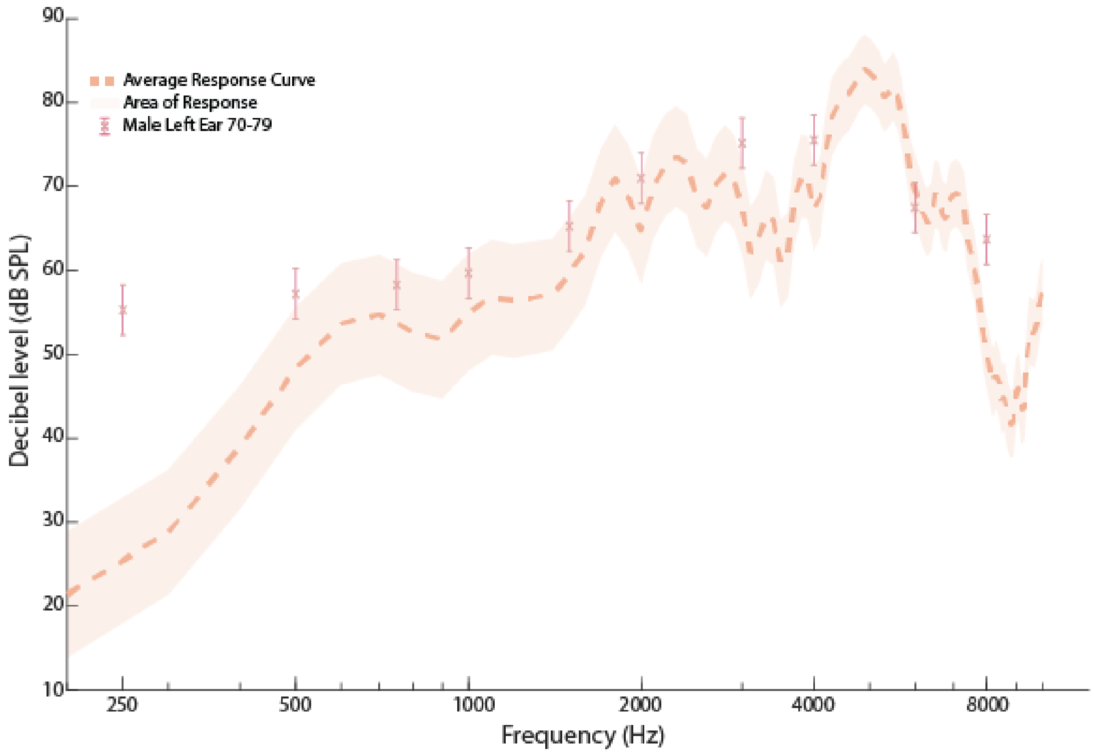


FIG. S37. Shows the G.R.A.S KEMAR 65 db SPL ISTS input Response (dB SPL) Curve (Left Ear) with Male 70-79 Left Ear Targets (dB SPL).

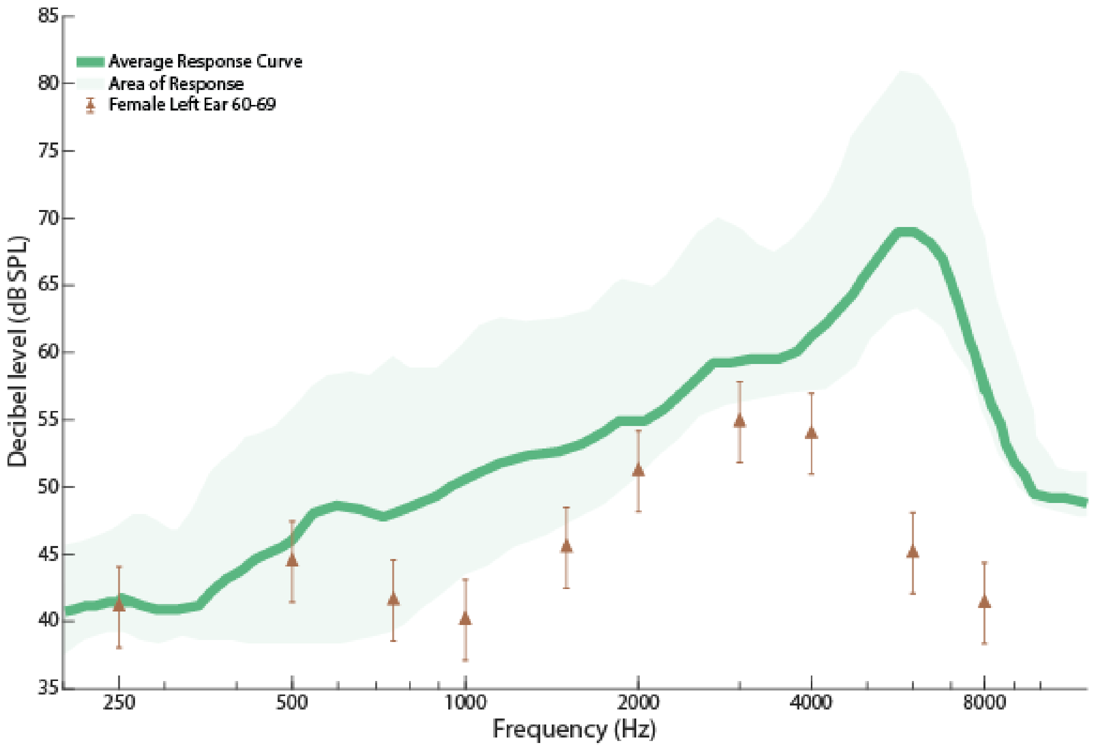


FIG. S38. Shows the ISTS 55 dB SPL Response (dB SPL) Curve with Female 60-69 Left Ear Targets (dB SPL).

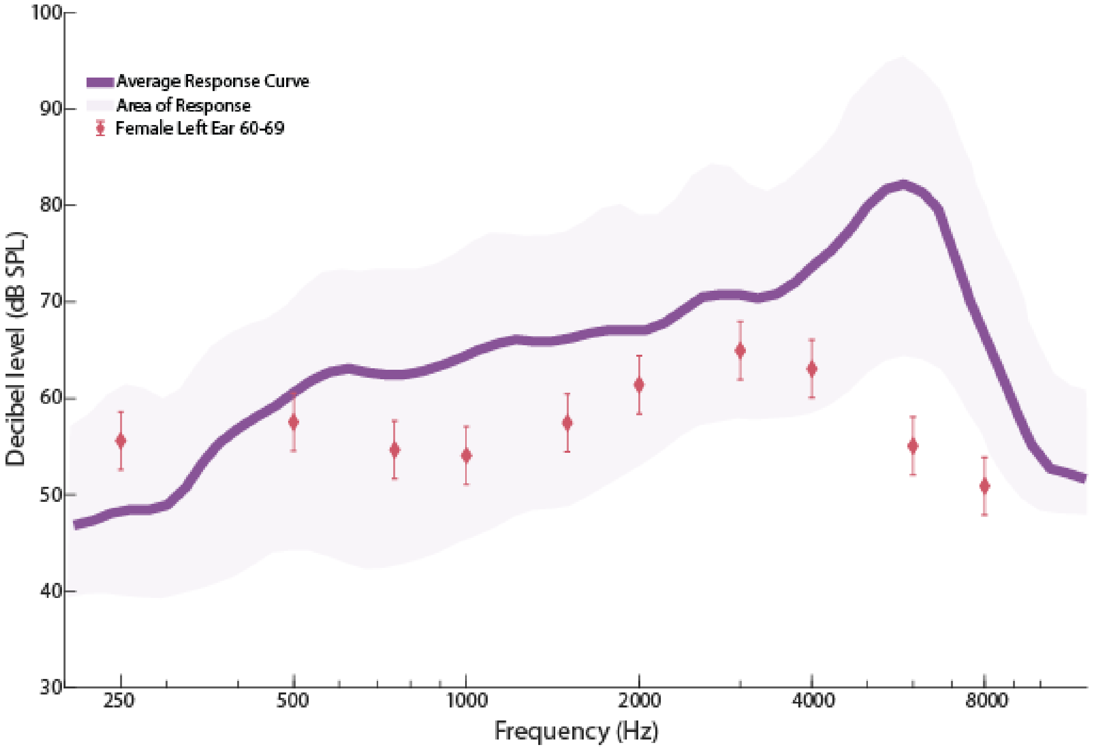


FIG. S39. Shows the ISTS 65 dB SPL Response (dB SPL) Curve with Female 60-69 Left Ear Targets (dB SPL).

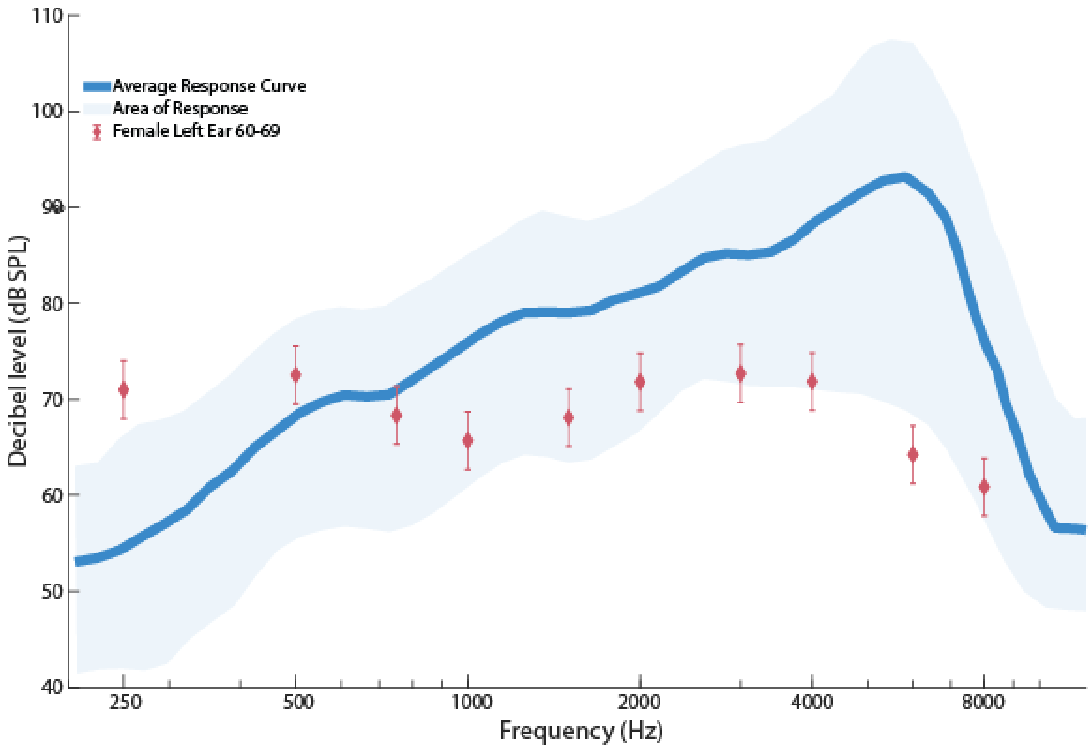


FIG. S40. Shows the ISTS 80 dB SPL Response (dB SPL) Curve with Female 60-69 Left Ear Targets (dB SPL).

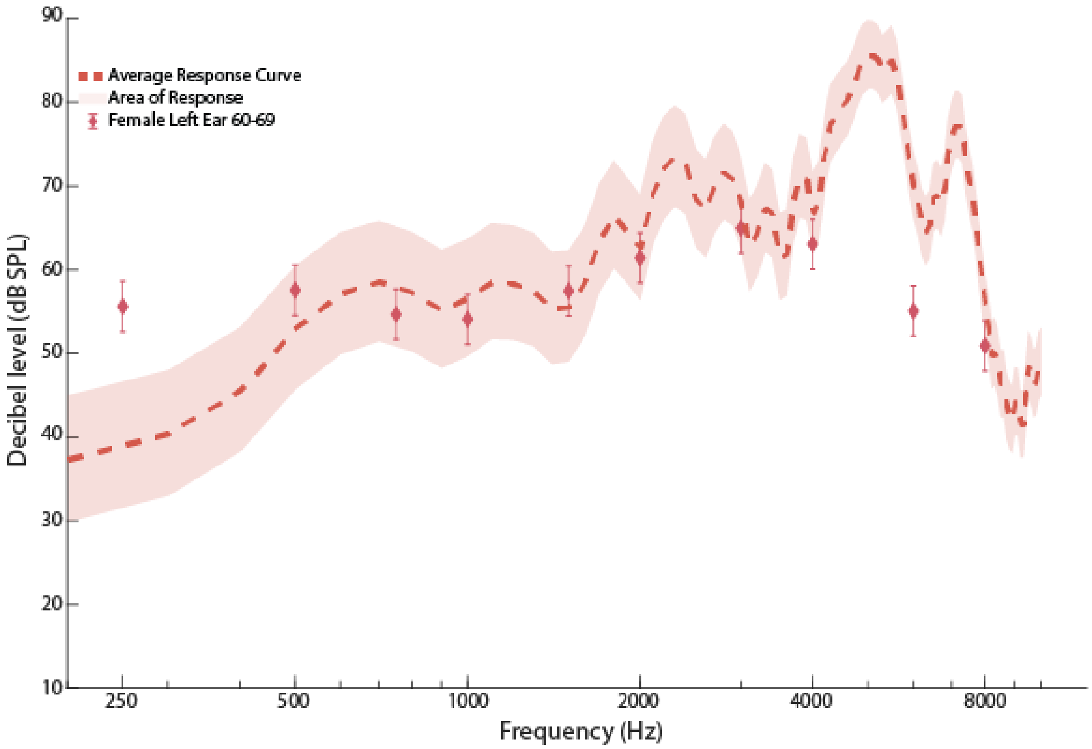


FIG. S41. Shows the G.R.A.S KEMAR 65 db SPL ISTS input Response (dB SPL) Curve (Right Ear) with Female 60-69 Left Ear Targets (dB SPL).

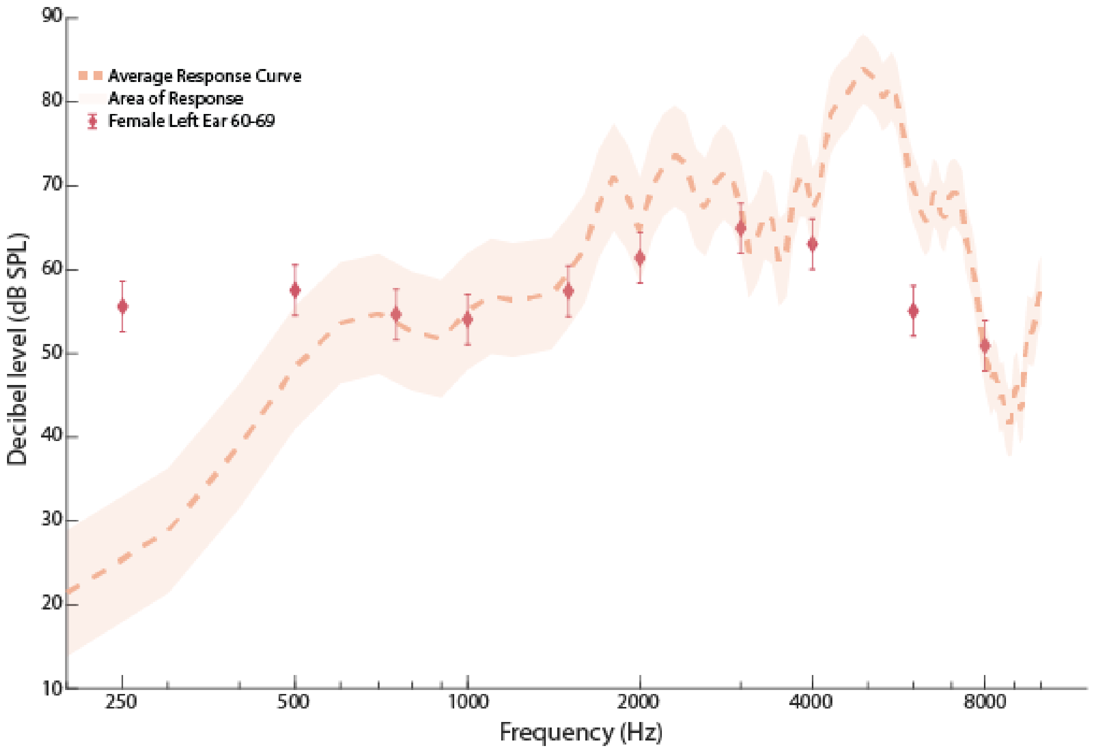


FIG. S42. Shows the G.R.A.S KEMAR 65 db SPL ISTS input Response (dB SPL) Curve (Left Ear) with Female 60-69 Left Ear Targets (dB SPL).

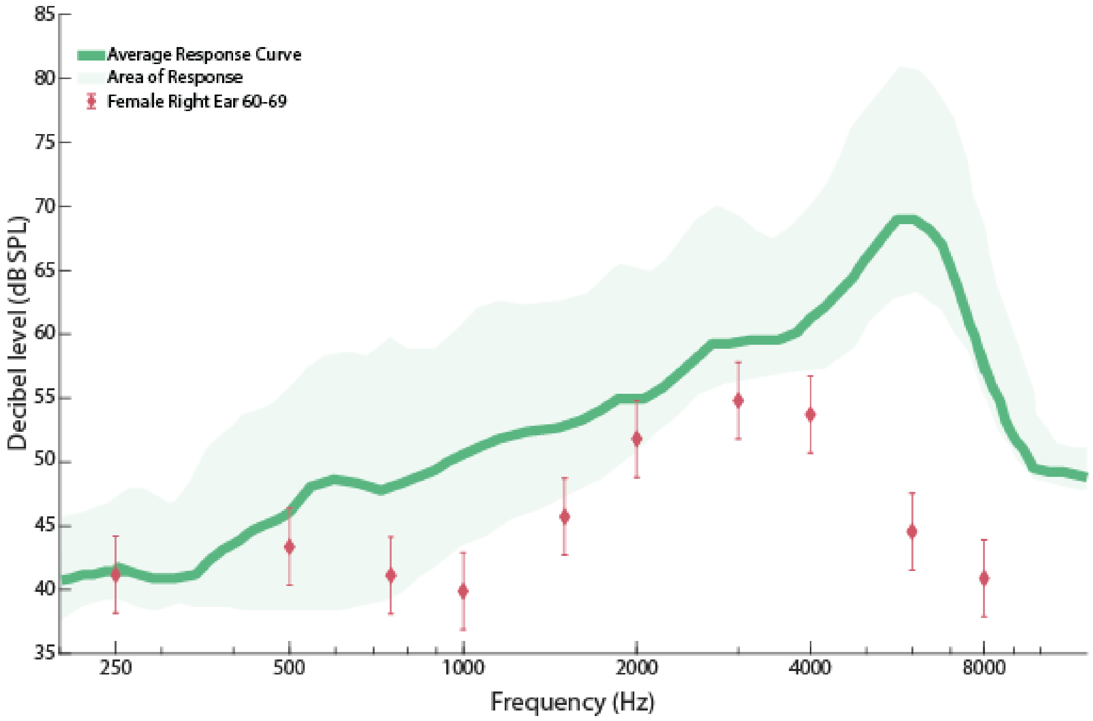


FIG. S43. Shows the ISTS 55 dB SPL Response (dB SPL) Curve with Female 60-69 Right Ear Targets (dB SPL).

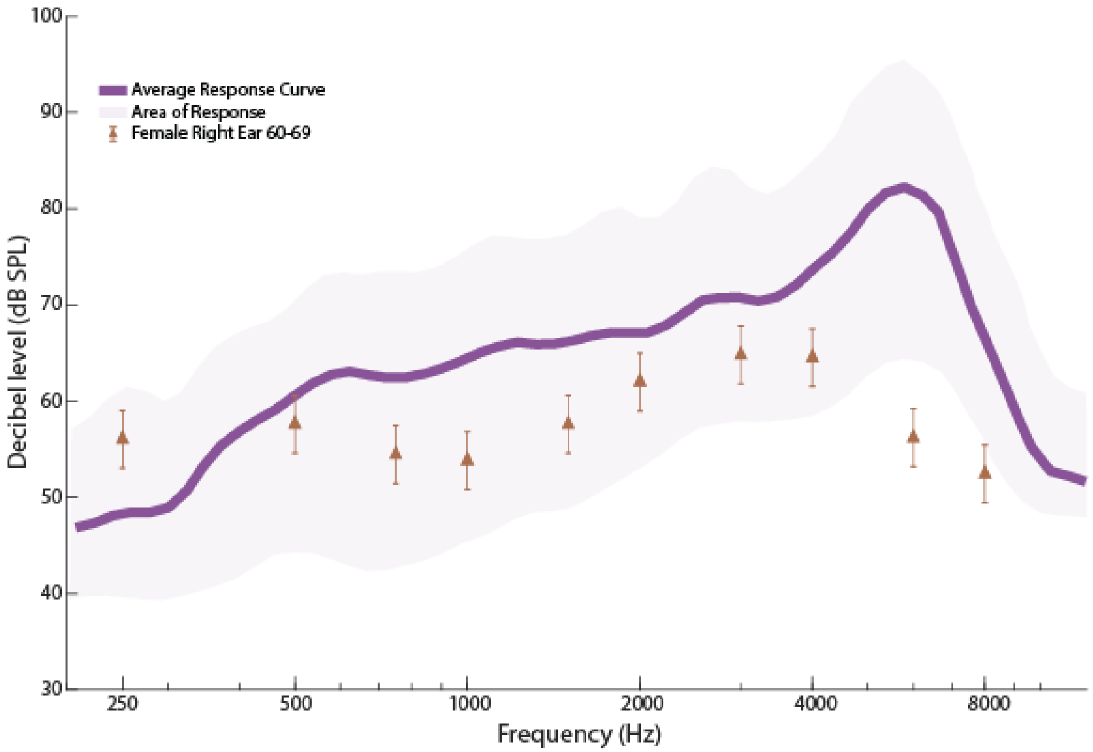


FIG. S44. Shows the ISTS 65 dB SPL Response (dB SPL) Curve with Female 60-69 Right Ear Targets (dB SPL).

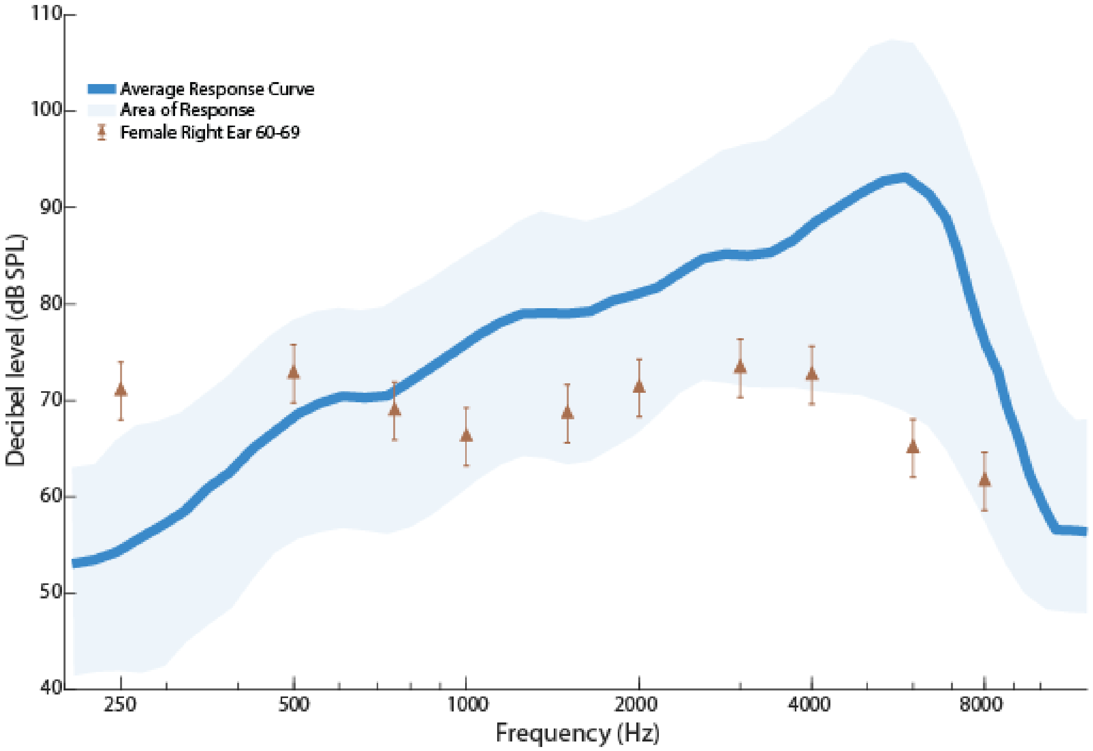


FIG. S45. Shows the ISTS 80 dB SPL Response (dB SPL) Curve with Female 60-69 Right Ear Targets (dB SPL).

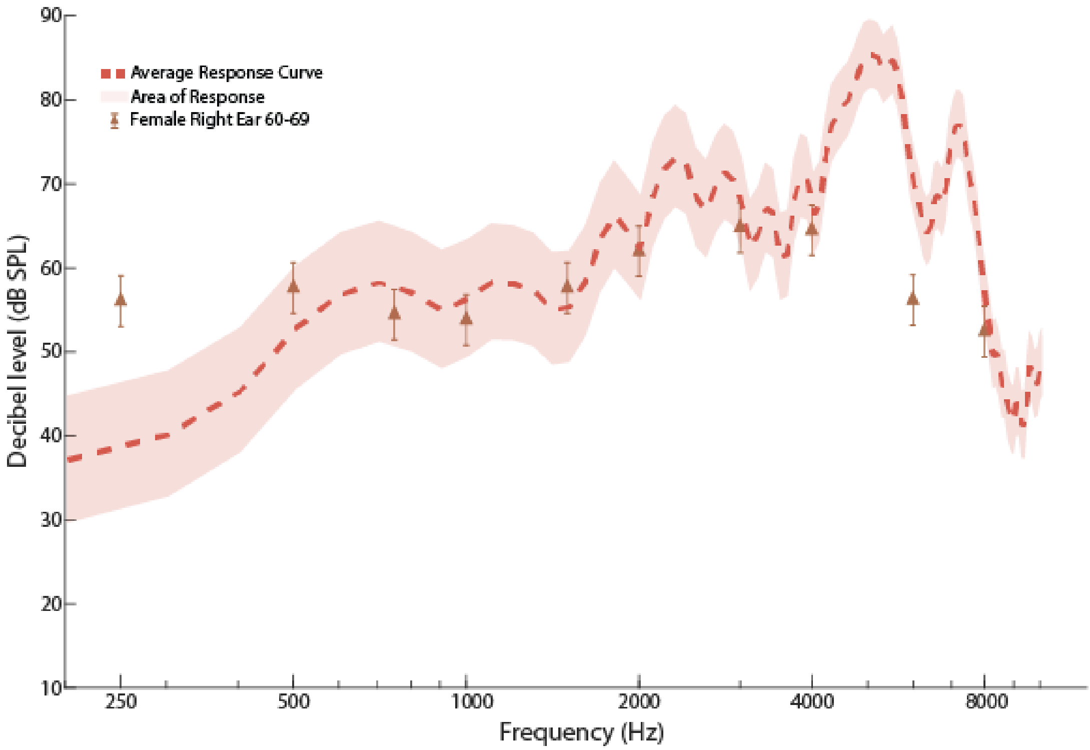


FIG. S46. Shows the G.R.A.S KEMAR 65 db SPL ISTS input Response (dB SPL) Curve (Right Ear) with Female 60-69 Right Ear Targets (dB SPL).

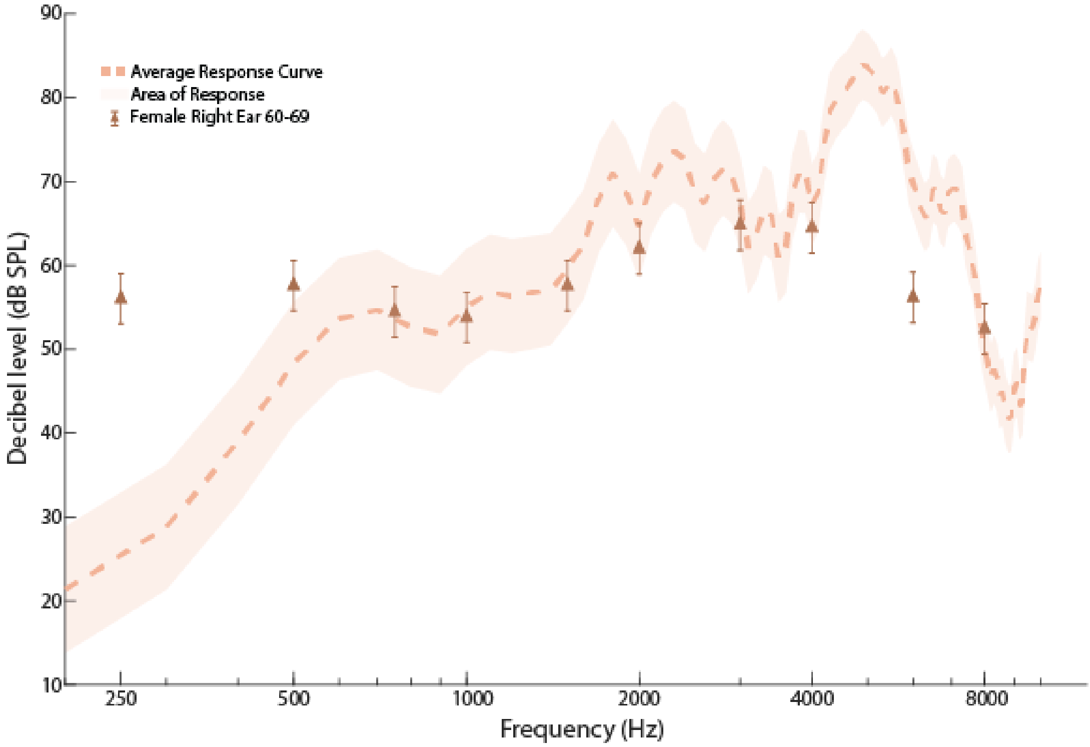


FIG. S47. Shows the G.R.A.S KEMAR 65 db SPL ISTS input Response (dB SPL) Curve (Left Ear) with Female 60-69 Right Ear Targets (dB SPL).

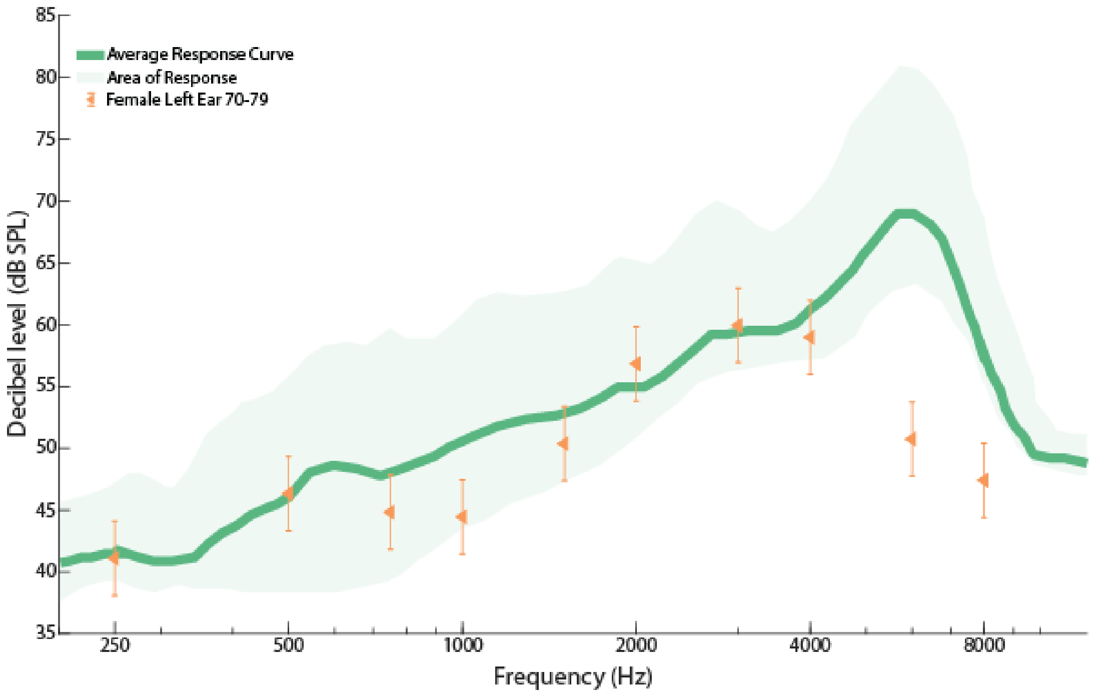


FIG. S48. Shows the ISTS 55 dB SPL Response (dB SPL) Curve with Female 70-79 Left Ear Targets (dB SPL).

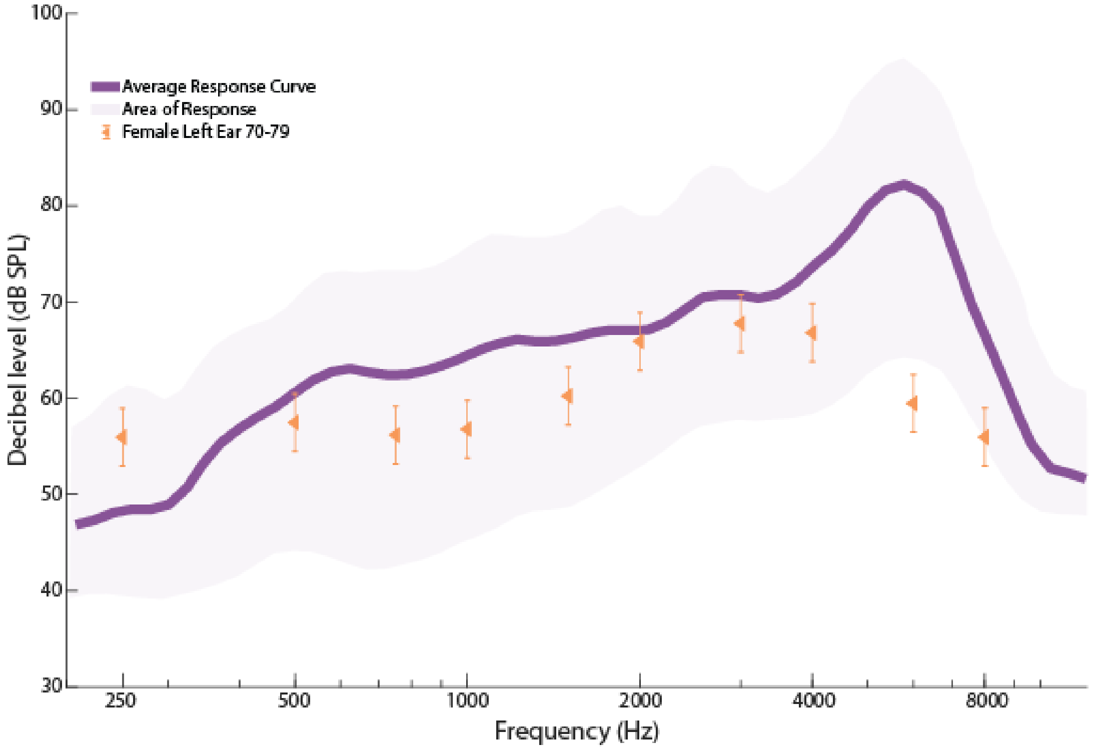


FIG. S49. Shows the ISTS 65 dB SPL Response (dB SPL) Curve with Female 70-79 Left Ear Targets (dB SPL).

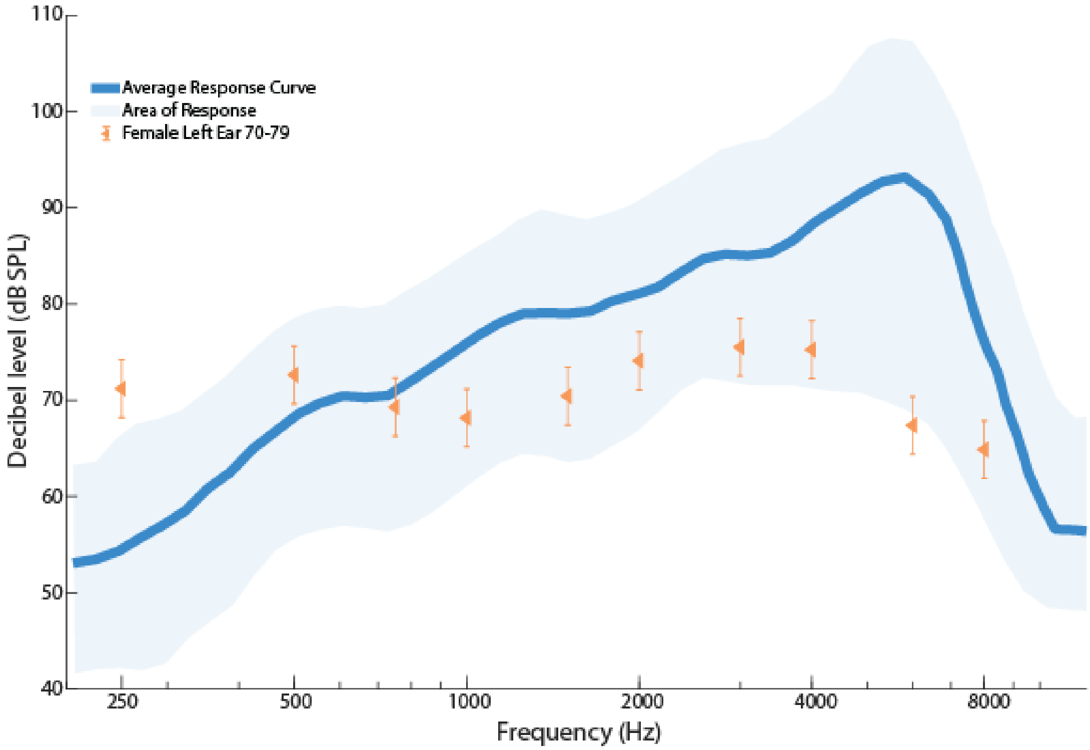


FIG. S50. Shows the ISTS 80 dB SPL Response (dB SPL) Curve with Female 70-79 Left Ear Targets (dB SPL).

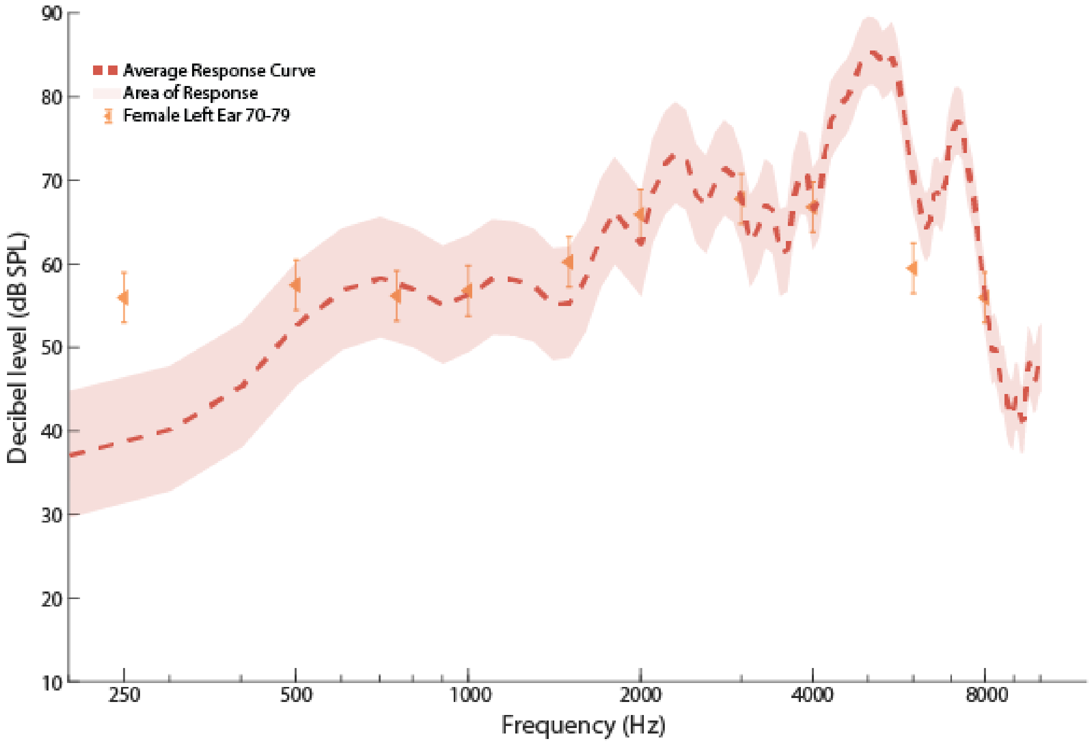


FIG. S51. Shows the G.R.A.S KEMAR 65 db SPL ISTS input Response (dB SPL) Curve (Right Ear) with Female 70-79 Left Ear Targets (dB SPL).

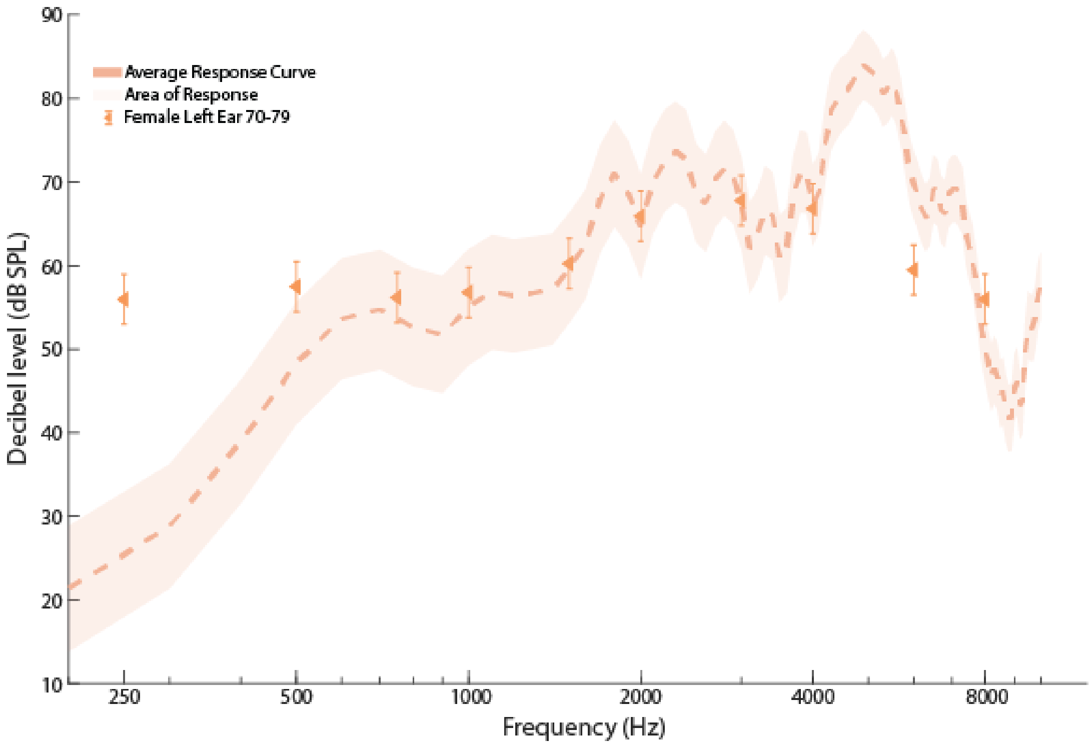


FIG. S52. Shows the G.R.A.S KEMAR 65 db SPL ISTS input Response (dB SPL) Curve (Left Ear) with Female 70-79 Left Ear Targets (dB SPL).

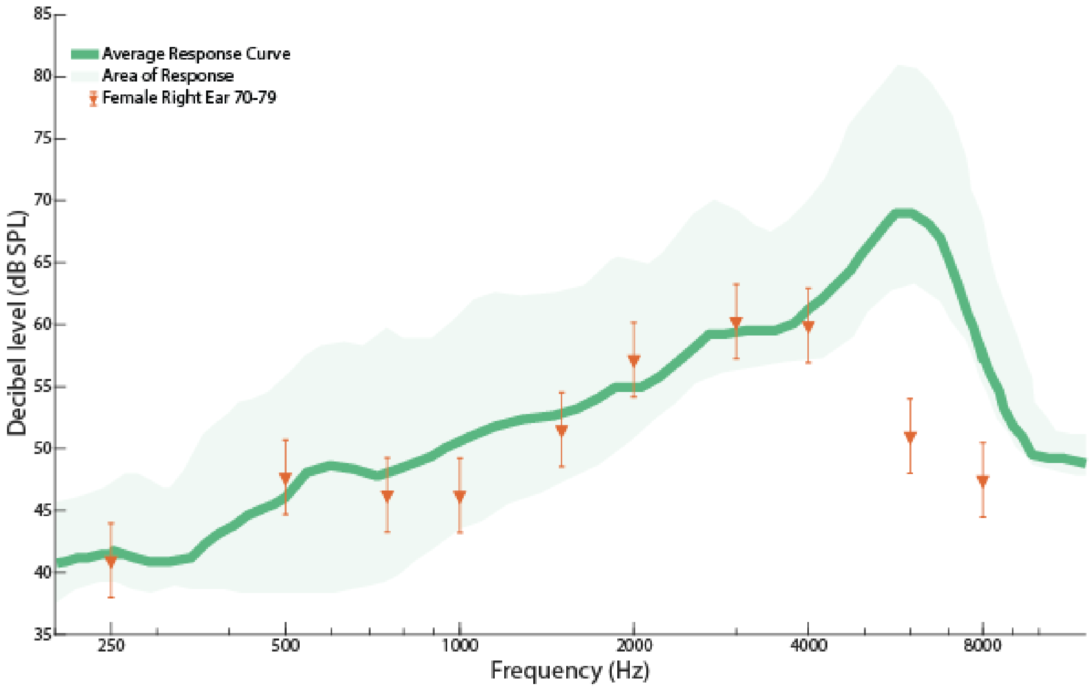


FIG. S53. Shows the ISTS 55 dB SPL Response (dB SPL) Curve with Male 70-79 Right Ear Targets (dB SPL).

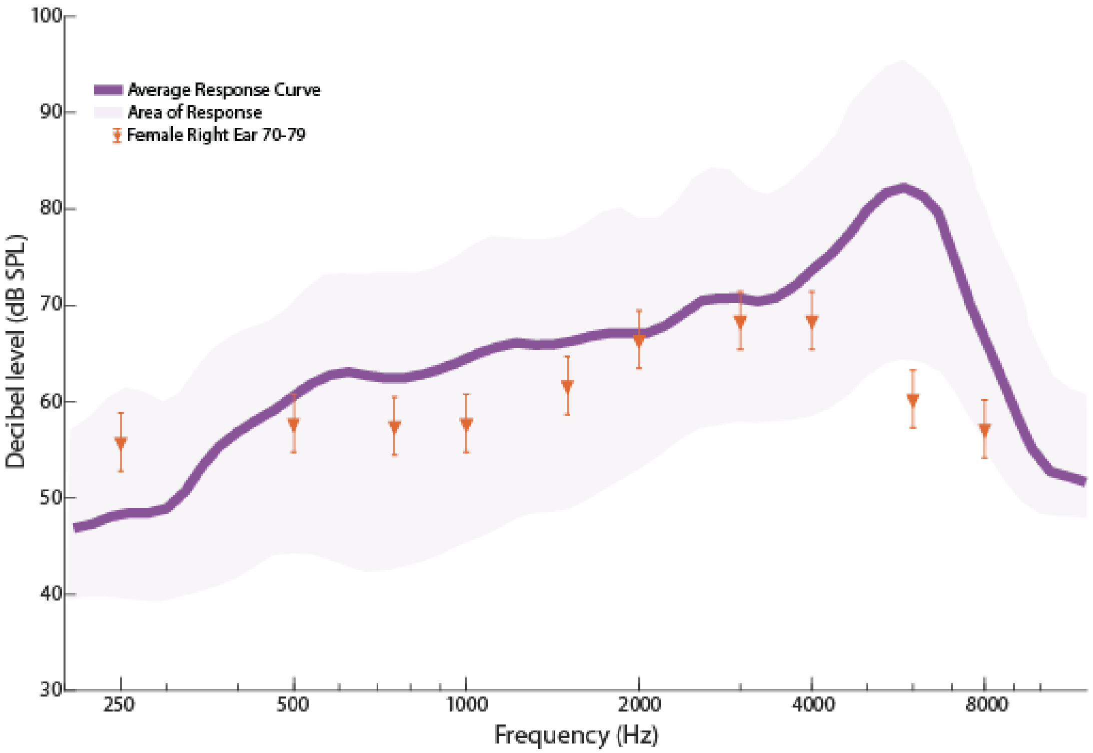


FIG. S54. Shows the ISTS 65 dB SPL Response (dB SPL) Curve with Female 70-79 Right Ear Targets (dB SPL).

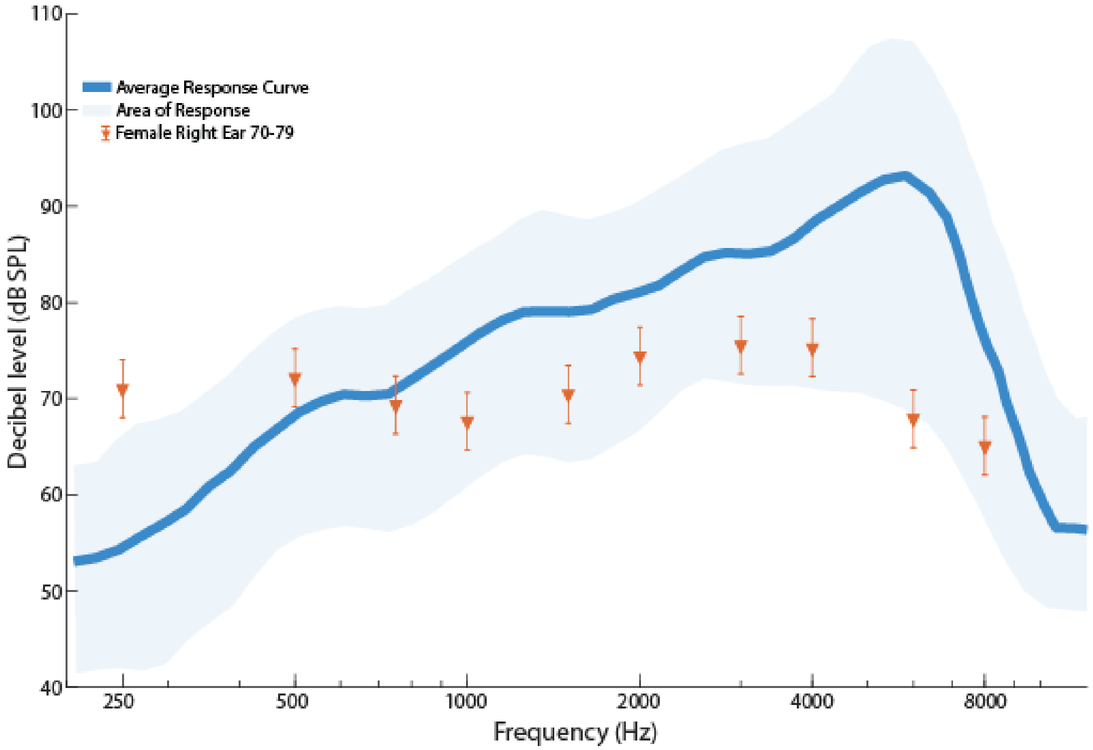


FIG. S55. Shows the ISTS 80 dB SPL Response (dB SPL) Curve with Female 70-79 Right Ear Targets (dB SPL).

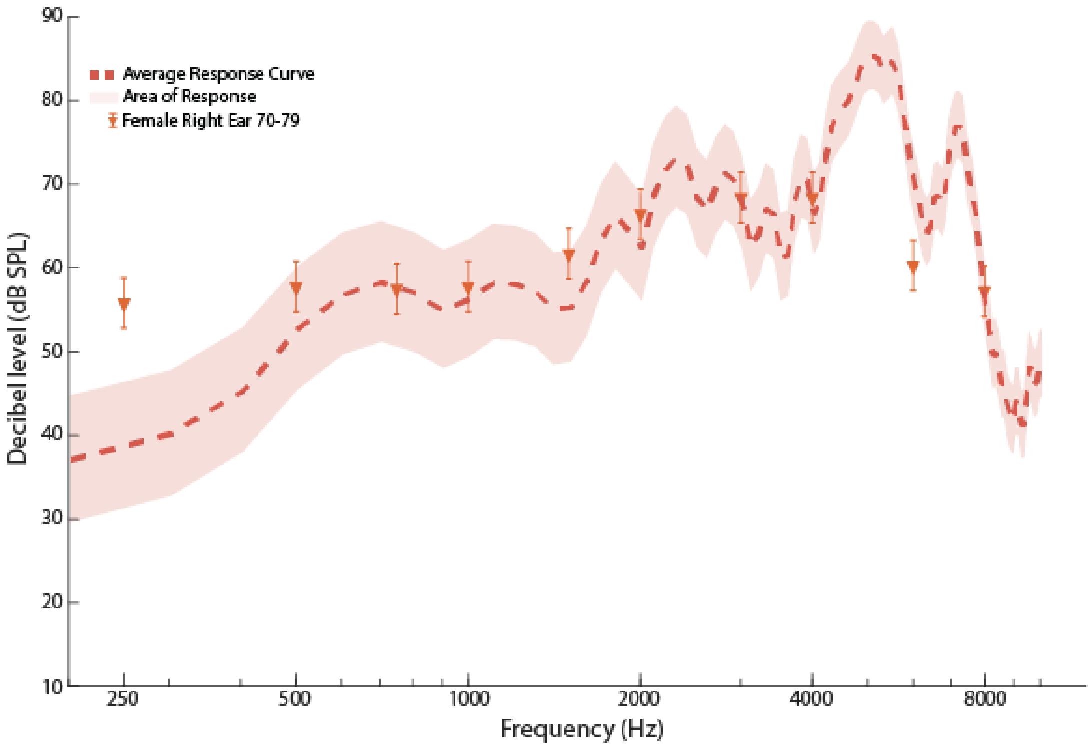


FIG. S56. Shows the G.R.A.S KEMAR 65 db SPL ISTS input Response (dB SPL) Curve (Right Ear) with Female 70-79 Right Ear Targets (dB SPL).

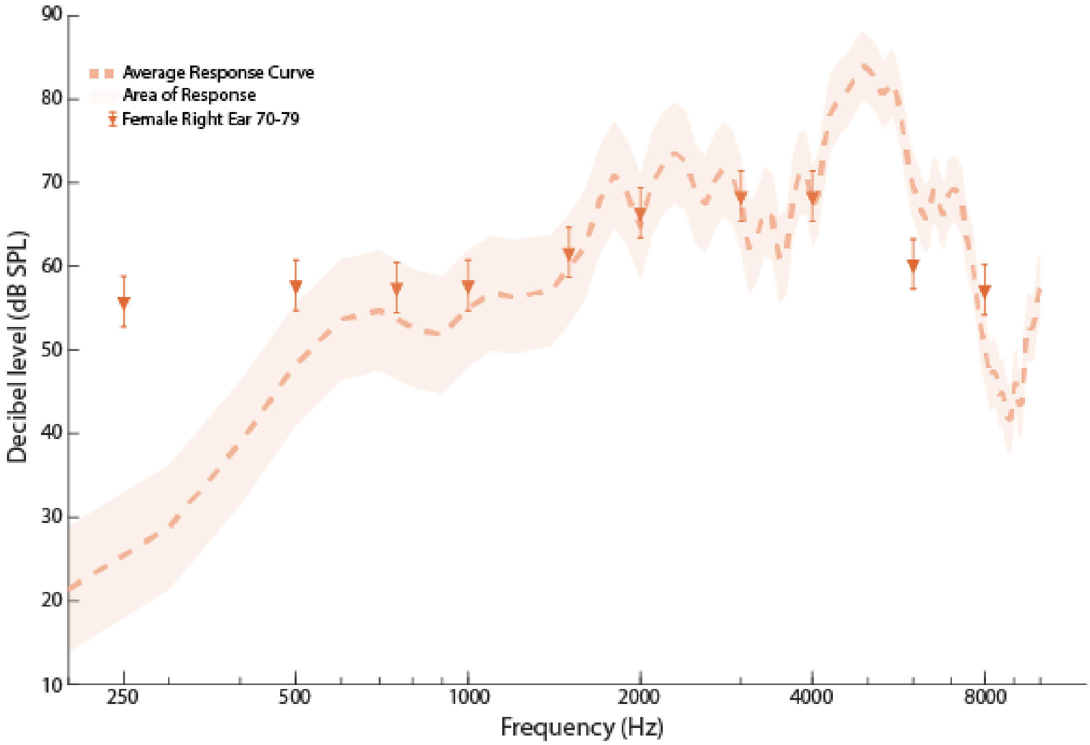


FIG. S57. Shows the G.R.A.S KEMAR 65 db SPL ISTS input Response (dB SPL) Curve (Left Ear) with Female 70-79 Right Ear Targets (dB SPL).

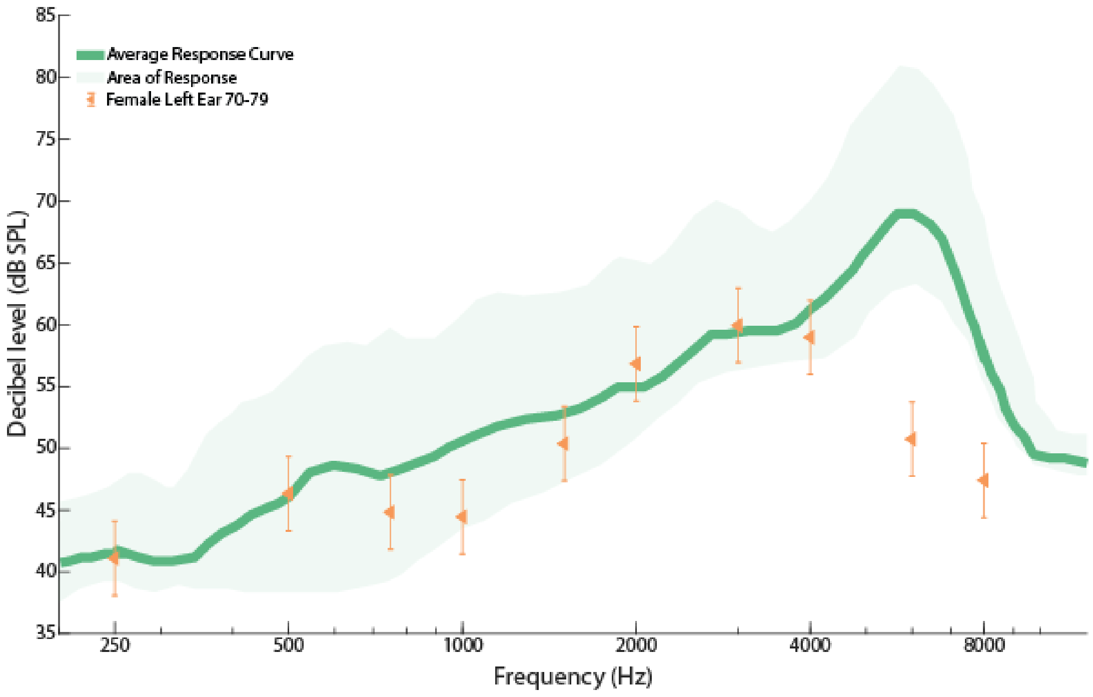


FIG. S58. Shows the ISTS 55 dB SPL Response (dB SPL) Curve with Female 70-79 Left Ear Targets (dB SPL).

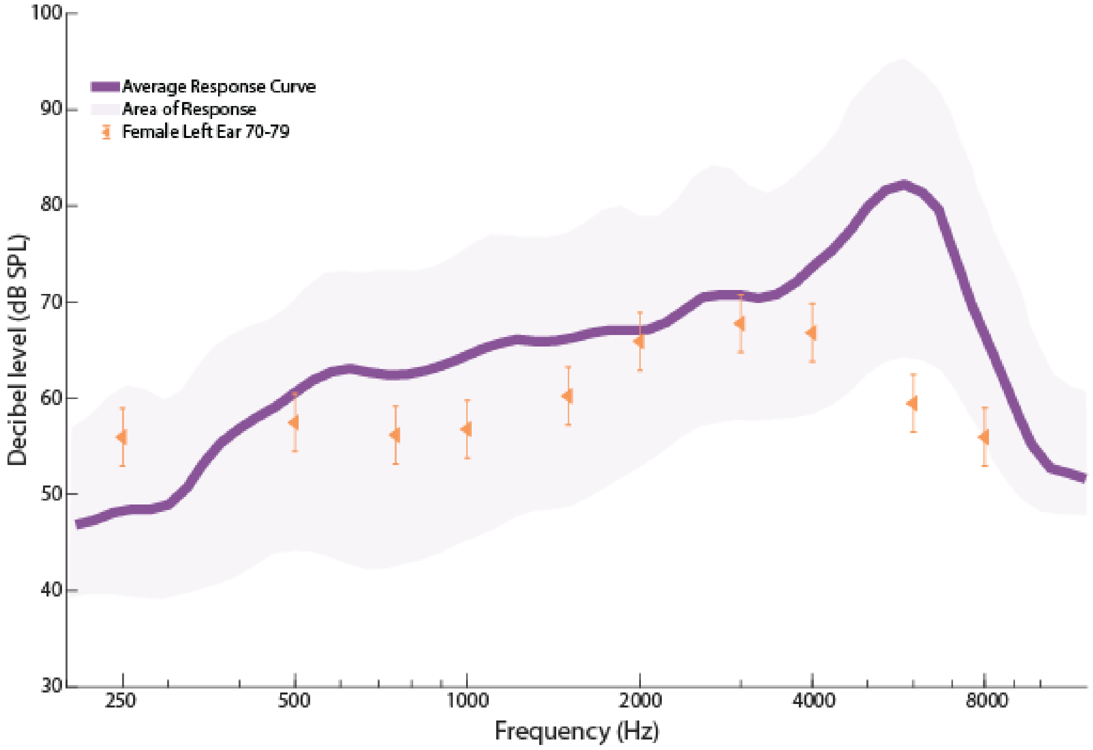


FIG. S59. Shows the ISTS 65 dB SPL Response (dB SPL) Curve with Female 70-79 Left Ear Targets (dB SPL).

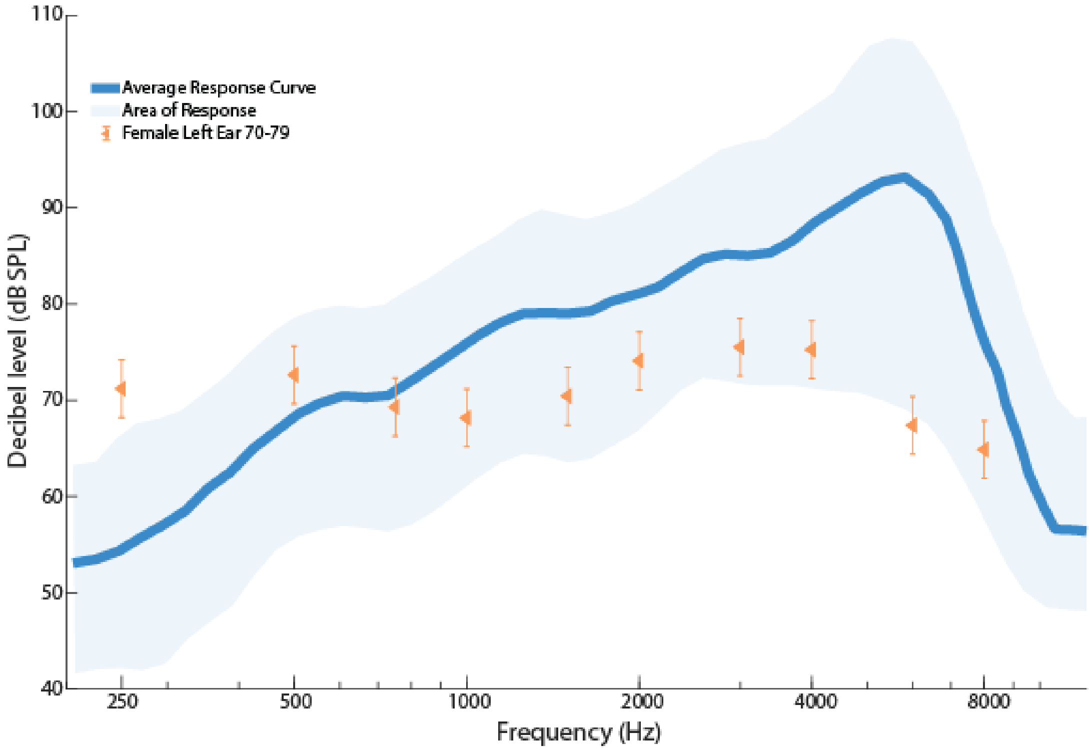


FIG. S60. Shows the ISTS 80 dB SPL Response (dB SPL) Curve with Female 70-79 Left Ear Targets (dB SPL).

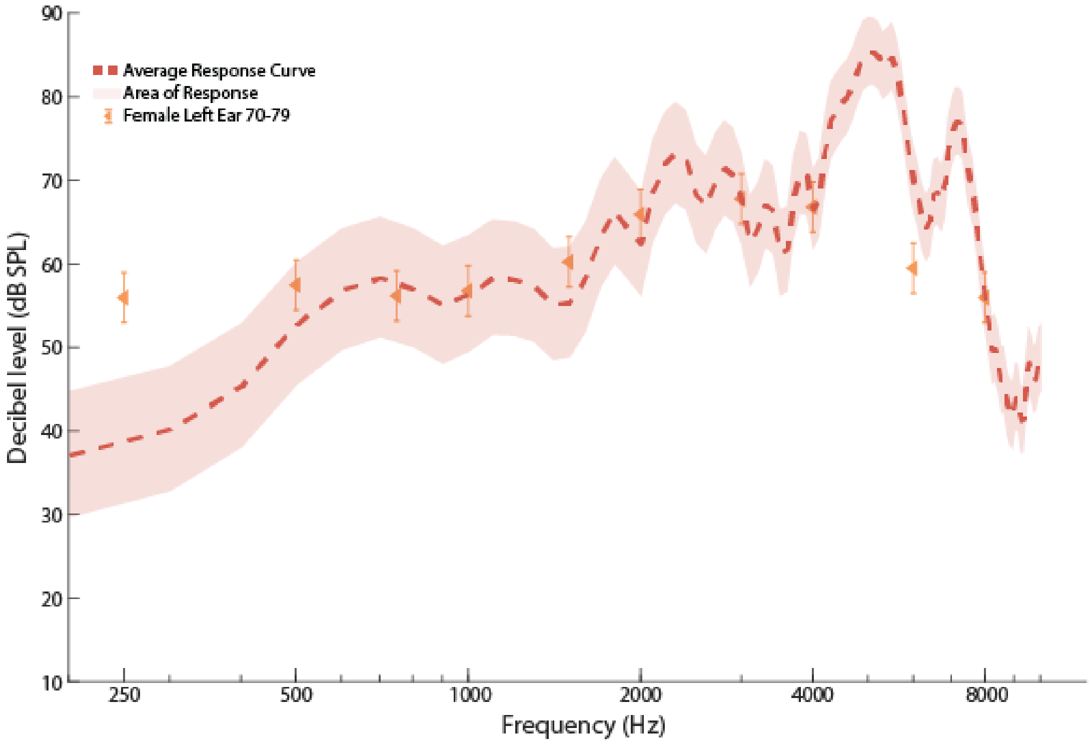


FIG. S61. Shows the G.R.A.S KEMAR 65 db SPL ISTS input Response (dB SPL) Curve (Right Ear) with Female 70-79 Left Ear Targets (dB SPL).

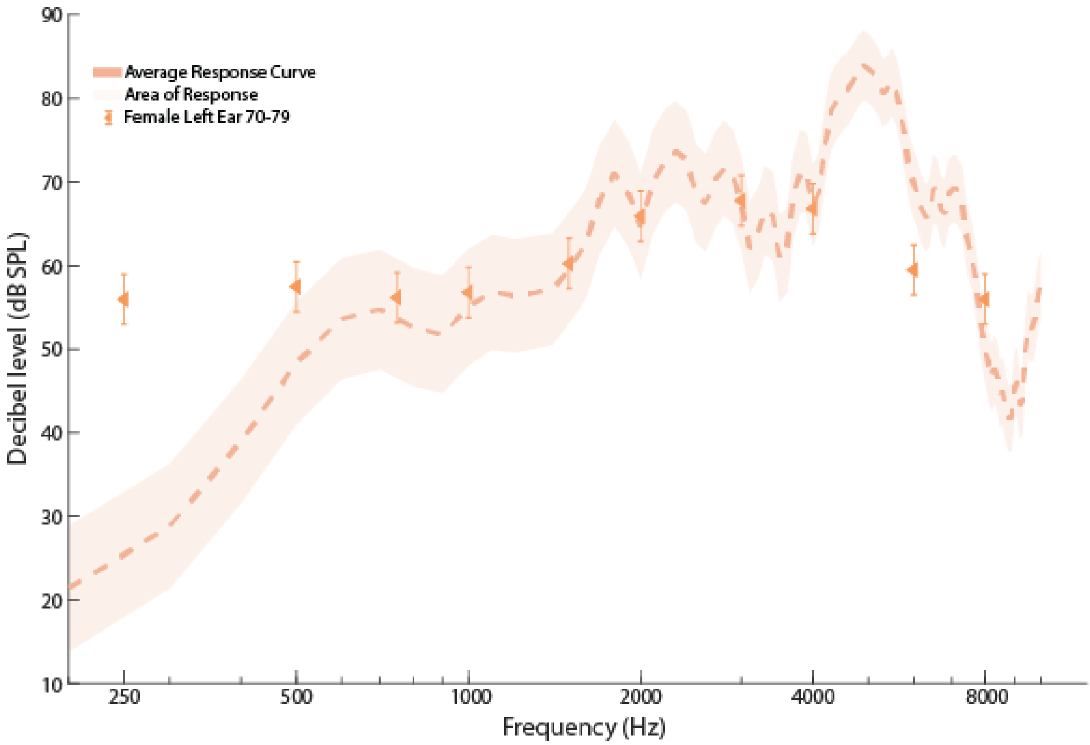


FIG. S62. Shows the G.R.A.S KEMAR 65 db SPL ISTS input Response (dB SPL) Curve (Left Ear) with Female 70-79 Left Ear Targets (dB SPL).

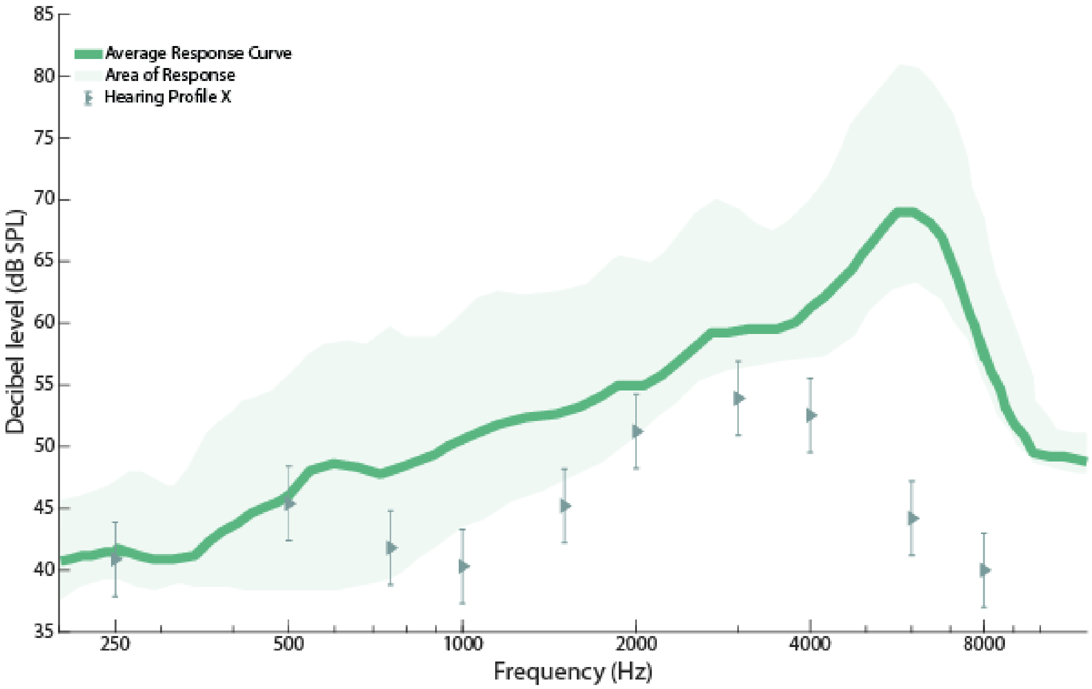


FIG. S63. Shows the ISTS 55 dB SPL Response (dB SPL) Curve with Hearing Profile X Targets (dB SPL).

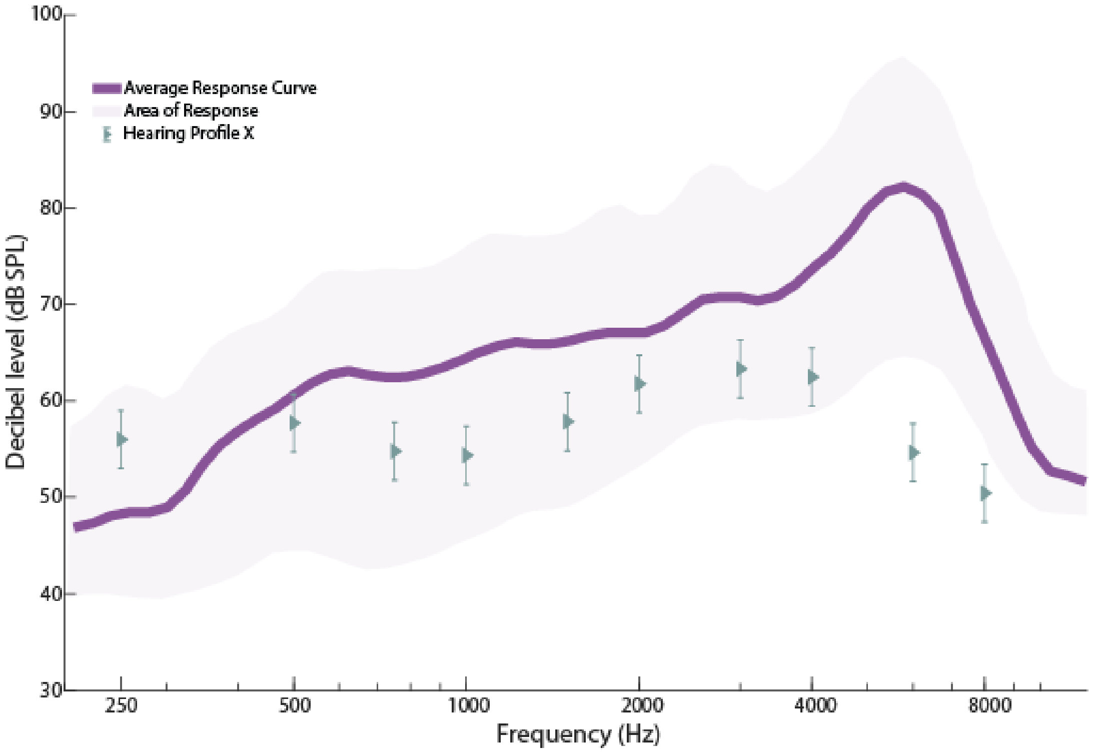


FIG. S64. Shows the ISTS 65 dB SPL Response (dB SPL) Curve with Hearing Profile X Targets (dB SPL).

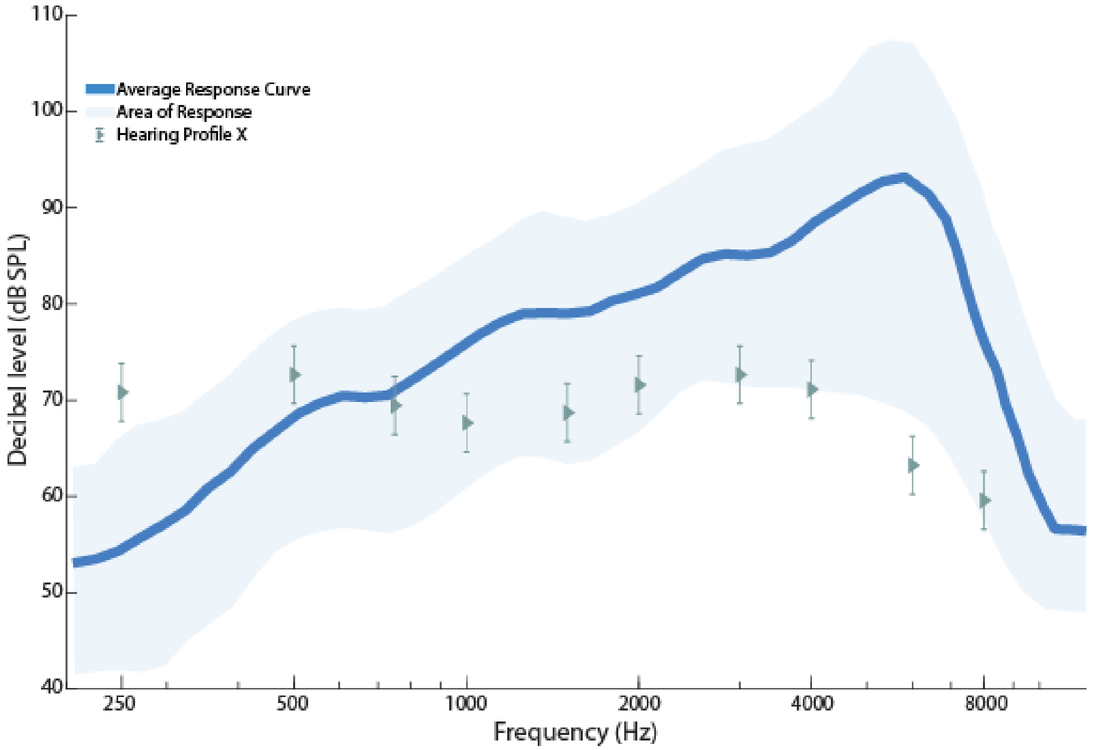


FIG. S65. Shows the ISTS 80 dB SPL Response (dB SPL) Curve with Hearing Profile X Targets (dB SPL).

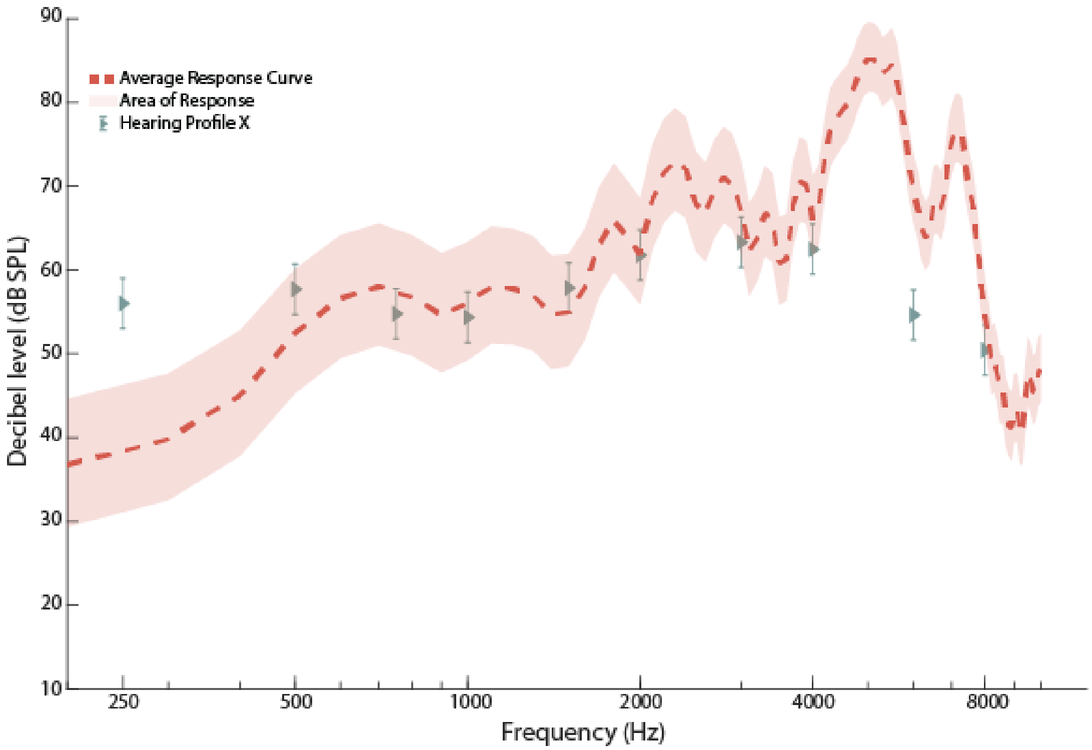


FIG. S66. Shows the G.R.A.S KEMAR 65 db SPL ISTS input Response (dB SPL) Curve (Right Ear) with Hearing Profile X Targets (dB SPL).

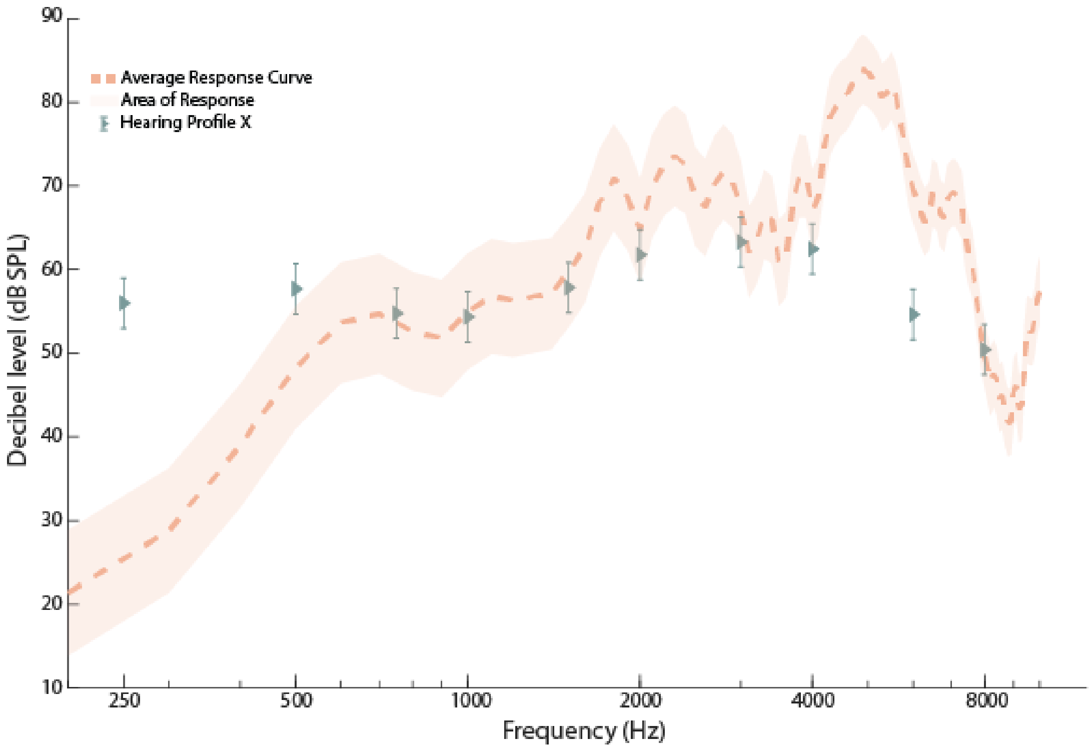


FIG. S67. Shows the G.R.A.S KEMAR 65 db SPL ISTS input Response (dB SPL) Curve (Left Ear) with Hearing Profile X Targets (dB SPL).

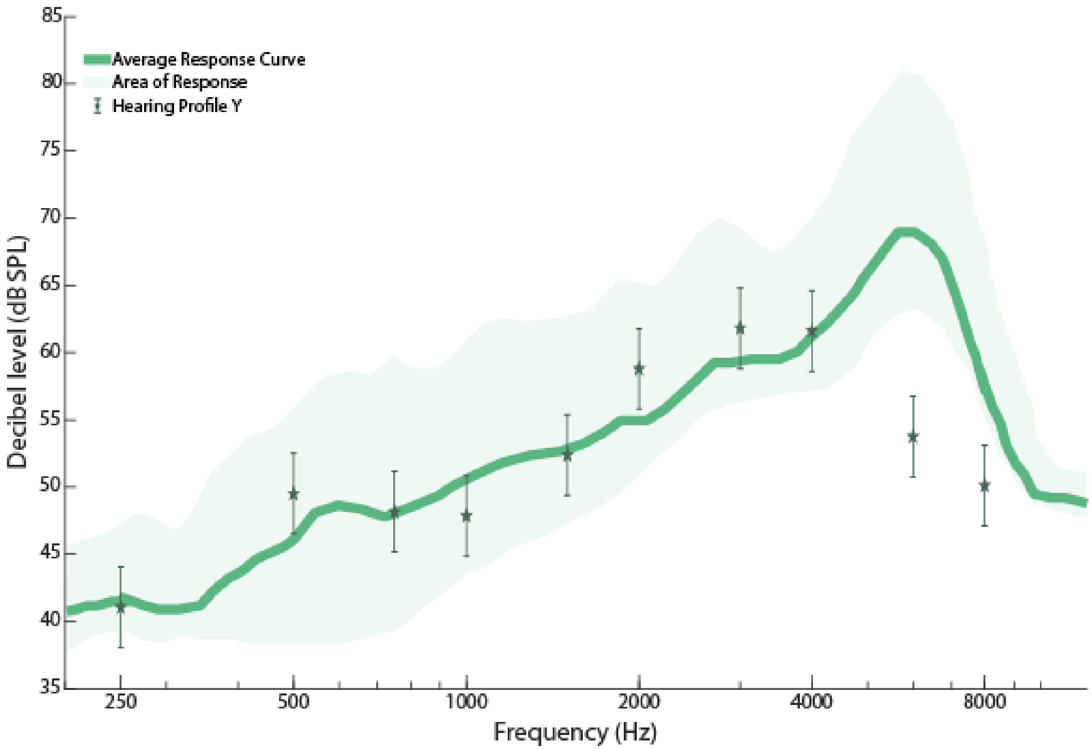


FIG. S68. Shows the ISTS 55 dB SPL Response (dB SPL) Curve with Hearing Profile Y Targets (dB SPL).

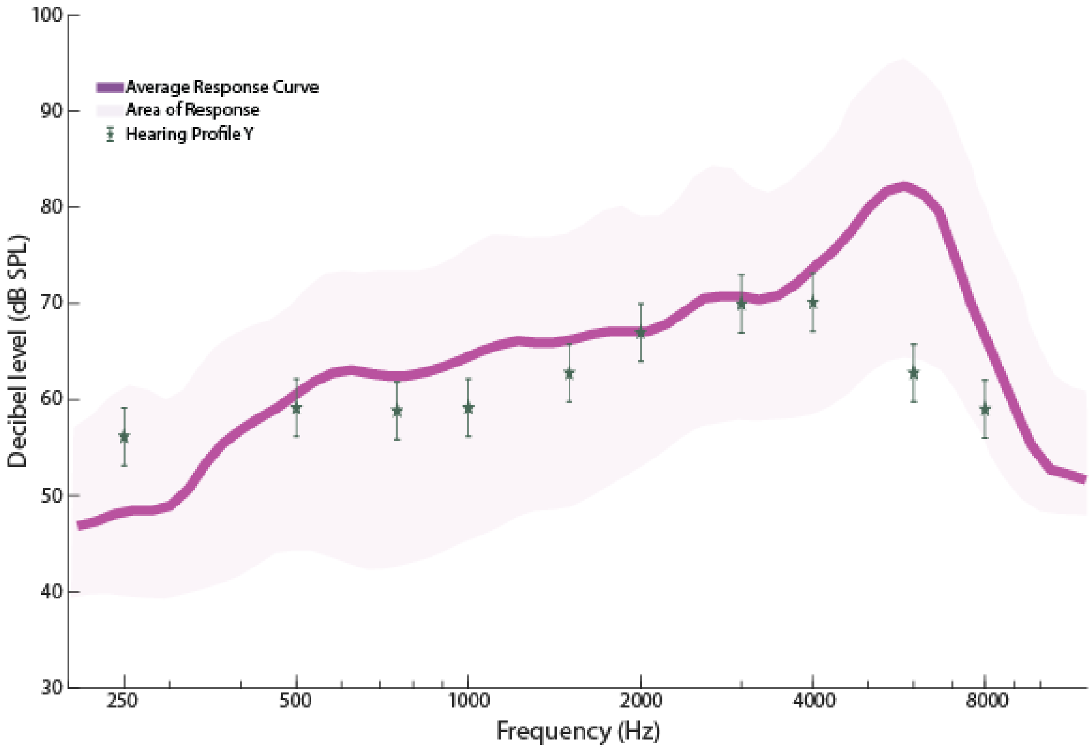


FIG. S69. Shows the ISTS 65 dB SPL Response (dB SPL) Curve with Hearing Profile Y Targets (dB SPL).

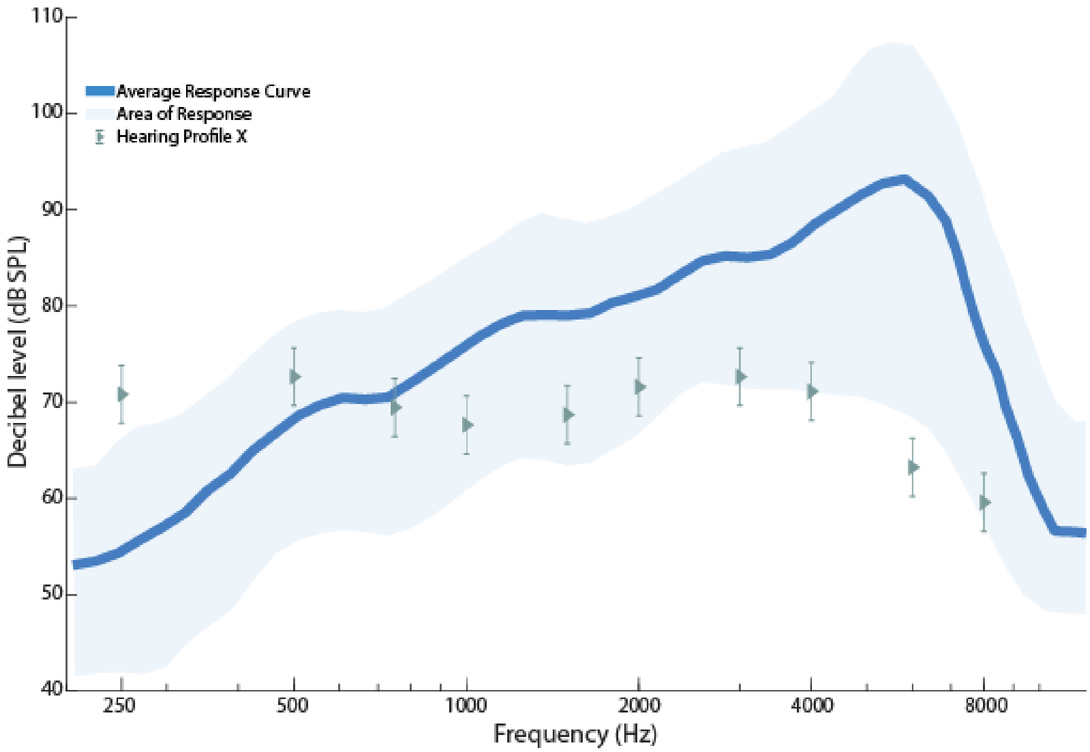


FIG. S70. Shows the ISTS 80 dB SPL Response (dB SPL) Curve with Hearing Profile Y Targets (dB SPL).

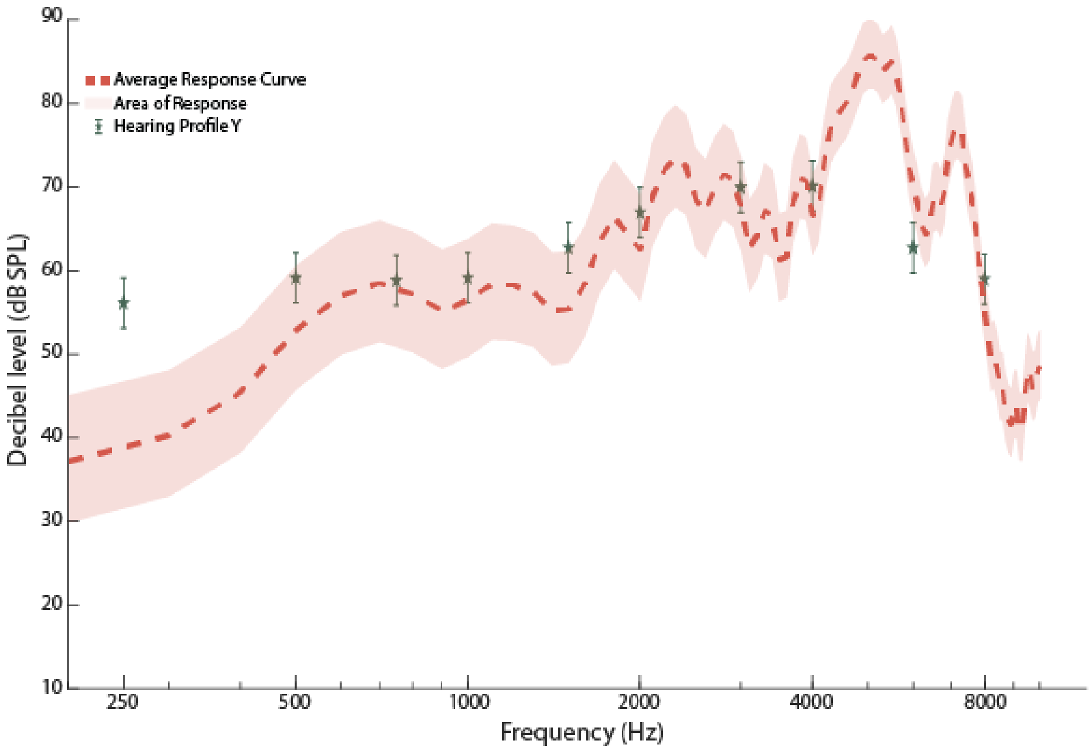


FIG. S71. Shows the G.R.A.S KEMAR 65 db SPL ISTS input Response (dB SPL) Curve (Right Ear) with Hearing Profile Y Targets (dB SPL).

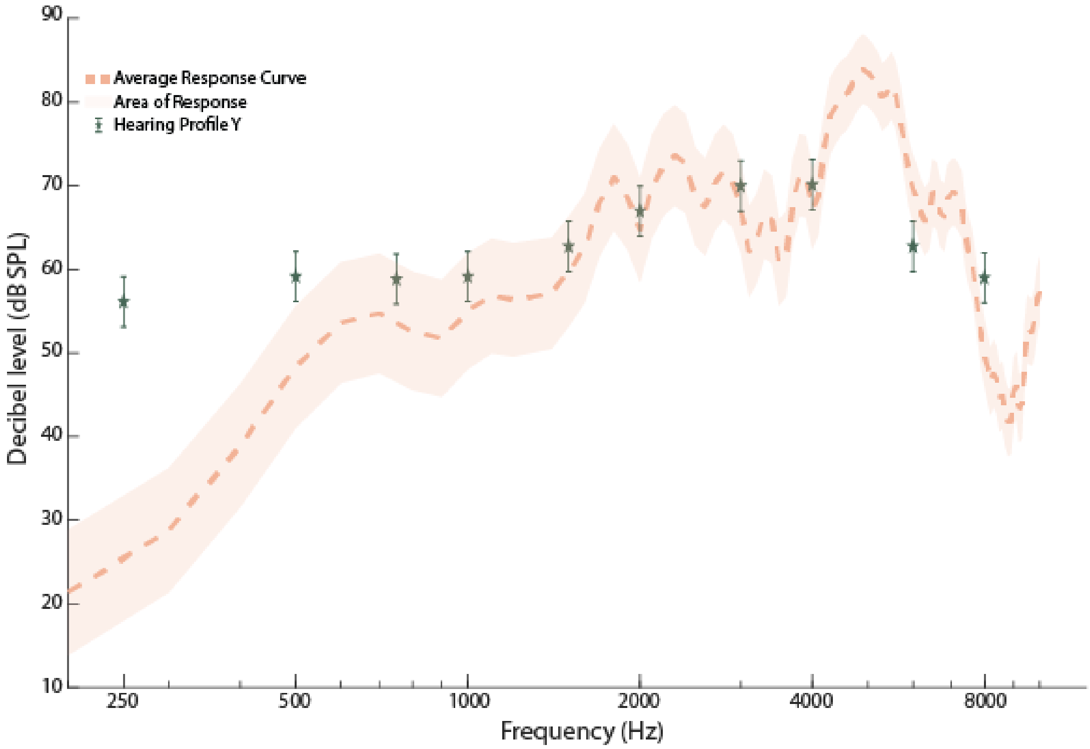


FIG. S72. Shows the G.R.A.S KEMAR 65 db SPL ISTS input Response (dB SPL) Curve (Left Ear) with Hearing Profile Y Targets (dB SPL).

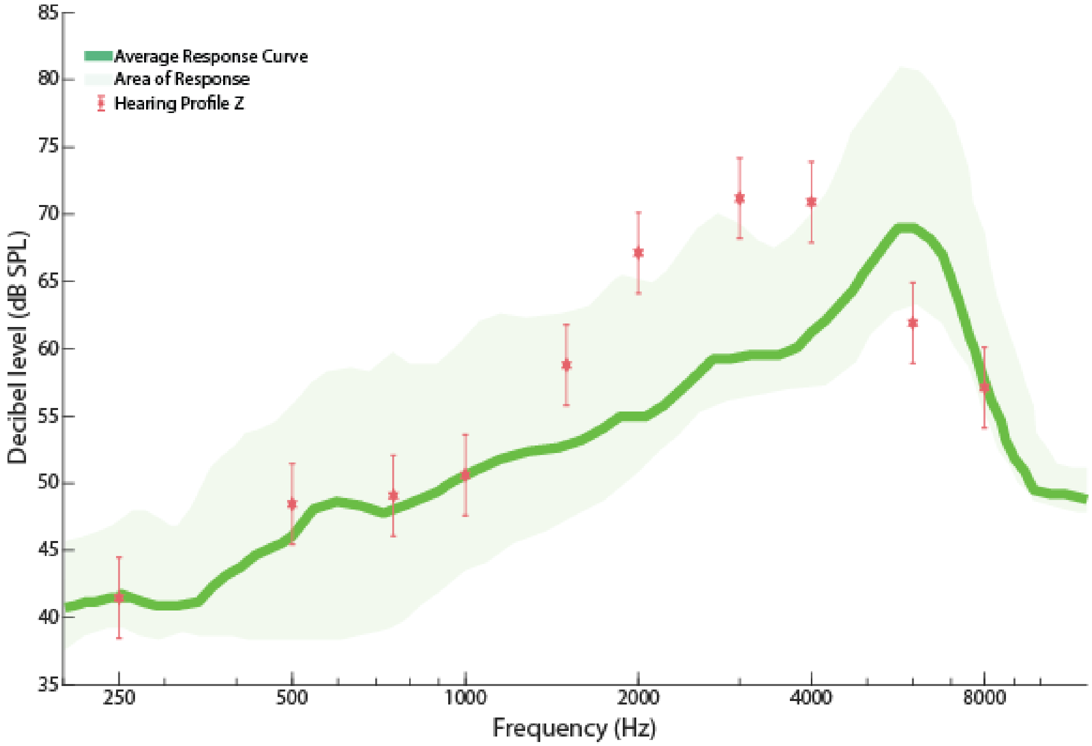


FIG. S73. Shows the ISTS 55 dB SPL Response (dB SPL) Curve with Hearing Profile Z Targets (dB SPL).

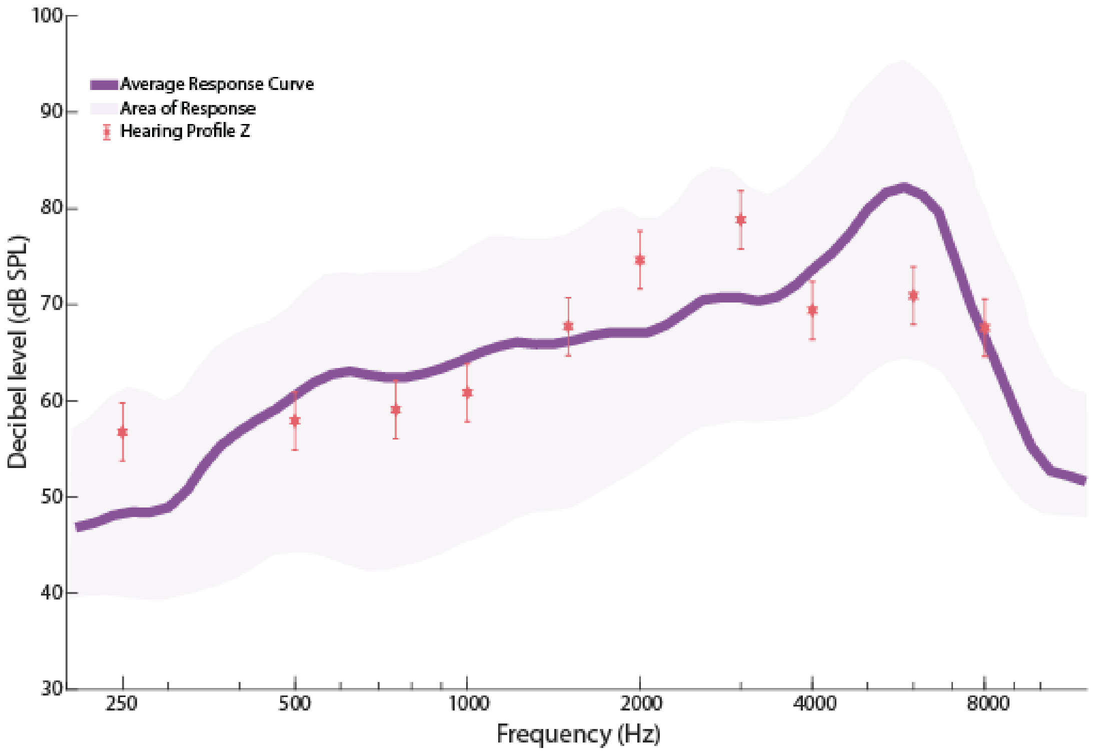


FIG. S74. Shows the ISTS 65 dB SPL Response (dB SPL) Curve with Hearing Profile Z Targets (dB SPL).

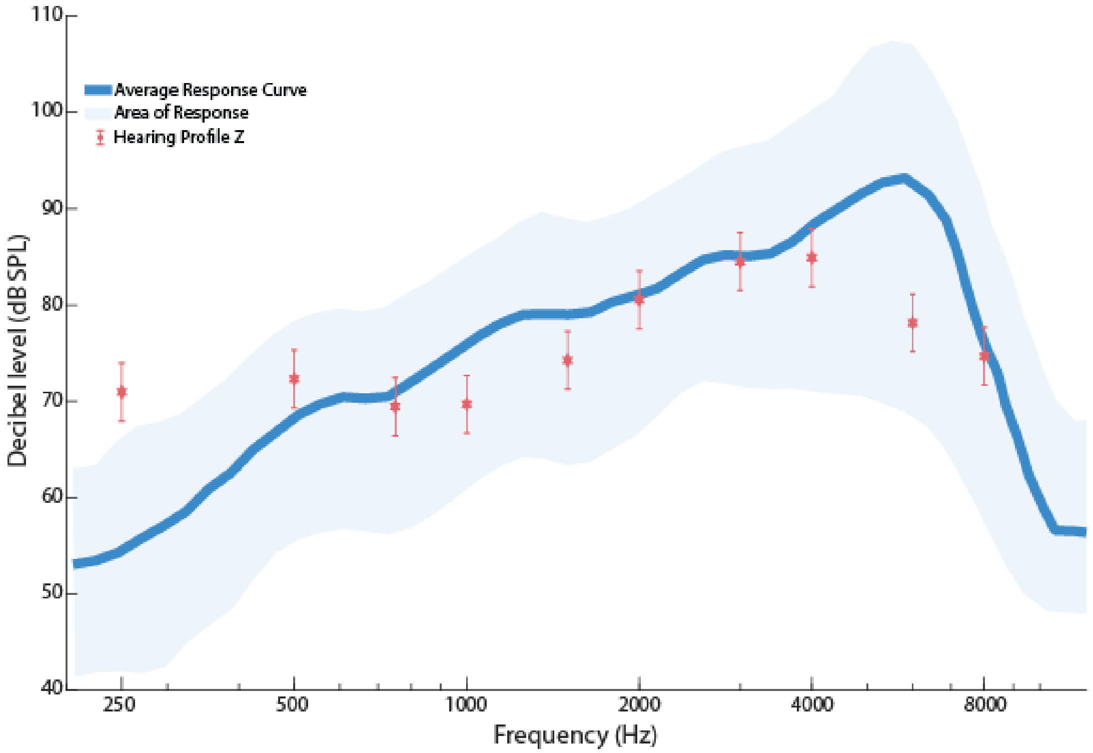


FIG. S75. Shows the ISTS 80 dB SPL Response (dB SPL) Curve with Hearing Profile Z Targets (dB SPL).

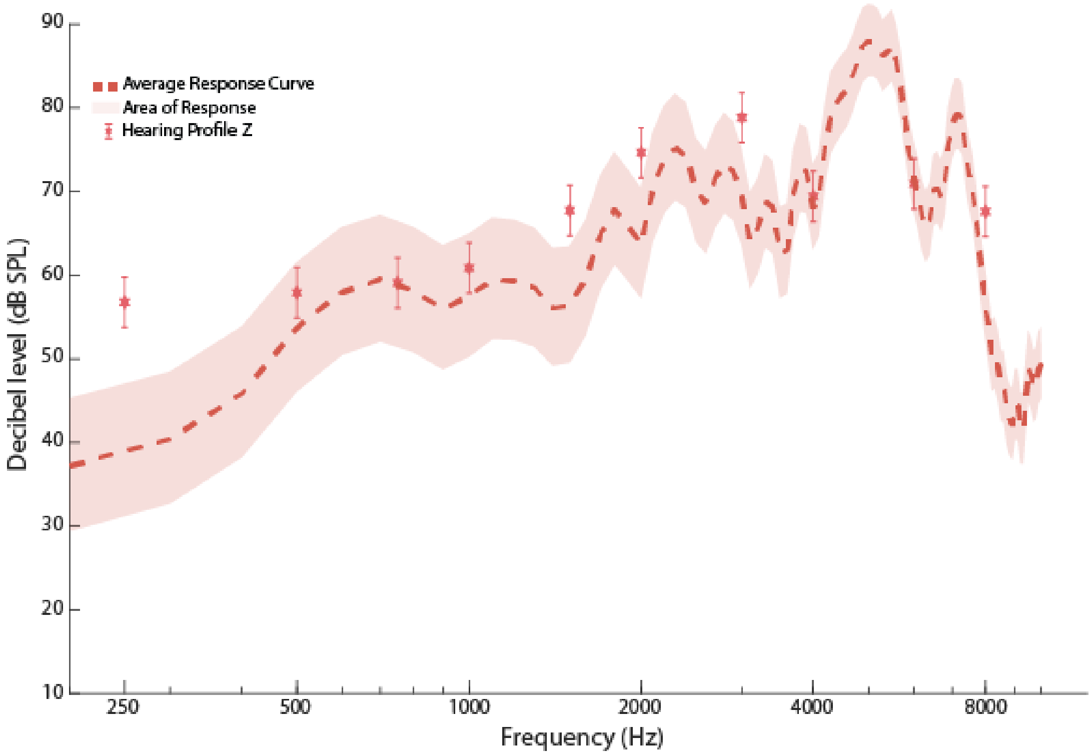


FIG. S76. Shows the G.R.A.S KEMAR 65 db SPL ISTS input Response (dB SPL) Curve (Right Ear) with Hearing Profile Z Targets (dB SPL).

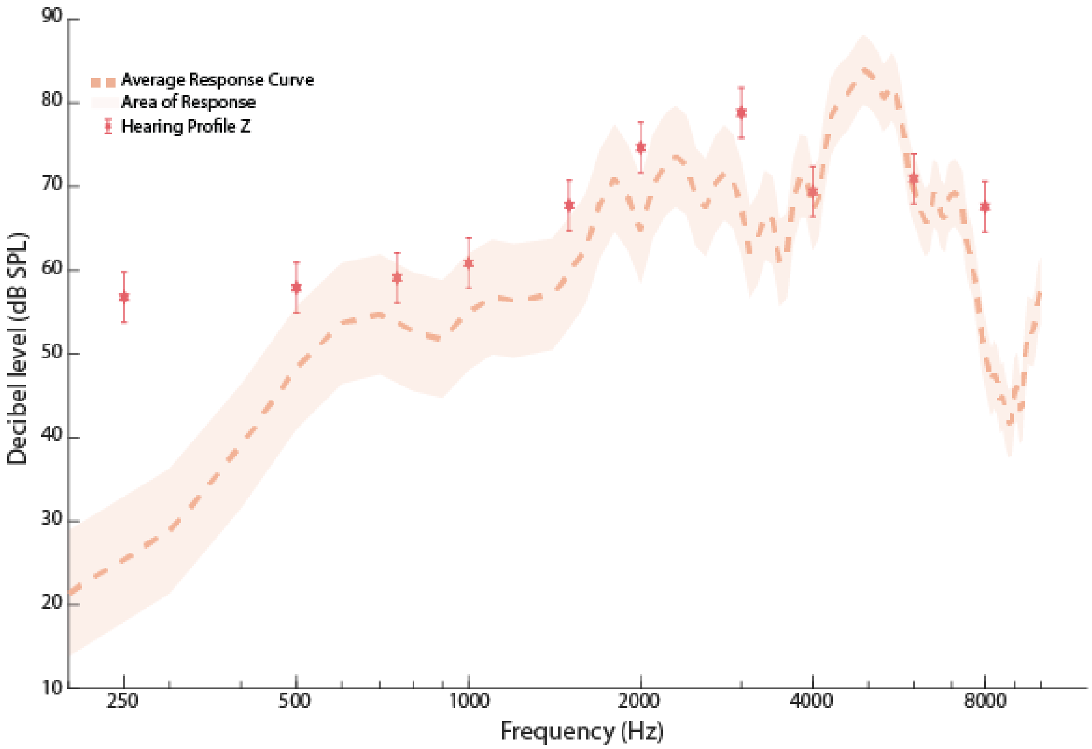


FIG. S77. Shows the G.R.A.S KEMAR 65 db SPL ISTS input Response (dB SPL) Curve (Left Ear) with Hearing Profile Z Targets (dB SPL).

| Frequencies (Hz) | Response (dB SPL) | Targets (dB SPL) | Meets Strict Criteria | Meets Loose Criteria |
|---|---|---|---|---|
| 250 | 41 | 40 | Yes | Yes |
| 500 | 46 | 43.56 | Yes | Yes |
| 750 | 48 | 43.52 | Yes | Yes |
| 1000 | 51 | 44.67 | No | Yes |
| 1500 | 53 | 51.3 | Yes | Yes |
| 2000 | 55 | 58.5 | Yes | Yes |
| 3000 | 59 | 64.3 | No | Yes |
| 4000 | 61 | 63.7 | Yes | Yes |
| 6000 | 69 | 54.4 | No | No |
| 8000 | 57 | 50.29 | No | Yes |

TABLE I. Shows the individual Targets (dB SPL) for Male Left Ear 60-69 ISTS 55 dB SPL for each Frequencies (Hz), and whether the device meets them under two criteria

.

| Frequencies (Hz) | Response (dB SPL) | Targets (dB SPL) | Meets Strict Criteria | Meets Loose Criteria |
|---|---|---|---|---|
| 250 | 48 | 55 | No | Yes |
| 500 | 61 | 57 | Yes | Yes |
| 750 | 62 | 56 | No | Yes |
| 1000 | 65 | 57 | No | Yes |
| 1500 | 66 | 61.6 | Yes | Yes |
| 2000 | 67 | 67.4 | Yes | Yes |
| 3000 | 71 | 71.86 | Yes | Yes |
| 4000 | 74 | 72 | Yes | Yes |
| 6000 | 81 | 64 | No | No |
| 8000 | 67 | 60.2 | No | Yes |

TABLE II. Shows the individual Targets (dB SPL) for Male Left Ear 60-69 ISTS 65 dB SPL for each Frequencies (Hz), and whether the device meets them under two criteria

.

| Frequencies (Hz) | Response (dB SPL) | Targets (dB SPL) | Meets Strict Criteria | Meets Loose Criteria |
|---|---|---|---|---|
| 250 | 54 | 71 | No | No |
| 500 | 69 | 72 | Yes | Yes |
| 750 | 71 | 69 | Yes | Yes |
| 1000 | 76 | 66 | No | No |
| 1500 | 79 | 70 | No | Yes |
| 2000 | 81 | 74.5 | No | Yes |
| 3000 | 85 | 77.76 | No | Yes |
| 4000 | 89 | 77.76 | No | No |
| 6000 | 92 | 70.5 | No | No |
| 8000 | 76 | 67.6 | No | Yes |

TABLE III. Shows the individual Targets (dB SPL) for Male Left Ear 60-69 ISTS 80 dB SPL for each Frequencies (Hz), and whether the device meets them under two criteria

.

| Frequencies (Hz) | Response (dB SPL) | Targets (dB SPL) | Meets Strict Criteria | Meets Loose Criteria |
|---|---|---|---|---|
| 250 | 39 | 55 | No | No |
| 500 | 53 | 57 | Yes | Yes |
| 750 | 58 | 56 | Yes | Yes |
| 1000 | 56 | 57 | Yes | Yes |
| 1500 | 55 | 61.6 | No | Yes |
| 2000 | 62 | 67 | Yes | Yes |
| 3000 | 67 | 71.86 | Yes | Yes |
| 4000 | 66 | 71 | Yes | Yes |
| 6000 | 69 | 64 | Yes | Yes |
| 8000 | 55 | 60 | Yes | Yes |

TABLE IV. Shows the individual Targets (dB SPL) for Male Left Ear 60-69 G.R.A.S KEMAR Left Ear ISTS 65 dB SPL for each Frequencies (Hz), and whether the device meets them under two criteria

| Frequencies (Hz) | Response (dB SPL) | Targets (dB SPL) | Meets Strict Criteria | Meets Loose Criteria |
|---|---|---|---|---|
| 250 | 25 | 55 | No | No |
| 500 | 48 | 57 | No | Yes |
| 750 | 54 | 56 | Yes | Yes |
| 1000 | 55 | 57 | Yes | Yes |
| 1500 | 60 | 61.6 | Yes | Yes |
| 2000 | 65 | 67.4 | Yes | Yes |
| 3000 | 67 | 71.86 | Yes | Yes |
| 4000 | 67 | 72 | No | Yes |
| 6000 | 70 | 64 | No | Yes |
| 8000 | 50 | 60.2 | No | No |

TABLE V. Shows the individual Targets (dB SPL) for Male Left Ear 60-69 G.R.A.S KEMAR Right Ear ISTS 65 dB SPL for each Frequencies (Hz), and whether the device meets them under two criteria

| Frequencies (Hz) | Response (dB SPL) | Targets (dB SPL) | Meets Strict Criteria | Meets Loose Criteria |
|---|---|---|---|---|
| 250 | 41 | 41.36 | Yes | Yes |
| 500 | 46 | 43.3 | Yes | Yes |
| 750 | 48 | 42.7 | No | Yes |
| 1000 | 51 | 42.9 | No | Yes |
| 1500 | 53 | 48.9 | Yes | Yes |
| 2000 | 55 | 56 | Yes | Yes |
| 3000 | 59 | 61.95 | Yes | Yes |
| 4000 | 61 | 64 | Yes | Yes |
| 6000 | 69 | 53.6 | No | No |
| 8000 | 57 | 48.77 | No | Yes |

TABLE VI. Shows the individual Targets (dB SPL) for Male Right Ear 60-69 ISTS 55 dB SPL for each Frequencies (Hz), and whether the device meets them under two criteria

.

| Frequencies (Hz) | Response (dB SPL) | Targets (dB SPL) | Meets Strict Criteria | Meets Loose Criteria |
|---|---|---|---|---|
| 250 | 48 | 55.67 | No | Yes |
| 500 | 61 | 57 | Yes | Yes |
| 750 | 62 | 56 | No | Yes |
| 1000 | 65 | 56 | No | Yes |
| 1500 | 66 | 60.4 | No | Yes |
| 2000 | 67 | 65 | Yes | Yes |
| 3000 | 71 | 70 | Yes | Yes |
| 4000 | 74 | 71.19 | Yes | Yes |
| 6000 | 81 | 62.6 | No | No |
| 8000 | 67 | 59.05 | No | Yes |

TABLE VII. Shows the individual Targets (dB SPL) for Male Right Ear 60-69 ISTS 65 dB SPL for each Frequencies (Hz), and whether the device meets them under two criteria

.

| Frequencies (Hz) | Response (dB SPL) | Targets (dB SPL) | Meets Strict Criteria | Meets Loose Criteria |
|---|---|---|---|---|
| 250 | 54 | 71 | No | No |
| 500 | 69 | 72 | Yes | Yes |
| 750 | 71 | 68.88 | Yes | Yes |
| 1000 | 76 | 66.89 | No | Yes |
| 1500 | 79 | 69.9 | No | Yes |
| 2000 | 81 | 73.55 | No | Yes |
| 3000 | 85 | 76.03 | No | Yes |
| 4000 | 89 | 76.8 | No | No |
| 6000 | 92 | 69.4 | No | No |
| 8000 | 76 | 66.51 | No | Yes |

TABLE VIII. Shows the individual Targets (dB SPL) for Male Right Ear 60-69 ISTS 80 dB SPL for each Frequencies (Hz), and whether the device meets them under two criteria

.

| Frequencies (Hz) | Response (dB SPL) | Targets (dB SPL) | Meets Strict Criteria | Meets Loose Criteria |
|---|---|---|---|---|
| 250 | 25 | 55.67 | No | No |
| 500 | 48 | 57 | No | Yes |
| 750 | 54 | 56 | Yes | Yes |
| 1000 | 55 | 56 | Yes | Yes |
| 1500 | 60 | 60.4 | Yes | Yes |
| 2000 | 65 | 65 | Yes | Yes |
| 3000 | 67 | 70 | Yes | Yes |
| 4000 | 67 | 71.19 | Yes | Yes |
| 6000 | 70 | 62.6 | No | Yes |
| 8000 | 50 | 59.05 | No | Yes |

TABLE IX. Shows the individual Targets (dB SPL) for Male Right Ear 60-69 G.R.A.S KEMAR Left Ear ISTS 65 dB SPL for each Frequencies (Hz), and whether the device meets them under two criteria

| Frequencies (Hz) | Response (dB SPL) | Targets (dB SPL) | Meets Strict Criteria | Meets Loose Criteria |
|---|---|---|---|---|
| 250 | 39 | 55.67 | No | No |
| 500 | 53 | 57 | Yes | Yes |
| 750 | 58 | 56 | Yes | Yes |
| 1000 | 56 | 56 | Yes | Yes |
| 1500 | 55 | 60 | Yes | Yes |
| 2000 | 62 | 65 | Yes | Yes |
| 3000 | 67 | 70 | Yes | Yes |
| 4000 | 66 | 71 | Yes | Yes |
| 6000 | 69 | 62.6 | No | Yes |
| 8000 | 55 | 59.05 | Yes | Yes |

TABLE X. Shows the individual Targets (dB SPL) for Male Right Ear 60-69 G.R.A.S KEMAR Right Ear ISTS 65 dB SPL for each Frequencies (Hz), and whether the device meets them under two criteria

| Frequencies (Hz) | Response (dB SPL) | Targets (dB SPL) | Meets Strict Criteria | Meets Loose Criteria |
|---|---|---|---|---|
| 250 | 41 | 41.35 | Yes | Yes |
| 500 | 46 | 47.72 | Yes | Yes |
| 750 | 48 | 48.55 | Yes | Yes |
| 1000 | 51 | 49.66 | Yes | Yes |
| 1500 | 53 | 56.58 | Yes | Yes |
| 2000 | 55 | 64.334 | No | Yes |
| 3000 | 59 | 67.94 | No | Yes |
| 4000 | 61 | 67.94 | No | Yes |
| 6000 | 69 | 59.07 | No | Yes |
| 8000 | 57 | 54.64 | Yes | Yes |

TABLE XI. Shows the individual Targets (dB SPL) for Male Left Ear 70-79 ISTS 55 dB SPL for each Frequencies (Hz), and whether the device meets them under two criteria

.

| Frequencies (Hz) | Response (dB SPL) | Targets (dB SPL) | Meets Strict Criteria | Meets Loose Criteria |
|---|---|---|---|---|
| 250 | 48 | 55.28 | No | Yes |
| 500 | 61 | 57.2 | Yes | Yes |
| 750 | 62 | 58.29 | Yes | Yes |
| 1000 | 65 | 59.68 | No | Yes |
| 1500 | 66 | 65.27 | Yes | Yes |
| 2000 | 67 | 71 | Yes | Yes |
| 3000 | 71 | 75.19 | Yes | Yes |
| 4000 | 74 | 75.54 | Yes | Yes |
| 6000 | 81 | 67.5 | No | No |
| 8000 | 67 | 63.72 | Yes | Yes |

TABLE XII. Shows the individual Targets (dB SPL) for Male Left Ear 70-79 ISTS 65 dB SPL for each Frequencies (Hz), and whether the device meets them under two criteria

.

| Frequencies (Hz) | Response (dB SPL) | Targets (dB SPL) | Meets Strict Criteria | Meets Loose Criteria |
|---|---|---|---|---|
| 250 | 54 | 71 | No | No |
| 500 | 69 | 72.7 | Yes | Yes |
| 750 | 71 | 69.78 | Yes | Yes |
| 1000 | 76 | 69.17 | No | Yes |
| 1500 | 79 | 73.2 | No | Yes |
| 2000 | 81 | 77.85 | Yes | Yes |
| 3000 | 85 | 81.02 | Yes | Yes |
| 4000 | 89 | 81.58 | No | Yes |
| 6000 | 92 | 74.56 | No | No |
| 8000 | 76 | 71.62 | Yes | Yes |

TABLE XIII. Shows the individual Targets (dB SPL) for Male Left Ear 70-79 ISTS 80 dB SPL for each Frequencies (Hz), and whether the device meets them under two criteria

.

| Frequencies (Hz) | Response (dB SPL) | Targets (dB SPL) | Meets Strict Criteria | Meets Loose Criteria |
|---|---|---|---|---|
| 250 | 39 | 55.28 | No | No |
| 500 | 53 | 57.2 | Yes | Yes |
| 750 | 58 | 58.29 | Yes | Yes |
| 1000 | 56 | 59.68 | Yes | Yes |
| 1500 | 55 | 65.27 | No | No |
| 2000 | 62 | 71 | No | Yes |
| 3000 | 67 | 75.19 | No | Yes |
| 4000 | 66 | 75.54 | No | Yes |
| 6000 | 69 | 67.5 | Yes | Yes |
| 8000 | 55 | 63.72 | No | Yes |

TABLE XIV. Shows the individual Targets (dB SPL) for Male Left Ear 70-79 G.R.A.S KEMAR Right Ear ISTS 65 dB SPL for each Frequencies (Hz), and whether the device meets them under two criteria

| Frequencies (Hz) | Response (dB SPL) | Targets (dB SPL) | Meets Strict Criteria | Meets Loose Criteria |
|---|---|---|---|---|
| 250 | 25 | 55.28 | No | No |
| 500 | 48 | 57.2 | No | Yes |
| 750 | 54 | 58.29 | Yes | Yes |
| 1000 | 55 | 59.68 | Yes | Yes |
| 1500 | 60 | 65.27 | No | Yes |
| 2000 | 65 | 71 | No | Yes |
| 3000 | 67 | 75.19 | No | Yes |
| 4000 | 67 | 75.54 | No | Yes |
| 6000 | 70 | 67.5 | Yes | Yes |
| 8000 | 50 | 63.72 | No | No |

TABLE XV. Shows the individual Targets (dB SPL) for Male Left Ear 70-79 G.R.A.S KEMAR Left Ear ISTS 65 dB SPL for each Frequencies (Hz), and whether the device meets them under two criteria

| Frequencies (Hz) | Response (dB SPL) | Targets (dB SPL) | Meets Strict Criteria | Meets Loose Criteria |
|---|---|---|---|---|
| 250 | 41 | 41.14 | Yes | Yes |
| 500 | 46 | 48.28 | Yes | Yes |
| 750 | 48 | 47.43 | Yes | Yes |
| 1000 | 51 | 48 | Yes | Yes |
| 1500 | 53 | 53.43 | Yes | Yes |
| 2000 | 55 | 60.28 | No | Yes |
| 3000 | 59 | 64.28 | No | Yes |
| 4000 | 61 | 64.28 | Yes | Yes |
| 6000 | 69 | 55.71 | No | No |
| 8000 | 57 | 52.28 | Yes | Yes |

TABLE XVI. Shows the individual Targets (dB SPL) for Male Right Ear 70-79 ISTS 55 dB SPL for each Frequencies (Hz), and whether the device meets them under two criteria

.

| Frequencies (Hz) | Response (dB SPL) | Targets (dB SPL) | Meets Strict Criteria | Meets Loose Criteria |
|---|---|---|---|---|
| 250 | 48 | 56 | No | Yes |
| 500 | 61 | 57.45 | Yes | Yes |
| 750 | 62 | 58.3 | Yes | Yes |
| 1000 | 65 | 58.86 | No | Yes |
| 1500 | 66 | 63.39 | Yes | Yes |
| 2000 | 67 | 68.207 | Yes | Yes |
| 3000 | 71 | 72.17 | Yes | Yes |
| 4000 | 74 | 72.17 | Yes | Yes |
| 6000 | 81 | 64.52 | No | No |
| 8000 | 67 | 61.13 | No | Yes |

TABLE XVII. Shows the individual Targets (dB SPL) for Male Right Ear 70-79 ISTS 65 dB SPL for each Frequencies (Hz), and whether the device meets them under two criteria

.

| Frequencies (Hz) | Response (dB SPL) | Targets (dB SPL) | Meets Strict Criteria | Meets Loose Criteria |
|---|---|---|---|---|
| 250 | 54 | 70.8 | No | No |
| 500 | 69 | 72.3 | Yes | Yes |
| 750 | 71 | 69.19 | Yes | Yes |
| 1000 | 76 | 68.6 | No | Yes |
| 1500 | 79 | 71.8 | No | Yes |
| 2000 | 81 | 75.8 | No | Yes |
| 3000 | 85 | 78.44 | No | Yes |
| 4000 | 89 | 78.46 | No | No |
| 6000 | 92 | 71.6 | No | No |
| 8000 | 76 | 69.1 | No | Yes |

TABLE XVIII. Shows the individual Targets (dB SPL) for Male Right Ear 70-79 ISTS 80 dB SPL for each Frequencies (Hz), and whether the device meets them under two criteria

.

| Frequencies (Hz) | Response (dB SPL) | Targets (dB SPL) | Meets Strict Criteria | Meets Loose Criteria |
|---|---|---|---|---|
| 250 | 39 | 56 | No | No |
| 500 | 53 | 57.45 | Yes | Yes |
| 750 | 58 | 58.3 | Yes | Yes |
| 1000 | 56 | 58.86 | Yes | Yes |
| 1500 | 55 | 63.39 | No | No |
| 2000 | 62 | 68.207 | No | Yes |
| 3000 | 67 | 72.17 | No | Yes |
| 4000 | 66 | 72.17 | No | Yes |
| 6000 | 69 | 64.52 | Yes | Yes |
| 8000 | 55 | 61.13 | No | Yes |

TABLE XIX. Shows the individual Targets (dB SPL) for Male Right Ear 70-79 G.R.A.S KEMAR Right Ear ISTS 65 dB SPL for each Frequencies (Hz), and whether the device meets them under two criteria

| Frequencies (Hz) | Response (dB SPL) | Targets (dB SPL) | Meets Strict Criteria | Meets Loose Criteria |
|---|---|---|---|---|
| 250 | 25 | 56 | No | No |
| 500 | 48 | 57.45 | No | Yes |
| 750 | 54 | 58.3 | Yes | Yes |
| 1000 | 55 | 58.86 | Yes | Yes |
| 1500 | 60 | 63.39 | Yes | Yes |
| 2000 | 65 | 68.207 | Yes | Yes |
| 3000 | 67 | 72.17 | No | Yes |
| 4000 | 67 | 72.17 | No | Yes |
| 6000 | 70 | 64.52 | No | Yes |
| 8000 | 50 | 61.13 | No | No |

TABLE XX. Shows the individual Targets (dB SPL) for Male Right Ear 70-79 G.R.A.S KEMAR Left Ear ISTS 65 dB SPL for each Frequencies (Hz), and whether the device meets them under two criteria

| Frequencies (Hz) | Response (dB SPL) | Targets (dB SPL) | Meets Strict Criteria | Meets Loose Criteria |
|---|---|---|---|---|
| 250 | 41 | 41.35 | Yes | Yes |
| 500 | 46 | 47.72 | Yes | Yes |
| 750 | 48 | 48.55 | Yes | Yes |
| 1000 | 51 | 49.66 | Yes | Yes |
| 1500 | 53 | 56.58 | Yes | Yes |
| 2000 | 55 | 64.334 | No | Yes |
| 3000 | 59 | 67.94 | No | Yes |
| 4000 | 61 | 67.94 | No | Yes |
| 6000 | 69 | 59.07 | No | Yes |
| 8000 | 57 | 54.64 | Yes | Yes |

TABLE XXI. Shows the individual Targets (dB SPL) for Male Left Ear 70-79 ISTS 55 dB SPL for each Frequencies (Hz), and whether the device meets them under two criteria

.

| Frequencies (Hz) | Response (dB SPL) | Targets (dB SPL) | Meets Strict Criteria | Meets Loose Criteria |
|---|---|---|---|---|
| 250 | 48 | 55.28 | No | Yes |
| 500 | 61 | 57.2 | Yes | Yes |
| 750 | 62 | 58.29 | Yes | Yes |
| 1000 | 65 | 59.68 | No | Yes |
| 1500 | 66 | 65.27 | Yes | Yes |
| 2000 | 67 | 71 | Yes | Yes |
| 3000 | 71 | 75.19 | Yes | Yes |
| 4000 | 74 | 75.54 | Yes | Yes |
| 6000 | 81 | 67.5 | No | No |
| 8000 | 67 | 63.72 | Yes | Yes |

TABLE XXII. Shows the individual Targets (dB SPL) for Male Left Ear 70-79 ISTS 65 dB SPL for each Frequencies (Hz), and whether the device meets them under two criteria

.

| Frequencies (Hz) | Response (dB SPL) | Targets (dB SPL) | Meets Strict Criteria | Meets Loose Criteria |
|---|---|---|---|---|
| 250 | 54 | 71 | No | No |
| 500 | 69 | 72.7 | Yes | Yes |
| 750 | 71 | 69.78 | Yes | Yes |
| 1000 | 76 | 69.17 | No | Yes |
| 1500 | 79 | 73.2 | No | Yes |
| 2000 | 81 | 77.85 | Yes | Yes |
| 3000 | 85 | 81.02 | Yes | Yes |
| 4000 | 89 | 81.58 | No | Yes |
| 6000 | 92 | 74.56 | No | No |
| 8000 | 76 | 71.62 | Yes | Yes |

TABLE XXIII. Shows the individual Targets (dB SPL) for Male Left Ear 70-79 ISTS 80 dB SPL for each Frequencies (Hz), and whether the device meets them under two criteria

.

| Frequencies (Hz) | Response (dB SPL) | Targets (dB SPL) | Meets Strict Criteria | Meets Loose Criteria |
|---|---|---|---|---|
| 250 | 39 | 56 | No | No |
| 500 | 53 | 57.45 | Yes | Yes |
| 750 | 58 | 58.3 | Yes | Yes |
| 1000 | 56 | 58.86 | Yes | Yes |
| 1500 | 55 | 63.39 | No | No |
| 2000 | 62 | 68.207 | No | Yes |
| 3000 | 67 | 72.17 | No | Yes |
| 4000 | 66 | 72.17 | No | Yes |
| 6000 | 69 | 64.52 | Yes | Yes |
| 8000 | 55 | 61.13 | No | Yes |

TABLE XXIV. Shows the individual Targets (dB SPL) for Male Left Ear 70-79 G.R.A.S KEMAR Right Ear ISTS 65 dB SPL for each Frequencies (Hz), and whether the device meets them under two criteria

| Frequencies (Hz) | Response (dB SPL) | Targets (dB SPL) | Meets Strict Criteria | Meets Loose Criteria |
|---|---|---|---|---|
| 250 | 25 | 55.28 | No | No |
| 500 | 48 | 57.2 | No | Yes |
| 750 | 54 | 58.29 | Yes | Yes |
| 1000 | 55 | 59.68 | Yes | Yes |
| 1500 | 60 | 65.27 | No | Yes |
| 2000 | 65 | 71 | No | Yes |
| 3000 | 67 | 75.19 | No | Yes |
| 4000 | 67 | 75.54 | No | Yes |
| 6000 | 70 | 67.5 | Yes | Yes |
| 8000 | 50 | 63.72 | No | No |

TABLE XXV. Shows the individual Targets (dB SPL) for Male Left Ear 70-79 G.R.A.S KEMAR Left Ear ISTS 65 dB SPL for each Frequencies (Hz), and whether the device meets them under two criteria

| Frequencies (Hz) | Response (dB SPL) | Targets (dB SPL) | Meets Strict Criteria | Meets Loose Criteria |
|---|---|---|---|---|
| 250 | 41 | 41.08 | Yes | Yes |
| 500 | 46 | 44.44 | Yes | Yes |
| 750 | 48 | 41.56 | No | Yes |
| 1000 | 51 | 40.11 | No | No |
| 1500 | 53 | 45.49 | No | Yes |
| 2000 | 55 | 51.16 | Yes | Yes |
| 3000 | 59 | 54.83 | Yes | Yes |
| 4000 | 61 | 53.95 | No | Yes |
| 6000 | 69 | 45.09 | No | No |
| 8000 | 57 | 41.37 | No | No |

TABLE XXVI. Shows the individual Targets (dB SPL) for Female Left Ear 60-69 ISTS 55 dB SPL for each Frequencies (Hz), and whether the device meets them under two criteria

.

| Frequencies (Hz) | Response (dB SPL) | Targets (dB SPL) | Meets Strict Criteria | Meets Loose Criteria |
|---|---|---|---|---|
| 250 | 48 | 56.03 | No | Yes |
| 500 | 61 | 57.59 | Yes | Yes |
| 750 | 62 | 54.45 | No | Yes |
| 1000 | 65 | 53.8 | No | No |
| 1500 | 66 | 57.59 | No | Yes |
| 2000 | 67 | 61.98 | No | Yes |
| 3000 | 71 | 64.808 | No | Yes |
| 4000 | 74 | 64.5 | No | Yes |
| 6000 | 81 | 56.18 | No | No |
| 8000 | 67 | 52.42 | No | No |

TABLE XXVII. Shows the individual Targets (dB SPL) for Female Left Ear 60-69 ISTS 65 dB SPL for each Frequencies (Hz), and whether the device meets them under two criteria

.

| Frequencies (Hz) | Response (dB SPL) | Targets (dB SPL) | Meets Strict Criteria | Meets Loose Criteria |
|---|---|---|---|---|
| 250 | 54 | 70.89 | No | No |
| 500 | 69 | 72.77 | Yes | Yes |
| 750 | 71 | 68.91 | Yes | Yes |
| 1000 | 76 | 66.23 | No | Yes |
| 1500 | 79 | 68.61 | No | No |
| 2000 | 81 | 71.28 | No | Yes |
| 3000 | 85 | 73.36 | No | No |
| 4000 | 89 | 72.62 | No | No |
| 6000 | 92 | 65.05 | No | No |
| 8000 | 76 | 61.63 | No | No |

TABLE XXVIII. Shows the individual Targets (dB SPL) for Female Left Ear 60-69 ISTS 80 dB SPL for each Frequencies (Hz), and whether the device meets them under two criteria

.

| Frequencies (Hz) | Response (dB SPL) | Targets (dB SPL) | Meets Strict Criteria | Meets Loose Criteria |
|---|---|---|---|---|
| 250 | 39 | 56.03 | No | No |
| 500 | 53 | 57.59 | Yes | Yes |
| 750 | 58 | 54.45 | Yes | Yes |
| 1000 | 56 | 53.8 | Yes | Yes |
| 1500 | 55 | 57.59 | Yes | Yes |
| 2000 | 62 | 61.98 | Yes | Yes |
| 3000 | 67 | 64.808 | Yes | Yes |
| 4000 | 66 | 64.5 | Yes | Yes |
| 6000 | 69 | 56.18 | No | No |
| 8000 | 55 | 52.42 | Yes | Yes |

TABLE XXIX. Shows the individual Targets (dB SPL) for Female Left Ear 60-69 G.R.A.S KEMAR Right Ear ISTS 65 dB SPL for each Frequencies (Hz), and whether the device meets them under two criteria.

| Frequencies (Hz) | Response (dB SPL) | Targets (dB SPL) | Meets Strict Criteria | Meets Loose Criteria |
|---|---|---|---|---|
| 250 | 25 | 56 | No | No |
| 500 | 48 | 57.59 | No | Yes |
| 750 | 54 | 54.45 | Yes | Yes |
| 1000 | 55 | 53.8 | Yes | Yes |
| 1500 | 60 | 57.59 | Yes | Yes |
| 2000 | 65 | 61.98 | Yes | Yes |
| 3000 | 67 | 64.808 | Yes | Yes |
| 4000 | 67 | 64.5 | Yes | Yes |
| 6000 | 70 | 56.18 | No | No |
| 8000 | 50 | 52.42 | Yes | Yes |

TABLE XXX. Shows the individual Targets (dB SPL) for Female Left Ear 60-69 G.R.A.S KEMAR Left Ear ISTS 65 dB SPL for each Frequencies (Hz), and whether the device meets them under two criteria.

| Frequencies (Hz) | Response (dB SPL) | Targets (dB SPL) | Meets Strict Criteria | Meets Loose Criteria |
|---|---|---|---|---|
| 250 | 41 | 41.18 | Yes | Yes |
| 500 | 46 | 43.36 | Yes | Yes |
| 750 | 48 | 41.13 | No | Yes |
| 1000 | 51 | 39.89 | No | No |
| 1500 | 53 | 45.71 | No | Yes |
| 2000 | 55 | 51.8 | Yes | Yes |
| 3000 | 59 | 54.8 | Yes | Yes |
| 4000 | 61 | 53.7 | No | Yes |
| 6000 | 69 | 44.55 | No | No |
| 8000 | 57 | 40.8 | No | No |

TABLE XXXI. Shows the individual Targets (dB SPL) for Female Right Ear 60-69 ISTS 55 dB SPL for each Frequencies (Hz), and whether the device meets them under two criteria

.

| Frequencies (Hz) | Response (dB SPL) | Targets (dB SPL) | Meets Strict Criteria | Meets Loose Criteria |
|---|---|---|---|---|
| 250 | 48 | 55.6 | No | Yes |
| 500 | 61 | 57.54 | Yes | Yes |
| 750 | 62 | 54.66 | No | Yes |
| 1000 | 65 | 54.07 | No | No |
| 1500 | 66 | 57.45 | No | Yes |
| 2000 | 67 | 61.41 | No | Yes |
| 3000 | 71 | 64.95 | No | Yes |
| 4000 | 74 | 63.06 | No | No |
| 6000 | 81 | 55.06 | No | No |
| 8000 | 67 | 50.81 | No | No |

TABLE XXXII. Shows the individual Targets (dB SPL) for Female Right Ear 60-69 ISTS 65 dB SPL for each Frequencies (Hz), and whether the device meets them under two criteria

.

| Frequencies (Hz) | Response (dB SPL) | Targets (dB SPL) | Meets Strict Criteria | Meets Loose Criteria |
|---|---|---|---|---|
| 250 | 54 | 70.89 | No | No |
| 500 | 69 | 72.77 | Yes | Yes |
| 750 | 71 | 68.91 | Yes | Yes |
| 1000 | 76 | 66.23 | No | Yes |
| 1500 | 79 | 68.61 | No | No |
| 2000 | 81 | 71.28 | No | Yes |
| 3000 | 85 | 73.36 | No | No |
| 4000 | 89 | 72.62 | No | No |
| 6000 | 92 | 65.05 | No | No |
| 8000 | 76 | 61.63 | No | No |

TABLE XXXIII. Shows the individual Targets (dB SPL) for Female Right Ear 60-69 ISTS 80 dB SPL for each Frequencies (Hz), and whether the device meets them under two criteria

.

| Frequencies (Hz) | Response (dB SPL) | Targets (dB SPL) | Meets Strict Criteria | Meets Loose Criteria |
|---|---|---|---|---|
| 250 | 39 | 55.6 | No | No |
| 500 | 53 | 57.54 | Yes | Yes |
| 750 | 58 | 54.66 | Yes | Yes |
| 1000 | 56 | 54.07 | Yes | Yes |
| 1500 | 55 | 57.45 | Yes | Yes |
| 2000 | 62 | 61.41 | Yes | Yes |
| 3000 | 67 | 64.95 | Yes | Yes |
| 4000 | 66 | 63.06 | Yes | Yes |
| 6000 | 69 | 55.06 | No | No |
| 8000 | 55 | 50.81 | Yes | Yes |

TABLE XXXIV. Shows the individual Targets (dB SPL) for Female Right Ear 60-69 G.R.A.S KEMAR Right Ear ISTS 65 dB SPL for each Frequencies (Hz), and whether the device meets them under two criteria.

| Frequencies (Hz) | Response (dB SPL) | Targets (dB SPL) | Meets Strict Criteria | Meets Loose Criteria |
|---|---|---|---|---|
| 250 | 25 | 55.6 | No | No |
| 500 | 48 | 57.54 | No | Yes |
| 750 | 54 | 54.66 | Yes | Yes |
| 1000 | 55 | 54.07 | Yes | Yes |
| 1500 | 60 | 57.45 | Yes | Yes |
| 2000 | 65 | 61.41 | Yes | Yes |
| 3000 | 67 | 64.95 | Yes | Yes |
| 4000 | 67 | 63.06 | Yes | Yes |
| 6000 | 70 | 55.06 | No | No |
| 8000 | 50 | 50.81 | Yes | Yes |

TABLE XXXV. Shows the individual Targets (dB SPL) for Female Right Ear 60-69 G.R.A.S KEMAR Left Ear ISTS 65 dB SPL for each Frequencies (Hz), and whether the device meets them under two criteria.

| Frequencies (Hz) | Response (dB SPL) | Targets (dB SPL) | Meets Strict Criteria | Meets Loose Criteria |
|---|---|---|---|---|
| 250 | 41 | 41.09 | Yes | Yes |
| 500 | 46 | 46.34 | Yes | Yes |
| 750 | 48 | 44.83 | Yes | Yes |
| 1000 | 51 | 44.45 | No | Yes |
| 1500 | 53 | 50.37 | Yes | Yes |
| 2000 | 55 | 56.83 | Yes | Yes |
| 3000 | 59 | 59.94 | Yes | Yes |
| 4000 | 61 | 58.98 | Yes | Yes |
| 6000 | 69 | 50.84 | No | No |
| 8000 | 57 | 47.41 | No | Yes |

TABLE XXXVI. Shows the individual Targets (dB SPL) for Female Left Ear 70-79 ISTS 55 dB SPL for each Frequencies (Hz), and whether the device meets them under two criteria

.

| Frequencies (Hz) | Response (dB SPL) | Targets (dB SPL) | Meets Strict Criteria | Meets Loose Criteria |
|---|---|---|---|---|
| 250 | 48 | 55.79 | No | Yes |
| 500 | 61 | 57.75 | Yes | Yes |
| 750 | 62 | 57.47 | Yes | Yes |
| 1000 | 65 | 57.75 | No | Yes |
| 1500 | 66 | 61.68 | Yes | Yes |
| 2000 | 67 | 66.44 | Yes | Yes |
| 3000 | 71 | 68.41 | Yes | Yes |
| 4000 | 74 | 68.41 | No | Yes |
| 6000 | 81 | 60.28 | No | No |
| 8000 | 67 | 57.19 | No | Yes |

TABLE XXXVII. Shows the individual Targets (dB SPL) for Female Left Ear 70-79 ISTS 65 dB SPL for each Frequencies (Hz), and whether the device meets them under two criteria

.

| Frequencies (Hz) | Response (dB SPL) | Targets (dB SPL) | Meets Strict Criteria | Meets Loose Criteria |
|---|---|---|---|---|
| 250 | 54 | 71.21 | No | No |
| 500 | 69 | 72.64 | Yes | Yes |
| 750 | 71 | 69.29 | Yes | Yes |
| 1000 | 76 | 68.17 | No | Yes |
| 1500 | 79 | 70.44 | No | Yes |
| 2000 | 81 | 74.11 | No | Yes |
| 3000 | 85 | 75.53 | No | Yes |
| 4000 | 89 | 75.26 | No | No |
| 6000 | 92 | 67.4 | No | No |
| 8000 | 76 | 64.88 | No | No |

TABLE XXXVIII. Shows the individual Targets (dB SPL) for Female Left Ear 70-79 ISTS 80 dB SPL for each Frequencies (Hz), and whether the device meets them under two criteria

.

| Frequencies (Hz) | Response (dB SPL) | Targets (dB SPL) | Meets Strict Criteria | Meets Loose Criteria |
|---|---|---|---|---|
| 250 | 39 | 55.95 | No | No |
| 500 | 53 | 57.47 | Yes | Yes |
| 750 | 58 | 56.17 | Yes | Yes |
| 1000 | 56 | 56.78 | Yes | Yes |
| 1500 | 55 | 60.23 | No | Yes |
| 2000 | 62 | 65.91 | Yes | Yes |
| 3000 | 67 | 67.77 | Yes | Yes |
| 4000 | 66 | 66.8 | Yes | Yes |
| 6000 | 69 | 59.47 | No | Yes |
| 8000 | 55 | 55.96 | Yes | Yes |

TABLE XXXIX. Shows the individual Targets (dB SPL) for Female Left Ear 70-79 G.R.A.S KEMAR Right Ear ISTS 65 dB SPL for each Frequencies (Hz), and whether the device meets them under two criteria.

| Frequencies (Hz) | Response (dB SPL) | Targets (dB SPL) | Meets Strict Criteria | Meets Loose Criteria |
|---|---|---|---|---|
| 250 | 25 | 55.95 | No | No |
| 500 | 48 | 57.47 | No | Yes |
| 750 | 54 | 56.17 | Yes | Yes |
| 1000 | 55 | 56.78 | Yes | Yes |
| 1500 | 60 | 60.23 | Yes | Yes |
| 2000 | 65 | 65.91 | Yes | Yes |
| 3000 | 67 | 67.77 | Yes | Yes |
| 4000 | 67 | 66.8 | Yes | Yes |
| 6000 | 70 | 59.47 | No | No |
| 8000 | 50 | 55.96 | No | Yes |

TABLE XL. Shows the individual Targets (dB SPL) for Female Left Ear 70-79 G.R.A.S KEMAR Left Ear ISTS 65 dB SPL for each Frequencies (Hz), and whether the device meets them under two criteria.

| Frequencies (Hz) | Response (dB SPL) | Targets (dB SPL) | Meets Strict Criteria | Meets Loose Criteria |
|---|---|---|---|---|
| 250 | 41 | 40.87 | Yes | Yes |
| 500 | 46 | 47.69 | Yes | Yes |
| 750 | 48 | 46.25 | Yes | Yes |
| 1000 | 51 | 46.22 | Yes | Yes |
| 1500 | 53 | 51.53 | Yes | Yes |
| 2000 | 55 | 57.17 | Yes | Yes |
| 3000 | 59 | 60.24 | Yes | Yes |
| 4000 | 61 | 59.93 | Yes | Yes |
| 6000 | 69 | 51.01 | No | No |
| 8000 | 57 | 47.467 | No | Yes |

TABLE XLI. Shows the individual Targets (dB SPL) for Female Right Ear 70-79 ISTS 55 dB SPL for each Frequencies (Hz), and whether the device meets them under two criteria

.

| Frequencies (Hz) | Response (dB SPL) | Targets (dB SPL) | Meets Strict Criteria | Meets Loose Criteria |
|---|---|---|---|---|
| 250 | 48 | 55.79 | No | Yes |
| 500 | 61 | 57.75 | Yes | Yes |
| 750 | 62 | 57.47 | Yes | Yes |
| 1000 | 65 | 57.75 | No | Yes |
| 1500 | 66 | 61.68 | Yes | Yes |
| 2000 | 67 | 66.44 | Yes | Yes |
| 3000 | 71 | 68.41 | Yes | Yes |
| 4000 | 74 | 68.41 | No | Yes |
| 6000 | 81 | 60.28 | No | No |
| 8000 | 67 | 57.19 | No | Yes |

TABLE XLII. Shows the individual Targets (dB SPL) for Female Right Ear 70-79 ISTS 65 dB SPL for each Frequencies (Hz), and whether the device meets them under two criteria.

| Frequencies (Hz) | Response (dB SPL) | Targets (dB SPL) | Meets Strict Criteria | Meets Loose Criteria |
|---|---|---|---|---|
| 250 | 54 | 71.03 | No | No |
| 500 | 69 | 72.17 | Yes | Yes |
| 750 | 71 | 69.34 | Yes | Yes |
| 1000 | 76 | 67.64 | No | Yes |
| 1500 | 79 | 70.47 | No | Yes |
| 2000 | 81 | 74.43 | No | Yes |
| 3000 | 85 | 75.56 | No | Yes |
| 4000 | 89 | 75.28 | No | No |
| 6000 | 92 | 67.92 | No | No |
| 8000 | 76 | 65.09 | No | No |

TABLE XLIII. Shows the individual Targets (dB SPL) for Female Right Ear 70-79 ISTS 80 dB SPL for each Frequencies (Hz), and whether the device meets them under two criteria.

| Frequencies (Hz) | Response (dB SPL) | Targets (dB SPL) | Meets Strict Criteria | Meets Loose Criteria |
|---|---|---|---|---|
| 250 | 39 | 55.79 | No | No |
| 500 | 53 | 57.75 | Yes | Yes |
| 750 | 58 | 57.47 | Yes | Yes |
| 1000 | 56 | 57.75 | Yes | Yes |
| 1500 | 55 | 61.68 | No | Yes |
| 2000 | 62 | 66.44 | Yes | Yes |
| 3000 | 67 | 68.41 | Yes | Yes |
| 4000 | 66 | 68.41 | Yes | Yes |
| 6000 | 69 | 60.28 | No | Yes |
| 8000 | 55 | 57.19 | Yes | Yes |

TABLE XLIV. Shows the individual Targets (dB SPL) for Female Right Ear 70-79 G.R.A.S KEMAR Right Ear ISTS 65 dB SPL for each Frequencies (Hz), and whether the device meets them under two criteria.

| Frequencies (Hz) | Response (dB SPL) | Targets (dB SPL) | Meets Strict Criteria | Meets Loose Criteria |
|---|---|---|---|---|
| 250 | 25 | 55.79 | No | No |
| 500 | 48 | 57.75 | No | Yes |
| 750 | 54 | 57.47 | Yes | Yes |
| 1000 | 55 | 57.75 | Yes | Yes |
| 1500 | 60 | 61.68 | Yes | Yes |
| 2000 | 65 | 66.44 | Yes | Yes |
| 3000 | 67 | 68.41 | Yes | Yes |
| 4000 | 67 | 68.41 | Yes | Yes |
| 6000 | 70 | 60.28 | No | Yes |
| 8000 | 50 | 57.19 | No | Yes |

TABLE XLV. Shows the individual Targets (dB SPL) for Female Right Ear 70-79 G.R.A.S KEMAR Left Ear ISTS 65 dB SPL for each Frequencies (Hz), and whether the device meets them under two criteria.

| Frequencies (Hz) | Response (dB SPL) | Targets (dB SPL) | Meets Strict Criteria | Meets Loose Criteria |
|---|---|---|---|---|
| 250 | 41 | 40.89 | Yes | Yes |
| 500 | 46 | 45.4 | Yes | Yes |
| 750 | 48 | 41.81 | No | Yes |
| 1000 | 51 | 40.31 | No | No |
| 1500 | 53 | 45.21 | No | Yes |
| 2000 | 55 | 51.23 | Yes | Yes |
| 3000 | 59 | 53.91 | No | Yes |
| 4000 | 61 | 52.54 | No | Yes |
| 6000 | 69 | 44.2 | No | No |
| 8000 | 57 | 40 | No | No |

TABLE XLVI. Shows the individual Targets (dB SPL) for Hearing Profile X ISTS 55 dB SPL for each Frequencies (Hz), and whether the device meets them under two criteria.

| Frequencies (Hz) | Response (dB SPL) | Targets (dB SPL) | Meets Strict Criteria | Meets Loose Criteria |
|---|---|---|---|---|
| 250 | 48 | 56 | No | Yes |
| 500 | 61 | 57.7 | Yes | Yes |
| 750 | 62 | 54.77 | No | Yes |
| 1000 | 65 | 54.35 | No | No |
| 1500 | 66 | 57.85 | No | Yes |
| 2000 | 67 | 61.77 | No | Yes |
| 3000 | 71 | 63.31 | No | Yes |
| 4000 | 74 | 62.47 | No | No |
| 6000 | 81 | 54.63 | No | No |
| 8000 | 67 | 50.43 | No | No |

TABLE XLVII. Shows the individual Targets (dB SPL) for Hearing Profile X ISTS 65 dB SPL for each Frequencies (Hz), and whether the device meets them under two criteria.

| Frequencies (Hz) | Response (dB SPL) | Targets (dB SPL) | Meets Strict Criteria | Meets Loose Criteria |
|---|---|---|---|---|
| 250 | 54 | 70.82 | No | No |
| 500 | 69 | 72.65 | Yes | Yes |
| 750 | 71 | 69.46 | Yes | Yes |
| 1000 | 76 | 67.64 | No | Yes |
| 1500 | 79 | 68.7 | No | No |
| 2000 | 81 | 71.59 | No | Yes |
| 3000 | 85 | 72.65 | No | No |
| 4000 | 89 | 71.12 | No | No |
| 6000 | 92 | 63.23 | No | No |
| 8000 | 76 | 59.59 | No | No |

TABLE XLVIII. Shows the individual Targets (dB SPL) for Hearing Profile X ISTS 80 dB SPL for each Frequencies (Hz), and whether the device meets them under two criteria.

| Frequencies (Hz) | Response (dB SPL) | Targets (dB SPL) | Meets Strict Criteria | Meets Loose Criteria |
|---|---|---|---|---|
| 250 | 39 | 56 | No | No |
| 500 | 53 | 57.7 | No | Yes |
| 750 | 58 | 54.77 | Yes | Yes |
| 1000 | 56 | 54.35 | Yes | Yes |
| 1500 | 55 | 57.85 | No | Yes |
| 2000 | 62 | 61.77 | Yes | Yes |
| 3000 | 67 | 63.31 | Yes | Yes |
| 4000 | 66 | 62.47 | Yes | Yes |
| 6000 | 69 | 54.63 | No | Yes |
| 8000 | 55 | 50.43 | Yes | Yes |

TABLE XLIX. Shows the individual Targets (dB SPL) for Hearing Profile X G.R.A.S KEMAR Right Ear ISTS 65 dB SPL for each Frequencies (Hz), and whether the device meets them under two criteria.

| Frequencies (Hz) | Response (dB SPL) | Targets (dB SPL) | Meets Strict Criteria | Meets Loose Criteria |
|---|---|---|---|---|
| 250 | 25 | 56 | No | No |
| 500 | 48 | 57.7 | No | Yes |
| 750 | 54 | 54.77 | Yes | Yes |
| 1000 | 55 | 54.35 | Yes | Yes |
| 1500 | 60 | 57.85 | Yes | Yes |
| 2000 | 65 | 61.77 | Yes | Yes |
| 3000 | 67 | 63.31 | Yes | Yes |
| 4000 | 67 | 62.47 | Yes | Yes |
| 6000 | 70 | 54.63 | No | No |
| 8000 | 50 | 50.43 | Yes | Yes |

TABLE L. Shows the individual Targets (dB SPL) for Hearing Profile X G.R.A.S KEMAR Left Ear ISTS 65 dB SPL for each Frequencies (Hz), and whether the device meets them under two criteria.

| Frequencies (Hz) | Response (dB SPL) | Targets (dB SPL) | Meets Strict Criteria | Meets Loose Criteria |
|---|---|---|---|---|
| 250 | 41 | 41.04 | Yes | Yes |
| 500 | 46 | 49.51 | Yes | Yes |
| 750 | 48 | 48.15 | Yes | Yes |
| 1000 | 51 | 47.85 | Yes | Yes |
| 1500 | 53 | 52.39 | Yes | Yes |
| 2000 | 55 | 58.79 | Yes | Yes |
| 3000 | 59 | 61.82 | Yes | Yes |
| 4000 | 61 | 61.61 | Yes | Yes |
| 6000 | 69 | 53.75 | No | No |
| 8000 | 57 | 50.11 | No | Yes |

TABLE LI. Shows the individual Targets (dB SPL) for Hearing Profile Y ISTS 55 dB SPL for each Frequencies (Hz), and whether the device meets them under two criteria.

| Frequencies (Hz) | Response (dB SPL) | Targets (dB SPL) | Meets Strict Criteria | Meets Loose Criteria |
|---|---|---|---|---|
| 250 | 48 | 56.14 | No | Yes |
| 500 | 61 | 59.14 | Yes | Yes |
| 750 | 62 | 58.84 | Yes | Yes |
| 1000 | 65 | 59.14 | No | No |
| 1500 | 66 | 62.75 | Yes | Yes |
| 2000 | 67 | 66.96 | Yes | Yes |
| 3000 | 71 | 69.96 | Yes | Yes |
| 4000 | 74 | 70.11 | Yes | Yes |
| 6000 | 81 | 62.75 | No | No |
| 8000 | 67 | 58.99 | No | Yes |

TABLE LII. Shows the individual Targets (dB SPL) for Hearing Profile Y ISTS 65 dB SPL for each Frequencies (Hz), and whether the device meets them under two criteria.

| Frequencies (Hz) | Response (dB SPL) | Targets (dB SPL) | Meets Strict Criteria | Meets Loose Criteria |
|---|---|---|---|---|
| 250 | 54 | 71.14 | No | No |
| 500 | 69 | 72.57 | Yes | Yes |
| 750 | 71 | 69.42 | Yes | Yes |
| 1000 | 76 | 68.85 | No | Yes |
| 1500 | 79 | 71.14 | No | Yes |
| 2000 | 81 | 74.85 | No | Yes |
| 3000 | 85 | 76.285 | No | Yes |
| 4000 | 89 | 77.14 | No | No |
| 6000 | 92 | 69.14 | No | No |
| 8000 | 76 | 66.85 | No | Yes |

TABLE LIII. Shows the individual Targets (dB SPL) for Hearing Profile Y ISTS 80 dB SPL for each Frequencies (Hz), and whether the device meets them under two criteria.

| Frequencies (Hz) | Response (dB SPL) | Targets (dB SPL) | Meets Strict Criteria | Meets Loose Criteria |
|---|---|---|---|---|
| 250 | 39 | 56 | No | No |
| 500 | 53 | 57.7 | Yes | Yes |
| 750 | 58 | 54.77 | Yes | Yes |
| 1000 | 56 | 54.35 | Yes | Yes |
| 1500 | 55 | 57.85 | Yes | Yes |
| 2000 | 62 | 61.77 | Yes | Yes |
| 3000 | 67 | 69.96 | Yes | Yes |
| 4000 | 66 | 70.11 | Yes | Yes |
| 6000 | 69 | 62.75 | No | No |
| 8000 | 55 | 58.99 | Yes | Yes |

TABLE LIV. Shows the individual Targets (dB SPL) for Hearing Profile Y G.R.A.S KEMAR Right Ear ISTS 65 dB SPL for each Frequencies (Hz), and whether the device meets them under two criteria.

| Frequencies (Hz) | Response (dB SPL) | Targets (dB SPL) | Meets Strict Criteria | Meets Loose Criteria |
|---|---|---|---|---|
| 250 | 25 | 56.14 | No | No |
| 500 | 48 | 59.14 | No | No |
| 750 | 54 | 58.84 | Yes | Yes |
| 1000 | 55 | 59.14 | Yes | Yes |
| 1500 | 60 | 62.75 | Yes | Yes |
| 2000 | 65 | 66.96 | Yes | Yes |
| 3000 | 67 | 69.96 | Yes | Yes |
| 4000 | 67 | 70.11 | Yes | Yes |
| 6000 | 70 | 62.75 | No | Yes |
| 8000 | 50 | 58.99 | No | Yes |

TABLE LV. Shows the individual Targets (dB SPL) for Hearing Profile Y G.R.A.S KEMAR Left Ear ISTS 65 dB SPL for each Frequencies (Hz), and whether the device meets them under two criteria.

| Frequencies (Hz) | Response (dB SPL) | Targets (dB SPL) | Meets Strict Criteria | Meets Loose Criteria |
|---|---|---|---|---|
| 250 | 41 | 41.46 | Yes | Yes |
| 500 | 46 | 48.44 | Yes | Yes |
| 750 | 48 | 49.08 | Yes | Yes |
| 1000 | 51 | 50.61 | Yes | Yes |
| 1500 | 53 | 58.78 | No | Yes |
| 2000 | 55 | 67.13 | No | No |
| 3000 | 59 | 71.18 | No | No |
| 4000 | 61 | 70.8 | No | Yes |
| 6000 | 69 | 61.91 | No | Yes |
| 8000 | 57 | 57.11 | Yes | Yes |

TABLE LVI. Shows the individual Targets (dB SPL) for Hearing Profile Z ISTS 55 dB SPL for each Frequencies (Hz), and whether the device meets them under two criteria.

| Frequencies (Hz) | Response (dB SPL) | Targets (dB SPL) | Meets Strict Criteria | Meets Loose Criteria |
|---|---|---|---|---|
| 250 | 48 | 56.76 | No | Yes |
| 500 | 61 | 57.91 | Yes | Yes |
| 750 | 62 | 59.08 | Yes | Yes |
| 1000 | 65 | 60.86 | Yes | No |
| 1500 | 66 | 67.74 | Yes | Yes |
| 2000 | 67 | 74.64 | No | Yes |
| 3000 | 71 | 78.82 | No | Yes |
| 4000 | 74 | 69.4 | Yes | Yes |
| 6000 | 81 | 70.83 | No | No |
| 8000 | 67 | 67.598 | Yes | Yes |

TABLE LVII. Shows the individual Targets (dB SPL) for Hearing Profile Z ISTS 65 dB SPL for each Frequencies (Hz), and whether the device meets them under two criteria.

| Frequencies (Hz) | Response (dB SPL) | Targets (dB SPL) | Meets Strict Criteria | Meets Loose Criteria |
|---|---|---|---|---|
| 250 | 54 | 70.86 | No | No |
| 500 | 69 | 72.33 | Yes | Yes |
| 750 | 71 | 69.44 | Yes | Yes |
| 1000 | 76 | 69.7 | No | Yes |
| 1500 | 79 | 74.25 | Yes | Yes |
| 2000 | 81 | 80.53 | Yes | Yes |
| 3000 | 85 | 84.51 | Yes | Yes |
| 4000 | 89 | 84.92 | Yes | Yes |
| 6000 | 92 | 78.15 | No | No |
| 8000 | 76 | 74.68 | Yes | Yes |

TABLE LVIII. Shows the individual Targets (dB SPL) for Hearing Profile Z ISTS 80 dB SPL for each Frequencies (Hz), and whether the device meets them under two criteria.

| Frequencies (Hz) | Response (dB SPL) | Targets (dB SPL) | Meets Strict Criteria | Meets Loose Criteria |
|---|---|---|---|---|
| 250 | 39 | 56.76 | No | No |
| 500 | 53 | 57.91 | Yes | Yes |
| 750 | 58 | 59 | Yes | Yes |
| 1000 | 56 | 60 | Yes | Yes |
| 1500 | 55 | 67.74 | No | No |
| 2000 | 62 | 74.64 | No | No |
| 3000 | 67 | 78.82 | No | No |
| 4000 | 66 | 69.4 | Yes | Yes |
| 6000 | 69 | 70.83 | Yes | Yes |
| 8000 | 55 | 67.598 | No | No |

TABLE LIX. Shows the individual Targets (dB SPL) for Hearing Profile Z G.R.A.S KEMAR Right Ear ISTS 65 dB SPL for each Frequencies (Hz), and whether the device meets them under two criteria.

| Frequencies (Hz) | Response (dB SPL) | Targets (dB SPL) | Meets Strict Criteria | Meets Loose Criteria |
|---|---|---|---|---|
| 250 | 25 | 56.76 | No | No |
| 500 | 48 | 57.91 | No | Yes |
| 750 | 54 | 59 | Yes | Yes |
| 1000 | 55 | 60 | Yes | Yes |
| 1500 | 60 | 67.74 | No | Yes |
| 2000 | 65 | 74.64 | No | Yes |
| 3000 | 67 | 78.82 | No | No |
| 4000 | 67 | 69.4 | Yes | Yes |
| 6000 | 70 | 70.83 | Yes | Yes |
| 8000 | 50 | 67.598 | No | No |

TABLE LX. Shows the individual Targets (dB SPL) for Hearing Profile Z G.R.A.S KEMAR Left Ear ISTS 65 dB SPL for each Frequencies (Hz), and whether the device meets them under two criteria.
